# Boolean modeling of mechanosensitive Epithelial to Mesenchymal Transition and its reversal

**DOI:** 10.1101/2022.08.29.505701

**Authors:** Emmalee Sullivan, Marlayna Harris, Arnav Bhatnagar, Eric Guberman, Ian Zonfa, Erzsébet Ravasz Regan

**Affiliations:** Biochemistry and Molecular Biology, The College of Wooster, Wooster, OH; Ohio University Heritage College of Osteopathic Medicine, Cleveland, OH; Stanford University School of Medicine, Stanford, CA; Molecular, Cellular, and Developmental Biology, UC Santa Barbara, Santa Barbara, CA; Clinical Trial Operations Development, AbbVie, Chicago, IL

## Abstract

The significance of biophysical modulators of the Epithelial to Mesenchymal Transition (EMT) is demonstrated by experiments that document full EMT on stiff, nano-patterned substrates in the absence of biochemical induction. Yet, current models focus on biochemical triggers of EMT without addressing its mechanosensitive nature. Here we built a Boolean model of EMT triggered by mechanosensing – mitogen crosstalk. Our model reproduces epithelial, hybrid E/M and mesenchymal phenotypes, the role of autocrine *TGFβ* signaling in maintaining mesenchymal cells in the absence of external drivers, inhibition of proliferation by *TGFβ*, and its apoptotic effects on soft ECM. We offer testable predictions on the density-dependence of partial EMT, its molecular drivers, and the conflict between mitosis and hybrid E/M stability. Our model opens the door to modeling the effects of the biomechanical environment on cancer cell stemness linked to the hybrid E/M state, as well as the mutually inhibitory crosstalk between EMT and senescence.

## INTRODUCTION

Carcinomas emerge from healthy epithelial tissue due to oncogenic mutations, their growth enabled by the loss of contact inhibition of proliferation ^1,2^. The weakening of cell junctions is critical for tissue invasion and migration leading to metastasis. This transformation from epithelial to an invasive, migratory mesenchymal phenotype is called the Epithelial to Mesenchymal Transition (EMT) ^3,4^ A rate-limiting step in the development of metastatic cancer, EMT remodels the cytoskeleton to enhance migration, disrupts the Extracellular Matrix (ECM) with proteolytic enzymes to promote tissue invasion, and enhances resistance to apoptosis as well as senescence ^5,6^. The reverse mesenchymal-epithelial transition (MET) often aids the settling of these migratory cells into distant metastases ^7^. As metastatic cancers account for 90% of cancer deaths worldwide ^8^, development of effective therapeutic approaches requires predictive models of the complex interaction between contact-mediated signaling, engagement of the EMT transcriptional program, and its reversal during MET.

A closer look at cancer cells within solid tumors revealed a striking cell-to-cell heterogeneity in the expression of epithelial and mesenchymal markers ^9–12^. The initial picture of an all-or-nothing transition from an epithelial to a mesenchymal state gave way to the insight that at least one but perhaps several stable cell phenotypes between these two extremes exist *in vitro* and *in vivo* ^7^. These hybrid E/M phenotypes show characteristics of both epithelial and mesenchymal cells, including an ability to maintain adherens junctions while polarizing horizontally to migrate in a mesenchymal fashion ^13^. As a result, hybrid E/M cells migrate and invade as clusters ^14^, which gives them an advantage in seeding colonies ^15,15–17^, traversing narrow capillary vessels by forming single-cell trains ^18^. Indeed, hybrid E/M cells were shown to have up to 50-fold greater metastatic potential than single circulating cancer cells ^15^. In healthy epithelial tissues such as blood vessels, lung, or liver epithelia, balancing birth and death in response to injury is complemented by partial EMT and migration at monolayer wounds ^3^. Accompanied by proliferation, this helps wounded epithelial sheets migrate as collectives, close wounds, and use MET to reestablish their junctional barriers. Yet, the precise cellular contexts that generate hybrid E/M rather than mesenchymal cells are not well understood ^7^.

EMT is triggered by a series of transforming signals such as *TGF-β, Wnt, Notch, Hedgehog* and *IL6*, aided by growth factors, hypoxia and stiff/fibrous ECM ^19–21^. These influences converge on a core transcriptional circuit rich in positive feedback that can robustly maintain both epithelial and mesenchymal states ^22^. The *Snai1* (*Snail*) transcription factor is a key convergence point; its double-negative feedback with the epithelial micro-RNA *miR-34* is the first switch in the transcriptional cascade of EMT ^23^. Once fully active, *Snai1* lowers the barrier for toggling a second switch generated by mutual inhibition between *miR-200* microRNAs and the transcription factor *Zeb1* by lowering *miR-200* expression while promoting *Zeb1* ^23^. High *Zeb1* and *Snai1* activity fully repress *E-cadherin* transcription, leading to complete disassembly of adherens junctions ^24^. Mesenchymal cells then secrete *TGFβ* to engage an autocrine positive feedback loop that stabilizes their phenotype in the absence of external stimuli ^25,26^. This core regulatory architecture was shown to generate a three-state cascading switch that can maintain a distinct epithelial, hybrid E/M and mesenchymal phenotypes ^27^.

This four-node *Snai1* / *miR-34* / *Zeb1* / *miR-200* circuit is far from capturing the richness of the regulatory links and feedback that modulate EMT. Yet its central role in generating the nonlinearity of the transition allowed computational models detailing their dynamics to reproduce and predict a wide range of experimentally observed behaviors ^22^. For example, the existence of a stable hybrid E/M phenotype was predicted by an ODE-based computational model of this core network ^26,27^ before it was confirmed *in vitro* ^23,28–30^ and *in vivo* ^31–34^. A series of continuous models followed to explore the modulating effect of additional regulators, such as phenotypic stability factors that increase the robustness or number of hybrid E/M states ^35,36^, the relationship between the hybrid E/M state and de-differentiation to a stem-like state ^30^, the effect of *Notch-mediated* spatial patterning in tumors ^37^, or biological noise generating population heterogeneity along the EMT spectrum ^38^. Integrating the crosstalk between signaling pathways controlling EMT and the core transcriptional network was accomplished by a large Boolean network model which linked *TGF-β, Wnt, Notch, Hedgehog* and growth signaling to well-established EMT TFs ^39^. They identified generalized positive feedback motifs (stable motifs) that lock in the mesenchymal state, involving feedback between the core EMT network, major growth pathways, and autocrine *TGF-β/Wnt/Hedgehog* signaling. They went on to characterize perturbations that stabilize hybrid E/M, providing testable interventions to control the network ^40,41^. A subsequent study using roughly the same network but simpler rules that sum positive and negative influences showed that the system can generate hundreds of metastable hybrid states along a spectrum between epithelial and mesenchymal cells, mimicking the heterogeneity of bulk and single-cell gene expression data ^42^.

As Boolean models characterize regulatory signals as expressed/active or absent/inactive ^43^, they rarely deal with processes that depend on the uneven distribution of regulatory molecules in different parts of a single cell. Yet, several key aspects of EMT controlled by the cell’s biophysical environment involve such asymmetry. The transition from apical/basal polarity in an intact monolayer to horizontal polarity and directed migration at lower density requires a different set of regulatory molecules to be ON at the cell’s leading vs. trailing edge ^44^. In a recent study modeling the crosstalk between epithelial contact inhibition of proliferation, migration, cell cycle progression, and anoikis we focused on the mechanosensitive regulation of cell behaviors by ECM stiffness and a cell’s neighbors ^45^. Rather than building a space-aware 3D model, we captured key features of the biology in a Boolean framework by modeling the logic of pathways that push cells past discrete thresholds: anoikis as cells loose ECM anchorage, establishment of apical/basal polarity and contact inhibition in dense monolayers, and the coupling between contact inhibition of proliferation and migration. Our model predicted a metastable migratory state in non-dividing cells at low density but stopped short of integrating these mechanosensing mechanisms with the regulation of EMT-driving transcription factors.

The importance of the biophysical microenvironment in controlling EMT is demonstrated by a series of experiments published EMT model cannot yet reproduce. These include the stiffness-dependent effects of *TGF-β*, leading to apoptosis on soft ECM vs. EMT on stiff substrates ^46^, the demonstration of full EMT in the absence of any biochemical transforming signal on stiff nanopatterned ECMs ^47^, mechanical stress-mediated EMT transcription factor induction ^48–50^, or the effect of cytoskeletal and nuclear morphology on the transcriptional regulation of EMT ^51^. Yet, we lack predictive mechanistic model that can reproduce and predict the effects of a single cell’s biomechanical environment on its behavior in different signaling contexts, such as varying growth stimuli or transforming signals. Moreover, current computational models only address the process of MET in response to either the withdrawal of input signals that activate *Snai1* ^26,27,30^, or to drugs / mutations ^40,41^.

To address these needs, we have built on our published mechanosensitive model of epithelial contact inhibition, proliferation, and apoptosis ^45^ to model the process of EMT triggered by biomechanical and growth signaling crosstalk. Our goal was to examine the stability of hybrid E/M and M states in the absence vs. presence of autocrine *TGFβ* signaling. In addition to EMT regulation (**Fig. 1A-B**), our 150-node Boolean model explicitly accounts for the regulatory networks driving growth signaling, cell cycle commitment and progression, apoptosis, adhesion to the ECM and to neighboring cells, contact inhibition, and key regulators of migration (**Fig. 1C**). We reproduce a range of cell behaviors linked to EMT, including 1) the ability of the core EMT transcriptional network to maintain distinct epithelial, hybrid E/M and mesenchymal states, 2) EMT driven by strong mitogens such as EGF on stiff ECMs ^52,53^, 3) increased mesenchymal transcription factor activity and EMT in response to the loss of *E-cadherin* ^54,55^; 4) TGF*β*-induced EMT and the need for strong autocrine TGF*β* signaling to maintain the mesenchymal state in the absence of mitogens, on softer matrices, or at high density ^25^, 5) TGF*β*-induced inhibition of proliferation ^56^; 6) TGF*β*-induced apoptosis on soft ECM ^46^; 7) anoikis resistance in mesenchymal cells ^57^ driven by TGF*β* ^58^, and 8) lack of TGF*β*-induced EMT in very dense monolayers ^59^. In addition, we offer experimentally testable predictions on the effect of neighbors on partial vs. full EMT, the tug of war between mitosis and the maintenance of migratory hybrid E/M states, the biomechanical triggers of MET and its effect on autocrine TGF*β* signaling, and finally, cell cycle defects in heterogeneous populations of epithelial, hybrid E/M and mesenchymal cells.

**Figure 1.**
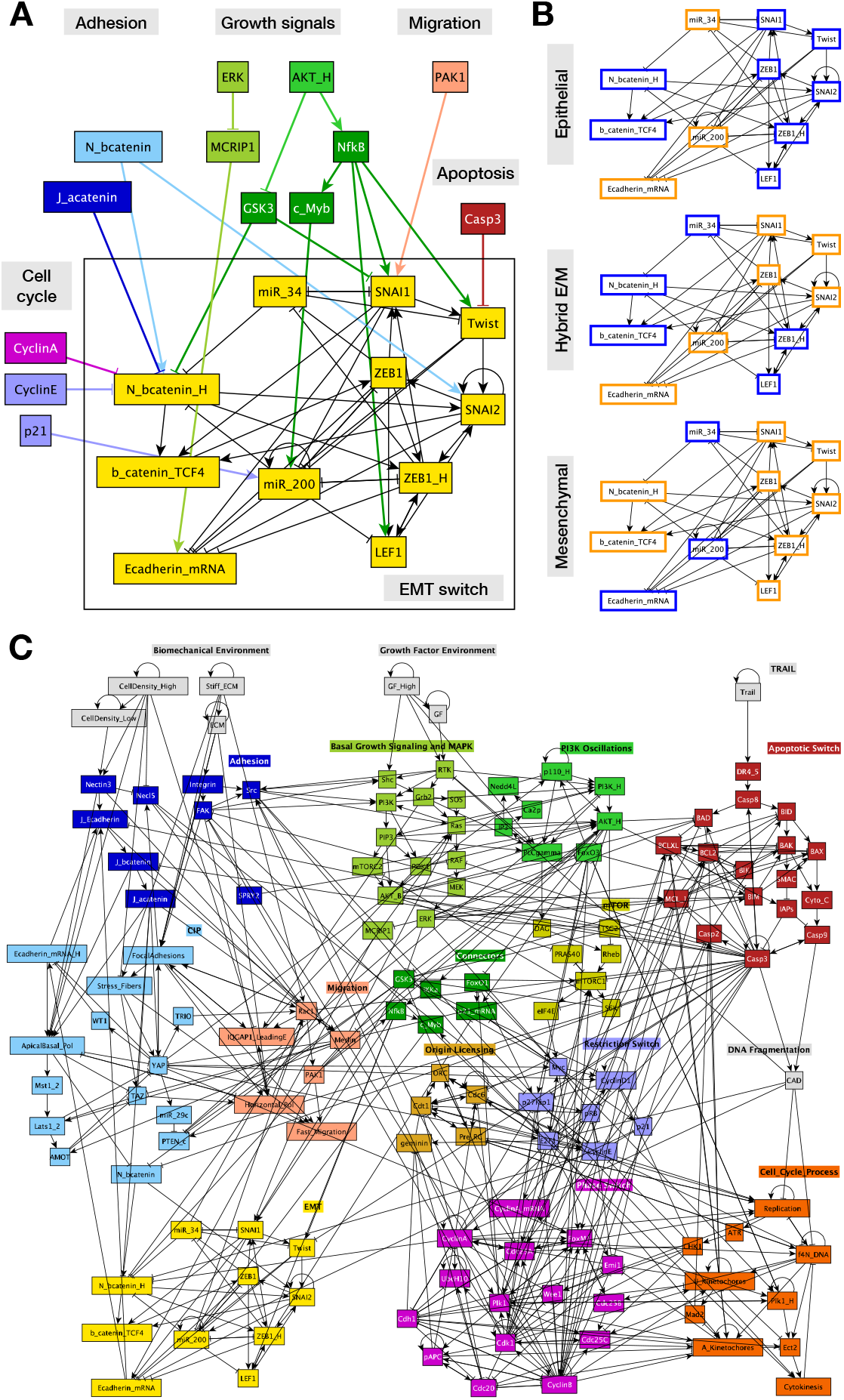
The transcriptional circuit driving EMT is a tree-way switch influenced by cell junctions, migration, and growth signaling. **A)** Regulatory switch controlling EMT (*yellow*) with relevant inputs from regulators of cell cycle (*purple, lavender*), adhesion (*light & dark blue*), growth (*greens*), migration (*pink*) and apoptosis (*dark red;* links from the EMT switch to the rest of the network not shown). **B)** Three stable attractors of the isolated EMT regulatory switch. *Orange/blue node borders:* ON/OFF. **C)** Modular network representation of our extended Boolean model. *Gray:* inputs representing environmental factors; *dark blue:* Adhesion signals; *red: TGFβ* signaling; *green:* Growth Signaling (*lime green:* basal AKT & MAPK, *bright green:* PI3K/AKT oscillations, *mustard:* mTORC1, *dark green: NF-κB, GSK3, FoxO1) dark red:* Apoptotic Switch*; light blue:* Contact Inhibition; *pink/light orange:* Migration; *light brown:* Origin of Replication Licensing; *lilac:* Restriction Switch; *purple:* Phase Switch; *dark orange:* cell cycle processes;*yellow:* EMT switch; → : activation; ⊣ : inhibition.

## RESULTS

### Modeling the epithelial to mesenchymal toggle switch

To model the effect of mechanosensitive signals on EMT, we added an EMT switch to our contact inhibition model ^45^ (**Fig. 1A**). We built this switch around feedback loops common to published continuous and Boolean models ^26,27,39^. These include direct double-negative feedback between *Snai1* and the epithelial microRNA *miR-34* ^60^ as well as between *Zeb1* and *miR-200* ^61^. They are reinforced by positive feedback between *Snai1*, *Zeb1*, nuclear *β-catenin* and the two miRs ^62–65^. This switch controls *E-cadherin* mRNA expression, repressed cooperatively by mesenchymal transcription factors ^66–68^. As *Zeb1* is not expressed in epithelial cells, expressed at a moderate level in hybrid E/M states, and is only high in mesenchymal cells ^69,70^, we used two Boolean nodes to distinguish between these three levels of *Zeb1.* Similarly, we modeled the effects of adherens junctions on *β-catenin* nuclear localization with two nodes, distinguishing between a) cells entirely surrounded by neighbors in a dense monolayer (all *β-catenin* is localized to adherens junctions; both nodes OFF), b) cells firmly attached to the edge of a monolayer (some but not all *β-catenin* is tied to adherens junctions; medium nuclear localization, *N_bcatenin=ON* but *N_bcatenin_H=OFF*), and c) cells that do not touch their neighbors (all *β-catenin* is freed from junctions even if *E-cadherin* is expressed; nuclear localization is maximal as long as *GSK3β* is blocked by growth signals; both nodes ON). **Fig. 1A** highlights key signals that converge on the EMT module from cell junctions (*blue*), growth signaling (*green*), and the migration prompting *Pak1* kinase (*pink*).

Isolated from the rest of the network, the EMT switch has three stable phenotypes (**Fig. 1B**; automated module isolation and synchronous attractor detection from ^46^; see *STAR Methods*). As expected from a switch constructed to mirror the known two-stage progression of EMT, its epithelial stable state is characterized by *E-cadherin*, *miR-34* and *miR-200* expression, and no transcriptional activity in EMT driver genes (**Fig. 1B**, *top*). The opposite pattern of expression and activity characterizes the mesenchymal state, a complete flip of the switch (**Fig. 1B**, *bottom*). In between the two, the network maintains a stable hybrid E/M state characterized by fully suppressed *miR-34* but not *miR-200* ^27^, along with *Snai1, Twist, Snai2* and moderate activation of both *Zeb1* and nuclear *β-catenin* (**Fig. 1B**, *middle*).

The regulatory environment outside the switch can bias or dictate the state(s) it can maintain in different environments. To model this we linked the EMT switch into our contact inhibition model ^45^, where it is influenced primarily by *i*) mechanosensing and junction-mediated contact inhibition via *β-catenin* localization, *ii*) migration via *Pak1*, and *iii*) growth signaling via *Akt1-* mediated *NF-κB* activation and *GSK3-β* suppression (**Fig. 1A,C**).

### Growth factors cooperate with matrix stiffness and loss of cell-cell contacts to drive reversible mechanosensitive EMT

The key difference between the EMT model on **Fig. 1C** and previous models is that it does not include the traditional exogenous drivers of EMT such as *TGF-β, Wnt, Notch, Sonic Hedgehog*, or *IL-6* ^39,71^. Instead, we focus on signals that can trigger EMT in the *absence* of the above signals – namely the cell’s biomechanical environment. Evidence from different cell lines indicates that epithelial cells undergo an epithelial to mesenchymal transition when plated on stiff substrates ^47–50^. To model this, we started by scanning all stable phenotypes of our model on a soft ECM at different cell densities and growth factor exposures (sampling algorithm that maps the model’s stable phenotypes / attractors detailed in *STAR Methods*). Due to an inability to form stress fibers and activate *YAP* on soft ECM, the only stable model phenotype is epithelial – even at very low densities that preclude adherens junctions (**Fig. 2A,** *initial state*). Plating our model epithelial cells on a stiff matrix triggered stress fiber formation, *YAP* activation and *Rac1/Pak1* -driven migration (**Fig. 2A**, transition in *Migration* module at the start of *Stiff ECM* exposure). In addition, *YAP* led to the induction of high *PI3K 110* (*p110*) protein expression ^72^. In growth-promoting environments (serum, EGF, HGF), high expression of *p110* increases the potency of *PI3K* / *AKT* signaling, which activates *NF-κB* and turns off *GSK3β* ^73^. *NF-κB*, in turn, lead to *Snai1* transcription, stabilized by low *GSK3β* activity ^74,75^. *Snai1* nuclear localization is subsequently aided by *Pak1*-mediated phosphorylation, a key early event leading to EMT ^76^. Thus, *YAP*-mediated sensing of a stiff ECM works cooperatively with growth signaling to promote *Snai1* activation and subsequent *miR-34* repression by *Snai1.* Without *miR-34* to lower *β-catenin* and neighbors to trap it at adherens junctions, nuclear *β-catenin* levels rise sufficiently to support high *Zeb1* activation, *miR-200* repression, and full EMT (**Fig. 2A**, *final state*). This effect is followed by cell cycle entry, which does not require EMT but is also supported by growth signals and a stiff ECM (**Fig. 2A**, *cell cycle oscillations*). In contrast, plating cells on a stiff ECM in the *absence* of strong growth signaling does not result in high *AKT* signaling, migration, *Snai1* activation, or EMT (**Fig. S1A**). An alternate route to the same mesenchymal phenotype is to start with isolated cells plated on a stiff ECM in the absence of strong growth signaling. Exposing these cells to growth factors also triggers EMT via the same cascade of signals (**Fig. 2B**).

**Figure 2.**
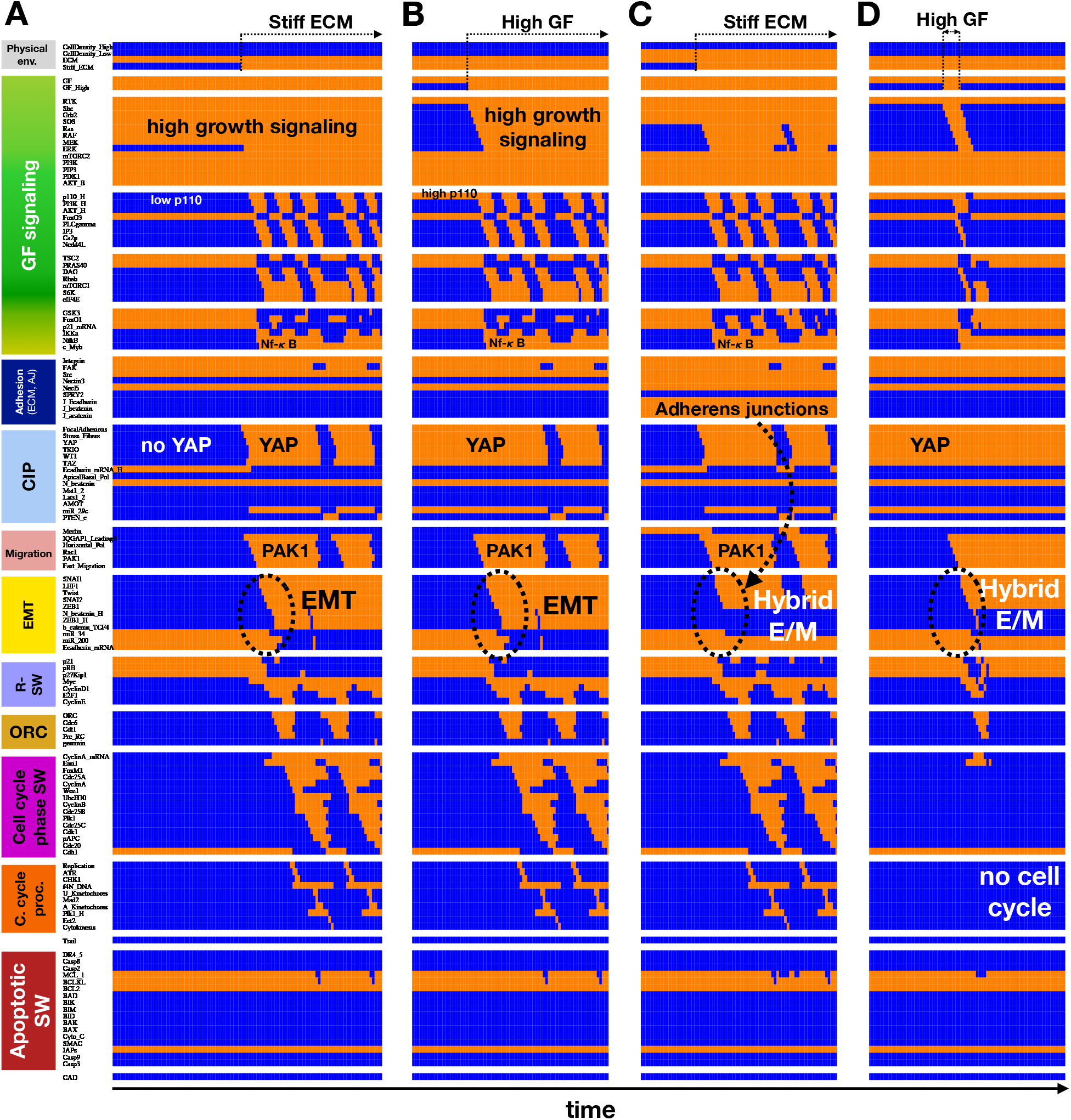
Strong growth signals on a stiff matrix can drive the epithelial to mesenchymal transition, leading to mesenchymal cells at very low density vs. a hybrid E/M state at monolayer edges. **A-D)** Dynamics of regulatory molecule expression/activity during exposure of (A) isolated, growth-stimulated epithelial cells to stiff ECM, (B) isolated epithelial cells in low growth conditions on a stiff ECM to strong mitogens, (C) isolated, growth-stimulated epithelial cells at a monolayer edge to stiff ECM, and (D) isolated epithelial cells in low growth conditions on a stiff ECM to a brief (6-step) pulse of strong mitogens. *X-axis:* time-steps; *y-axis:* nodes organized in regulatory modules; *orange/blue:* ON/OFF; *black/white labels & arrows:* molecular changes that drive EMT.

### Mechanosensitive EMT dynamics is robust to fluctuations in signal propagation, changes in Boolean update, and random errors in model construction

The Boolean nature of the model’s environmental inputs requires the assumption that the ON state of an input node such as *GF_High* activated its downstream signaling pathway to saturation. To model *intermediate* levels of growth signal exposure, we chose to stochastically toggle *GF_High* ON for a tunable fraction of time steps, which renders the growth signal stochastic. We assume that such intermediate signals correspond to an *in vitro* concentration range at which the targets of growth signals are in transition from non-responding to saturating (e.g., near the inflection point of an S-shaped response curve), a regime in which the response is most susceptible to biological noise ^77,78^. Our stochastic growth input’s downstream effects mimic this probabilistic, intermittent response ^79^. Similarly, medium (non-saturating) ECM stiffness modeled via a stochastic ON/OFF toggle mimics a cell that tugs on the ECM with imperfect, noisy reliability with respect to downstream *Rac1* activation and stress fiber generation. The comparable length scales of stress fibers and collagen fibers (or other ECM components), together with their inhomogeneous spatial organization, further supports a model with a noisy response to moderately stiff ECM. In our model, increasing the ECM stiffness under cells exposed to near-saturating mitogens (*GF_High* = 90%) shows a rapid increase in proliferation above ~50% *Stiff_ECM* exposure (**Fig. S1B,** *top left*), and vice versa (**Fig. S1B,** *bottom left*). **Fig. S1B** also shows that cells on stiffer matrices spend increasing amounts of time in a mesenchymal state (*top row, orange*). The response to growth signals is similar, except that we predict a high prevalence of hybrid E/M cells at intermediate mitogen exposure (**Fig. S1B,** *bottom row, orange*).

The results in figures **2** and **S1** were generated using synchronous update, where all Boolean nodes change state in each time-step at the same time. To ensure that the underlying biology of the network (its nodes, links, and regulatory logic) rather than the built-in deterministic dynamics of synchronous update, is responsible for the observed behavior, we next turned to random-order asynchronous update (sequential update in changing random order). A Boolean model’s stable states do not depend on the update scheme, but oscillations such as the cell cycle may be derailed by non-biological update orders generated by asynchronous update. We have previously shown that asynchronous update applied to our cell cycle model generates a mix of behaviors from normal cycles to errors that mimic defects in mitosis (including apoptosis due to mitotic catastrophe) ^80^. As fully randomized update enriches for errors, we introduced a biased update where a small subset of nodes is updated in order at the start or end of list (see *STAR Methods*). We used this method to rerun Figure 2 simulations, averaging over a large set of independent dynamical trajectories akin to a plate of noisy cells (**Fig. S2**). Despite the rapid desynchronization of cell cycle and *PI3K/Akt* oscillations across these runs, EMT remains robust to the change in update. Moreover, figures **S1A-B** and **S3A-B** portraying the same *in silico* experiments with different update, are qualitatively similar (note the small fraction of genome-duplication errors introduced by asynchronous update on **Fig. S3B**, *top left*).

Next, we tested the model’s robustness to minor structural errors by generating three distinct ensembles of random mutants (**Fig. S4**). First, we locked an increasing number of random nodes ON or OFF and averaged the dynamical behavior of the resulting mutant cells. Comparing figure **S4A** and **S4B** shows that both cell cycle progression and mechanosensitive EMT are robust to 2 random knockout/knock-in errors, then struggle to maintain the mesenchymal state with 5 errors per model. For the second ensemble, we removed an increasing number of randomly chosen links (**Fig. S4C**) showing that the network’s dynamics is robust to ~5 randomly missing links. Our third ensemble with random errors in gate output show a similar degree or robustness, indicating that missing links have an equivalent effect on the dynamics as flipping the expected output of a node for one combination of inputs. Overall, the ensemble’s dynamical behaviors break in a modular fashion with increasing damage, starting with a loss of cell cycle synchrony followed by stabilization of hybrid E/M, then the epithelial state.

### Modeling the emergence of hybrid E/M states: cells at the edge of a monolayer undergo partial EMT in response to stiff ECM exposure

Repeating the above simulations with cells placed in cell densities akin to the edge of a confluent monolayer leads to a different outcome. As **Fig. 2C** indicates, exposing cells that maintain adherens junctions with some neighbors but not enough to establish apical/basal polarity respond to stiff ECM exposure by undergoing partial EMT. In this case, *Snai1* activation and *miR-34* inhibition do not result in high nuclear *β-catenin.* Instead, cells stabilize in a state that matches the observed transcription factor expression and phenotype of hybrid E/M. Namely, *Snai1, Twist1* and *Snai2* are expressed, *Zeb1* activity is moderate, and *miR-34* is repressed but *miR-200* is still active (**Fig. 2C**, EMT module). These cells are migratory and have reduced *E-cadherin* expression compared to dense monolayers. Yet, they maintain adherens junctions with neighbors and migrate as a group (**Fig. 2C**, *Migration* and *CIP* modules). These behaviors are robust under biased asynchronous update (**Fig. S2C**), as well as in the presence of random errors in model structure (**Fig. S4**, *right column*) Modeling intermediate levels of stiffness also shows that cells with adherens junctions spend an increasing amount of time in a hybrid E/M on stiffer substrates, but do not undergo full EMT (**Fig. S1C/S3C**).

Interestingly, we predict that while a sufficiently stiff ECM to sustain stress fiber formation is mandatory for entry into and maintenance of a hybrid E/M phenotype, sustained *growth* signaling is not necessary. Rather, entry into hybrid E/M only requires a short-lived boost of growth signaling (**Fig. 2D**). Once established, positive feedback linking the *EMT* and *Migration* modules can sustain hybrid E/M. Consequently, even weak / transient growth factor exposure has a large effect on the amount of time these cells spend in a hybrid E/M state (**Fig. S1C/S3C**, *bottom right*, synchronous/asynchronous update). Saturating growth signals actually *decrease* the stability of the hybrid E/M state under synchronous update (**Fig. S1C**), though the effect is blunted by asynchronous update (**Fig. S3C**).

To further probe the importance of adherens junctions in partial vs. full EMT, we ran a large sample of cells stochastically exposed to a matrix with relatively high stiffness (*Stiff ECM* = ON in 95% of time-steps) and near-saturating growth factors (95% *GF_High_* = ON). In this environment, a randomly sampled cell population will be ~ 40% epithelial (**Fig. S1D/S3D,** *leftmost blue bars*, synchronous/asynchronous update), ~55% in hybrid E/M (*leftmost orange bar*), and 0% mesenchymal (the remaining 5% are in transition). Increasingly potent inhibition of junctional *E-cadherin* leads to a moderate decrease in epithelial states, but a drastic shift from hybrid E/M to full mesenchymal cells (**Fig. S1D/S3D**). This simulation, akin to either experimental inhibition of *E-cadherin* or gradual decrease in density, underscores the importance of cell contacts in stabilizing the hybrid E/M state even in the absence of *Notch*-mediated juxtracrine signaling ^37^.

### Modeling the Mesenchymal to Epithelial Transition: in the absence of strong autocrine signaling, loss of YAP or ERK activity reverses EMT

As our model lacks autocrine signals that stabilize the mesenchymal or hybrid E/M state, their stability relies on ongoing feedback between the EMT switch, growth signaling, and migration. To investigate the consequences of breaking this feedback, we simulated the response of isolated mesenchymal cells proliferating on a stiff matrix to two types of environmental change. First, we simulated their re-plating onto very soft ECM, where the maintenance of stress fibers, *YAP* activity, horizontal cell polarity and migration is no longer possible (**Fig. 3A/S5A**, synchronous/asynchronous update). This led to a loss of *Pak1* signaling, nuclear exclusion of *Snai1*, and MET. Indeed, even partial matrix softening was sufficient to lock cells into an epithelial state, as mesenchymal entry below 50% *Stiff_ECM* is rare (**Fig. S1B/S3B,** *top rows*).

**Figure 3.**
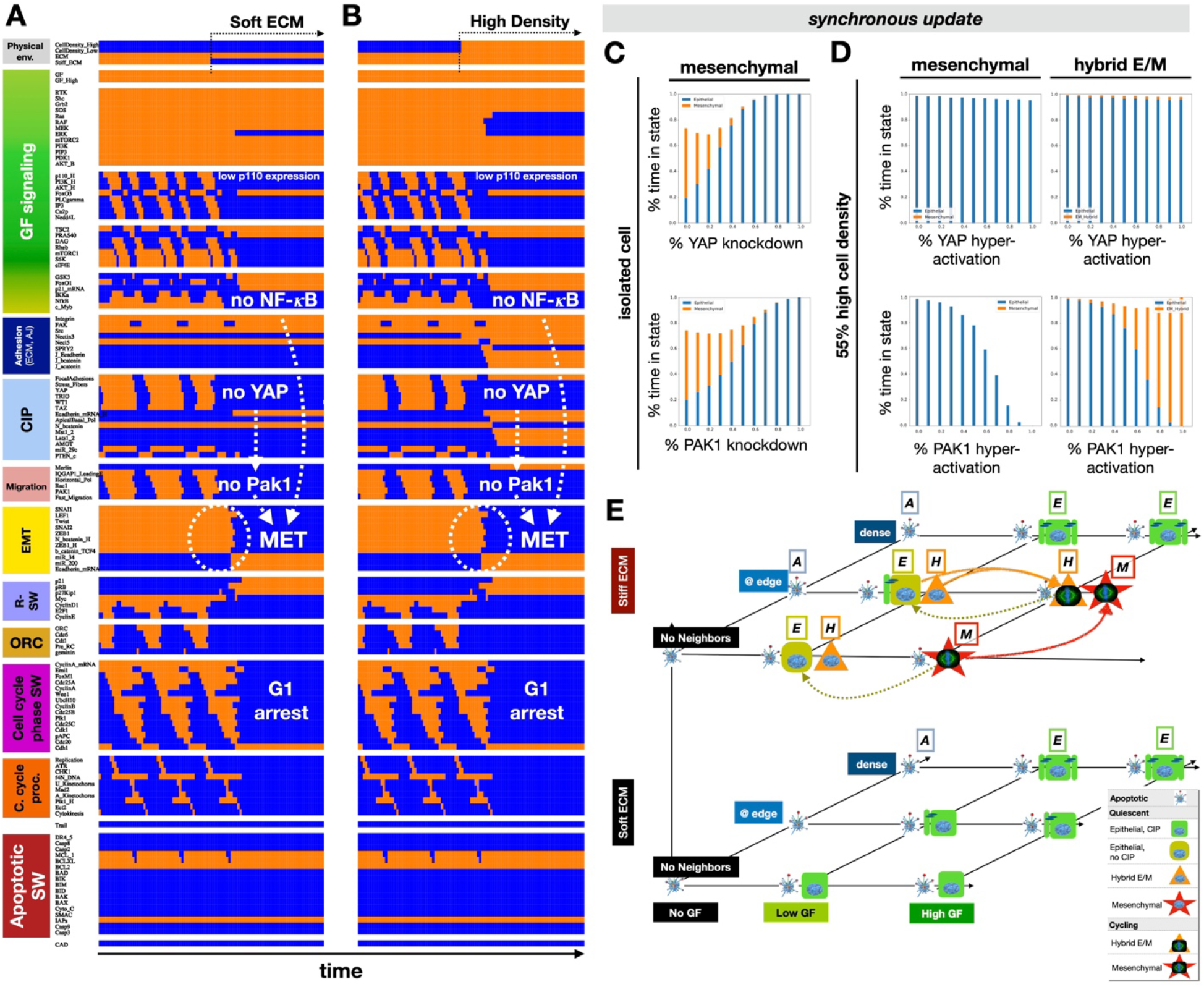
The mesenchymal to epithelial transition (MET) is triggered by both soft matrix exposure and high cell density in the absence of autocrine or external EMT-promoting signals. **A-B)** Dynamics of regulatory molecule expression/activity during exposure of an isolated, growth-stimulated and proliferating mesenchymal cell to (A) soft ECM, and (B) very high cell density. *X-axis:* time-steps; *y-axis:* nodes organized in regulatory modules; *orange/blue:* ON/OFF; *black/white labels & arrows:* molecular changes that drive MET. **C)** *Top/bottom:* fraction of time spent in a mesenchymal (*orange*) vs. epithelial (*blue*) state as a function of increasing *YAP1 (top*) or *PAK1 (bottom*) inhibition in isolated cells on 95% *Stiff ECM* and exposed to 95% saturating growth stimuli. **D)** *Top/bottom:* fraction of time spent in *i*) mesenchymal (*orange*) vs. epithelial (*blue*) state (*left*) or *ii*) hybrid E/M (*orange*) vs. epithelial (*blue*) state (*right*) as a function of increasing *YAP1 (top*) or *PAK1 (bottom*) inhibition in cells at 55% high density (no free space to spread 55% of the time) on 95% *Stiff ECM* and exposed to 95% saturating growth stimuli. *Total sampled live cell time:* 100,000 steps; synchronous update; *Initial state for sampling runs:* isolated epithelial cell in low mitogens on a soft ECM. **E)** Summary of model cell states predicted for every combination of no/low/high growth-factor (*x axis*), no/low/high cell density environments (*y axis*), and very soft ECM (<0.5 kPa) / very stiff ECM (*z axis). Blue fragmented cell & “A” label:* apoptotic state; *green box & “E” label:* quiescent epithelial cell; *orange triangle & “H” label:* hybrid E/M cell, *red star & “M” label:* mesenchymal cell; *symbol with blue nucleus:* non-dividing cell; *symbol with mitotic spindle:* proliferating cell. Cells suspended without ECM are all apoptotic (not shown); each quiescent state has a counterpart with double DNA content (not shown). Image credits: apoptotic cell ^118^; mitotic spindle: *https://en.wikipedia.org/wiki/Cell_division#/media/File:Kinetochore.jpg.*

Placing mesenchymal cells into a high cell density environment has a similar effect to soft ECM, as it also triggers MET (**Fig. 3B/S5B**). This, however, only occurs at densities that forbid cell spreading and stress fiber maintenance. Placing a mesenchymal cell into a neighborhood that resembles the edge of a monolayer does not alter its mesenchymal state (**Fig. SA**). The presence of neighbors alone fails to trigger partial MET to a hybrid E/M state because fully mesenchymal cells lack *E-cadherin* expression and do not form adherens junctions. Thus, MET only occurs in our model when feedback between the EMT switch and migration is interrupted, and *Snai1* / *Zeb1* activity is lost. Only then do cells re-express *E-cadherin* and establish adherens junctions.

To highlight the key role of *YAP* and *Pak1* in the maintenance of the mesenchymal state, we modeled partial *YAP* or *Pak1* knockdown in a 95% *Stiff_ECM*, 95% *GFHigh* environment. To do this, we locked *YAP* or *Pak1* OFF for a random fraction of time-steps (tunable from 0% to 100%) and allowed them to respond to their inputs the rest of the time. The rationale for such partial knockdown is that it offers a way to model the incomplete inhibition seen in cells treated with a chemical inhibitor or siRNA silencing, as opposed a genetic knockout. As **Fig. 3C** and **S5C** indicate, partial downregulation of both *YAP* or *Pak1* can trigger MET, despite the lack of neighbors and a stiff ECM. In contrast, forced activation of *Pak1* can partially counteract MET at relatively high density by stabilizing hybrid E/M, while hyperactive YAP alone cannot (**Fig. 3D/S5D**, synchronous/asynchronous update).

The fact that even transient interruption of migration can block *Snai1* and flip the EMT switch leads to an unexpected model prediction for proliferating mesenchymal cells. Namely, during mitosis *Cdk1/Cyclin B* trigger a temporary loss of *Pak1* as cells loosen their contact with the ECM, loose their stress fibers and round up to assemble the mitotic spindle (**Fig. 2** *Migration* module in mesenchymal cells) ^81^. The resulting *Pak1* inactivation does not impact the EMT switch directly, as both *miR-34* and *miR-200* remain repressed by *Zeb1* and *Snai1*. That said, in cells that complete mitosis in the *absence* of strong growth signals, loss of *NF-κB* and reactivation of *GSK3β* compounds the loss of *Pak1*, all tipping the scales towards the loss of active *Snai1.* Thus, our model predicts that the loss of mitogenic stimuli leading to cell cycle exit is accompanied by MET (**Fig. S6B/S6D**). A similar fate awaits hybrid E/M cells exiting the cell cycle (**Fig. S6C**). In contrast, cells that undergo partial EMT with the aid of a *short-lived* growth signal which does not push them into the cell cycle remain migratory and maintain a hybrid E/M state in low growth factor environments (**Fig. 2D/S2D**). This behavior explains the biphasic response to increasing growth factors observed with synchronous update (i.e., under simulation conditions where the response to a fixed level of growth stimulus is less noisy). Namely, moderate growth signals appear to increase, but saturating signals *decrease* the stability of the hybrid E/M state (**Fig. S1C** vs. **S3C**; effect blunted by asynchronous update).

### The full phenotype repertoire of our model showcases the mechanosensitive routes to EMT and MET

To summarize the model’s ability to reproduce the cell phenotypes we expect to see in different microenvironments, we visualized our model’s phenotypes (attractors) across all input-combinations (**Fig. 3E**; for sampling algorithm and complete list of stable phenotypes across all environments see *STAR Methods* and **Table S1**). To this end we created a coordinate system in which each axis represents an independent environmental factor ^45,80,82^. Omitting all *Trail* = ON and *ECM* = OFF conditions where all cells eventually die allowed us to reduce the dimensionality of the environment space to three independent inputs: growth signals (*x* axis), cell density (*y* axis) and ECM stiffness (*z* axis). Distinct positions (coordinates) along the resulting 3D grid represent unique environment-combinations. At each unique coordinate we placed a small picture to represent each stable cell phenotype. For example, apoptosis (*blue apoptotic cell*) is the only stable state in environments with no growth factors – the leftmost position along the *x* axis representing growth signals. In environments that also support non-apoptotic attractors, there is more than one phenotype-symbol (e.g., apoptotic cells and green epithelial cells in growth factors with soft ECM; **Fig. 3E** *bottom plane*).

As **Fig. 3E** indicates, a mesenchymal phenotype in the absence of external or autocrine drivers of EMT is only stable on stiff ECM, at medium to low cell density, in the presence of strong mitogens (*red stars*). Weaker, non-proliferative growth stimuli still permit the maintenance of hybrid E/M, provided they can stretch on stiff ECM (moderate / low density, *orange triangles*). If they become isolated when exposed to strong growth signals, these hybrid E/M cells undergo full EMT to a mesenchymal state (*lone red star*). In contrast, both soft ECM and high cell density force cells into an epithelial state (*green boxes*), while loss of all growth signals leads to apoptosis (*light blue images*).

Our model makes intriguing predictions about the bi-stable mix of epithelial and hybrid E/M cells that coexist under low growth signaling. Namely, we predict that short spikes of growth signaling that fail to trigger cell cycle entry can push epithelial cells into hybrid E/M at a monolayer’s edge. This hybrid E/M state is maintained, along with a migratory state, until interrupted by mitosis, soft ECM exposure, or high cell density. In contrast, transitions triggered by *sustained* growth signals trigger cell cycle entry and lead to reversible EMT (**Fig. 3E,** EMT: *red/orange arrows;* MET: *mustard arrows*).

### Autocrine TGF-β signaling locks in EMT triggered by growth signaling on stiff ECM and blocks mesenchymal cells proliferation

Next, we set out to model the crosstalk between the biomechanical environment and autocrine *TGFβ* signaling required to sustain the mesenchymal state *in vitro* ^25,26^. To this end we added a *TGFβ* signaling cascade, as well as a secreted *TGFβ* node to the EMT switch (**Figs. 4A, S7**). In this new module, externally applied or secreted *TGFβ* can bind to *TGFβ* receptors I and II, which in turn activate *Smad 2/3/4* ^83^. *Smads* drive EMT by promoting nuclear translocation of *β-catenin*, and inducing *Snai2, Lef1*, and *HMGA2* expression*. HMGA2* further induces *Snai1, Snai2*, and *Twist.* In addition, we included *TGFβ’s* paradoxical effect on the MAPK cascade along with PI3K/AKT activation, anti-proliferative *p15* induction, and pro-apoptotic *BIM/BIK* induction and *BCL-XL* repression (**Fig. 4A**; see **File S1** for details). To create autocrine positive feedback, we included *TGFβ* secretion downstream of nuclear *β-catenin/TCF4* (repressed by *miR-200*).

**Figure 4.**
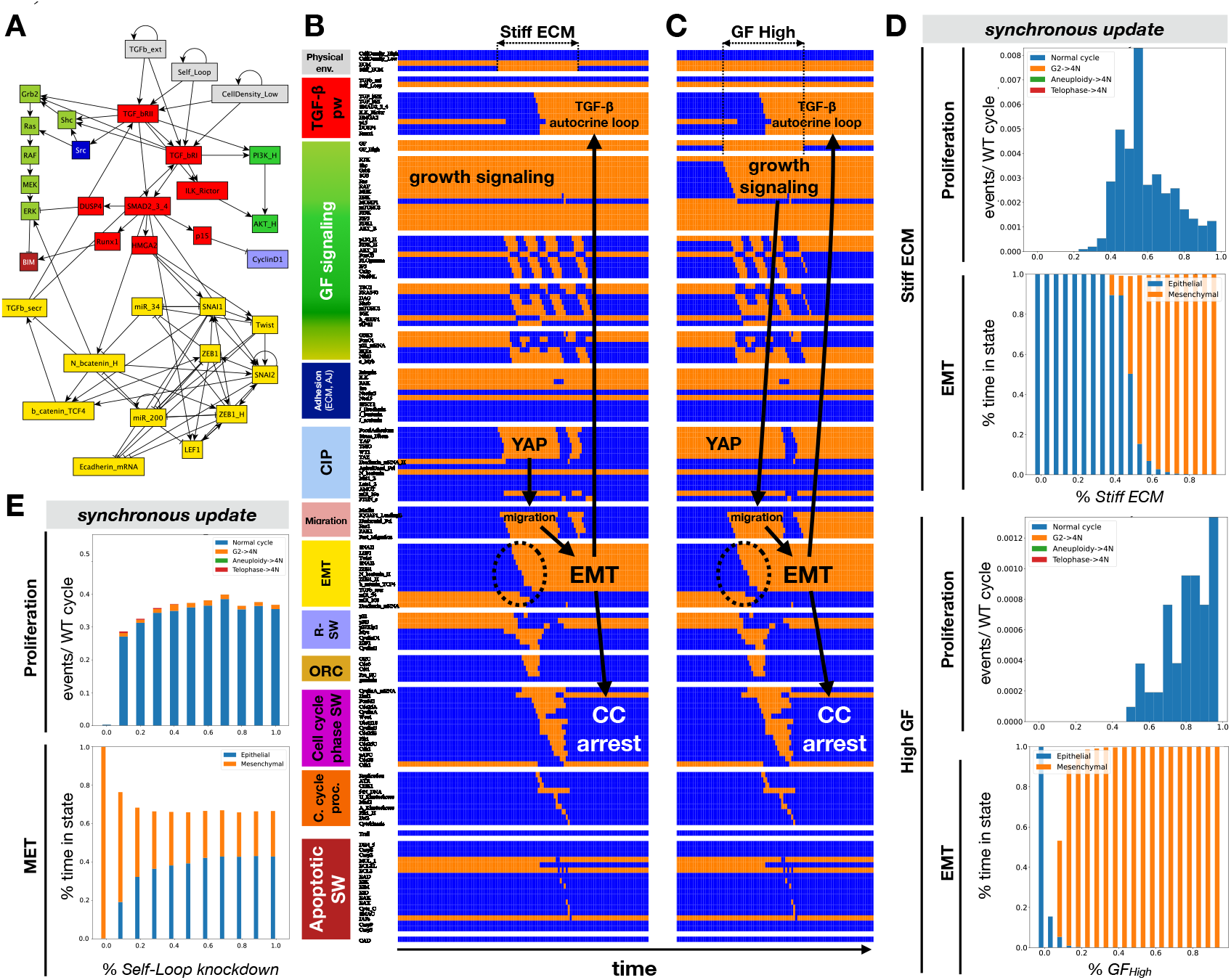
At very low density, stiff ECM and growth signaling trigger stable commitment to a mesenchymal state, sustained by strong autocrine *TGFβ* signaling. **A)** *TGFβ* signaling module (*red*) linked to the previous model’s EMT switch (*yellow;* augmented with a node for secreted *TGFβ*), growth signaling (*light green:* MAPK; *dark green:* AKT) supported by *Src* activation (*dark blue*), cell cycle entry (*lilac*) and apoptosis (*dark red*). **B, C)** Dynamics of regulatory molecule expression/activity during exposure of (B) an isolated, growth-stimulated epithelial cell to stiff ECM, and (C) an isolated epithelial cell on stiff ECM to strong mitogens. *X-axis:* time-steps; *y-axis:* nodes organized in regulatory modules; *orange/blue:* ON/OFF; *black/white labels & arrows:* molecular changes that drive EMT. **D)** *Top two vs. bottom 2 panels:* response of isolated cells to *i*) increasing *Stiff ECM* exposure in the presence of 95% saturating growth stimuli (*top two*) vs. ii) increasing growth factor exposure on 95% stiff ECM (*bottom two). Proliferation label:* rate of normal cell cycle completion (*blue*) vs. G2 → G1 reset (*orange*, not observed), aberrant mitosis (*green*, not observed), or failed cytokinesis (*red*, not observed) followed by genome duplication, relative to the minimum cell cycle length (21 time-steps), shown as stacked bar charts. *EMT label:* fraction of time spent in a mesenchymal (*orange*) vs. epithelial (*blue*) state. **E)** Response of isolated cells on 85% Stiff ECM exposed to 85% saturating growth stimuli to increasing inhibition of autocrine signaling (*Self-Loop* knockdown). *Top:* rate of normal cell cycle completion (*blue*) vs. G2 → G1 reset (*orange*), aberrant mitosis (*green*, not observed), or failed cytokinesis (*red*, not observed) followed by genome duplication. *Bottom:* fraction of time spent in a mesenchymal (*orange*) vs. epithelial (*blue*) state. *Total sampled live cell time:* 100,000 steps; synchronous update; *Initial state for sampling runs:* isolated epithelial cell in low mitogens on a soft ECM.

Adding an autocrine loop to a Boolean regulatory model makes the often-unexamined assumption that a single cell is capable of secreting sufficient signal (in our case *TGF-β*) to drive a *saturating* response (i.e., pathway is ON rather than OFF). As a result, the effects of autocrine loops can be easily overestimated in Boolean models, indicating irreversible lock-in of EMT in situations where cells *in vitro* show metastable EMT due to partial autocrine activation ^84^. To mitigate this, we included an additional input named *Self_Loop.* When set to 0, *TGFβ* secreted by an isolated cell is too weak to trigger signaling, whereas *Self_Loop=ON* represents saturating levels of autocrine *TGFβ* activation. Tuning this input to intermediate values allows us to model a range of autocrine signal strengths in sparsely plated cells. To further account for the possibility that cells with mesenchymal *TGFβ-secreting* neighbors have stronger autocrine signaling ^85^, our model allows secreted *TGFβ* to saturate signaling at cell densities of a monolayer’s edge or above regardless of the *Self_Loop* setting.

To test the effects of a potent autocrine loop, we re-ran the mechanosensitive experiments shown on **Fig. 2** with the extended model. Namely, we exposed isolated epithelial cells to stiff ECM and saturating growth factors (**Figs. 4B,C/S8A,B**, synchronous vs. asynchronous update). Cells exposed to growth factors on soft ECM remain epithelial (**Fig. 4B/S8A***, left*) but undergo EMT when plated on a stiff matrix for a sufficient interval (*middle*) and sustain their mesenchymal state when replated on soft ECM (*rigth*). As they fully engage autocrine *TGFβ* signaling, they also cease to proliferate (**Fig. 4B/S8A**). Indeed, autocrine *TGFβ* is responsible for non-monotone proliferation response with increasing matrix stiffness, where maximum proliferation is predicted at the stiffness boundary at which EMT is triggered (**Fig. 4D/S8C,** *top 2 panels*; ~55% / ~ 35% *Stiff_ECM for* synchronous / asynchronous update). Below this, soft ECM limit proliferation; above the threshold it is blocked by *TGF-β.* A similarly irreversible EMT is observed in cells plated on stiff matrix first, then exposed to high growth factors (**Fig. 4C/S8B**), except for the appearance of hybrid E/M cells in response to weak growth stimuli too weak to trigger proliferation (**Fig. 4D/S8C,** *bottom 2 panels*). This hybrid E/M is triggered by short bursts of growth factors that do not persist long enough to trigger cell cycle or lock in the autocrine signaling loop (**Fig. S11A/S12A** vs. **S11B/S12B**). In contrast, short-lived exposure to a stiff ECM either fails to trigger sustained EMT (**Fig. S11C/S12C**) or leads to *TGF-β*-mediated apoptosis (**Fig. S11D/S12D**) (see next section).

Next, we weakened autocrine *TGF-β* from 1 to 0 in isolated cells exposed to an environment that favors but cannot fully force mechanosensitive EMT; namely 85% growth signals and 85% Stiff ECM. As **Fig. 4E** and **S8E** indicate, even a slight weakening of the autocrine loop decreases the time cells spend in a mesenchymal state, restoring proliferation instead. Its full knockout leads to a mix with majority epithelial vs. mesenchymal cells (*bottom panel*). In summary, our model shows that growth signals and stiff ECM work together to drive EMT in sparsely plated cells, a transition that is subsequently maintained by strong autocrine *TGFβ* signaling.

In contrast to full EMT in sparse cells, partial EMT at the edge of a monolayer does not result in *TGFβ* secretion in our model. As a result, the extended model’s behavior is the same as that of the previous one (**Figs. 2C-D** vs. **S13-S14**). Namely, matrix stiffness and growth signaling induce reversible hybrid E/M and proliferation in cells at the edge of a monolayer (**Figs. S13A,B**), but sustained migratory hybrid E/M in cells exposed to sub-proliferative bursts of growth signaling (**Fig. S13C**). We predict that moderate levels of growth factor exposure on stiff ECMs leads to the most time spent in a hybrid E/M state (**Fig. S14B,** *bottom*), as cells reset from a hybrid E/M to an epithelial state every time they round up for mitosis. While short bursts are necessary for partial EMT, sustained exposure triggers a conflict between proliferation and migration (hybrid E/M) and requires partial EMT after each division. Knockdown of *E-cadherin* breaks the tug of war by favoring full EMT at the expense of proliferation (**Fig. S14C**).

Next, we set out to text whether altering the biomechanical environment can trigger MET in the presence of strong autocrine *TGFβ* signaling. After verifying that mesenchymal cells remained stable at high cell density (**Fig. 5A/S15A**, *left*), we exposed them to soft ECM. Loss of both space and stiff ECM triggered MET by activating *Sprouty2*, which in turn repressed both *Smad* activity and MAPK signaling. Mesenchymal cells plated on soft ECM and then crowded to high density underwent a similar MET (**Fig. 5B/S15B**). At intermediate densities that only trigger intermittent *Sprouty2* activity, our model predicts a mix of epithelial, mesenchymal and hybrid E/M states (**Fig. 5C/S15C**). As the density increases, mesenchymal cells shift to an epithelial - hybrid E/M mix generated by crowded cells undergoing full MET, followed by partial EMT whenever space opens (**Fig. S16**) – an event that disappears in a confluent epithelium. These behaviors are independent of update, though the asynchronous version of our model has lower proliferation and *E-cadherin* knockdown thresholds (**Fig. S14**), but higher resistance to cell density-mediated MET (**Fig. S15**). In addition, the model’s behaviors are robust to several simultaneous random mutations (**Figs. S9-10**).

**Figure 5.**
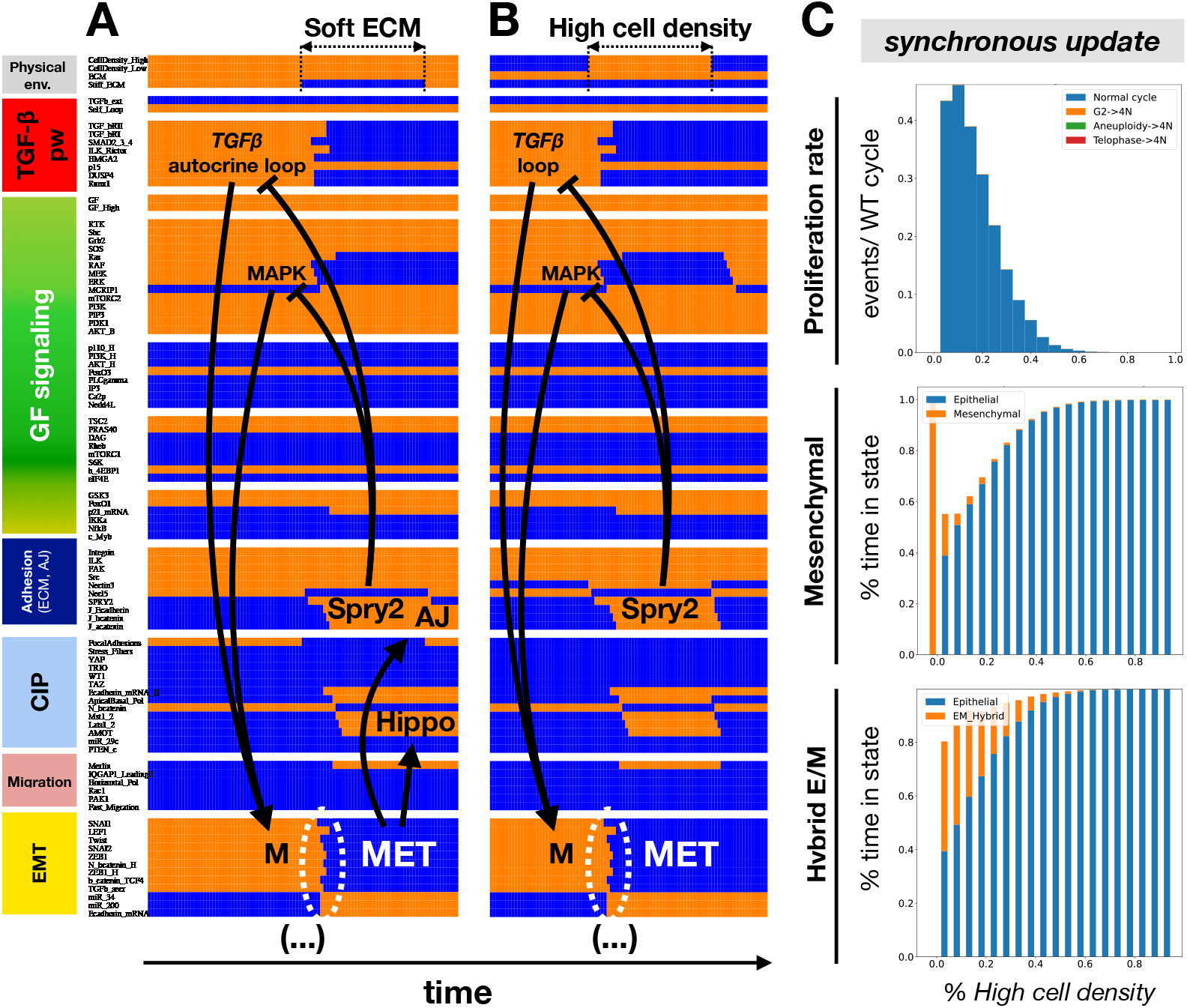
Soft ECM and high cell density work together to break the autocrine *TGFβ* loop and trigger MET via contact-induced *Sprouty-2 (Spry2*) activation. **A-B)** Dynamics of regulatory molecule expression/activity during exposure of (A) a growth-stimulated mesenchymal cell on a stiff matrix but at very high density to a soft ECM, and (B) an isolated mesenchymal cell on soft ECM to high density. *X-axis:* time-steps; *y-axis:* nodes organized in regulatory modules; *orange/blue:* ON/OFF; *black/white labels & arrows:* molecular changes that drive MET. **C)** Response of mesenchymal cells at the edge of a monolayer to increasing cell density **i**n the p**r**esence of 95% sa**tur**ating **g**rowth stimuli on 95% stiff ECM. *Top:* rate; of normal cell cycle completion (*blue*) vs. G2 → G1 reset (*orange*, not observed), aberrant mitosis (*green*, not observed), or failed cytokinesis (*red*, not observed) followed by genome duplication, relative to the minimum cell cycle length (21 time-steps), shown as stacked bar charts. *Middle:* fraction of time spent in a mesenchymal (*orange*) vs. epithelial (*blue*) state. *Bottom:* fraction of time spent in a hybrid E/M (*orange*) vs. epithelial (*blue*) state. *Total sampled live cell time:* 100,000 steps; synchronous update; *Initial state for sampling runs:* isolated epithelial cell in low mitogens on a soft ECM.

### Modeling Anoikis Resistance: mesenchymal cells lose anchorage dependence

A concerning consequence of EMT in the context of metastatic cancer is loss of anchorage dependence, or resistance to anoikis in mesenchymal cells ^57^. To test whether our model can reproduce this, we simulated loss of anchorage with an epithelial cell in low growth factor conditions (**Fig. S17A**) vs. a mesenchymal cell sustained by autocrine *TGFβ* signaling in the complete absence of growth factors (**Fig. S17B**). As expected, the epithelial cell underwent apoptosis while the mesenchymal one did not (**Fig. S17A-C**). Overexpression of either *AKT* or *ERK* rescued epithelial cells from anoikis (**Fig. S17D**, *left*). In mesenchymal cells, however, even complete knockout of *AKT* was not sufficient to induce anoikis, as *TGF-β*-mediated ERK activity could maintain survival. In contrast, *ERK* knockdown could re-sensitize these cells to anoikis (**Fig. S17C**, *right*).

### External TGFβ triggers EMT and cell cycle inhibition on stiff ECM, but apoptosis on soft substrates

*TGFβ* was shown promote EMT on stiff matrixes, but cause apoptosis on soft ECMs in the same epithelial culture ^46^. To test whether our model can reproduce both outcomes, we simulated cells at the edge of a monolayer (medium density) on soft vs. stiff ECM and exposed them to external *TGFβ*. Indeed, our model epithelial cells underwent apoptosis on soft ECM, an event driven by a combination of *BIM* overexpression, *BCL-XL* downregulation, and critically, lack of both *ERK* and high *AKT* activity (**Fig. 6A/S18A**). The latter is caused by weak activation of both mitogenic pathways in the absence of strong focal adhesions and stress fibers ^86,87^. In contrast, on stiff ECM *TGFβ* signaling activates both pathways, counteracts its own apoptotic effects, and triggers EMT instead (**Fig. 6B/C** and **S18B/C,** *low / high growth factor env*.). As cell density increases, the ability of *TGFβ* to trigger EMT weakens ^59^. Indeed, our model predicts that fully confluent monolayers dense enough to forbid stress fiber formation and at least some *YAP* activity stop responding to *TGFβ*, even if its receptors are accessible to external *TGFβ* (**Fig. 6D/S18D**). In a microenvironment with saturating growth signals and 50% external *TGFβ*, most cells on soft ECM have an epithelial phenotype and are prone to apoptosis, but also to occasional cell cycle entry (**Fig. 6E**, ~20% of saturating proliferation rate on ~20% stiff ECM with synchronous update). As matrix stiffness increases, the rate of both proliferation and apoptosis drops, giving way to EMT. Our asynchronous update results are qualitatively similar, though the model is substantially more sensitive to both *TGFβ-induced* apoptosis and EMT (**Fig. S18E**). In addition to its robustness to update, the model’s behaviors are largely unchanged by mutations up to the lockdown of 5 random nodes (**Fig. S19A**) or removal of ≥10 links (**Fig. S19B**).

**Figure 6.**
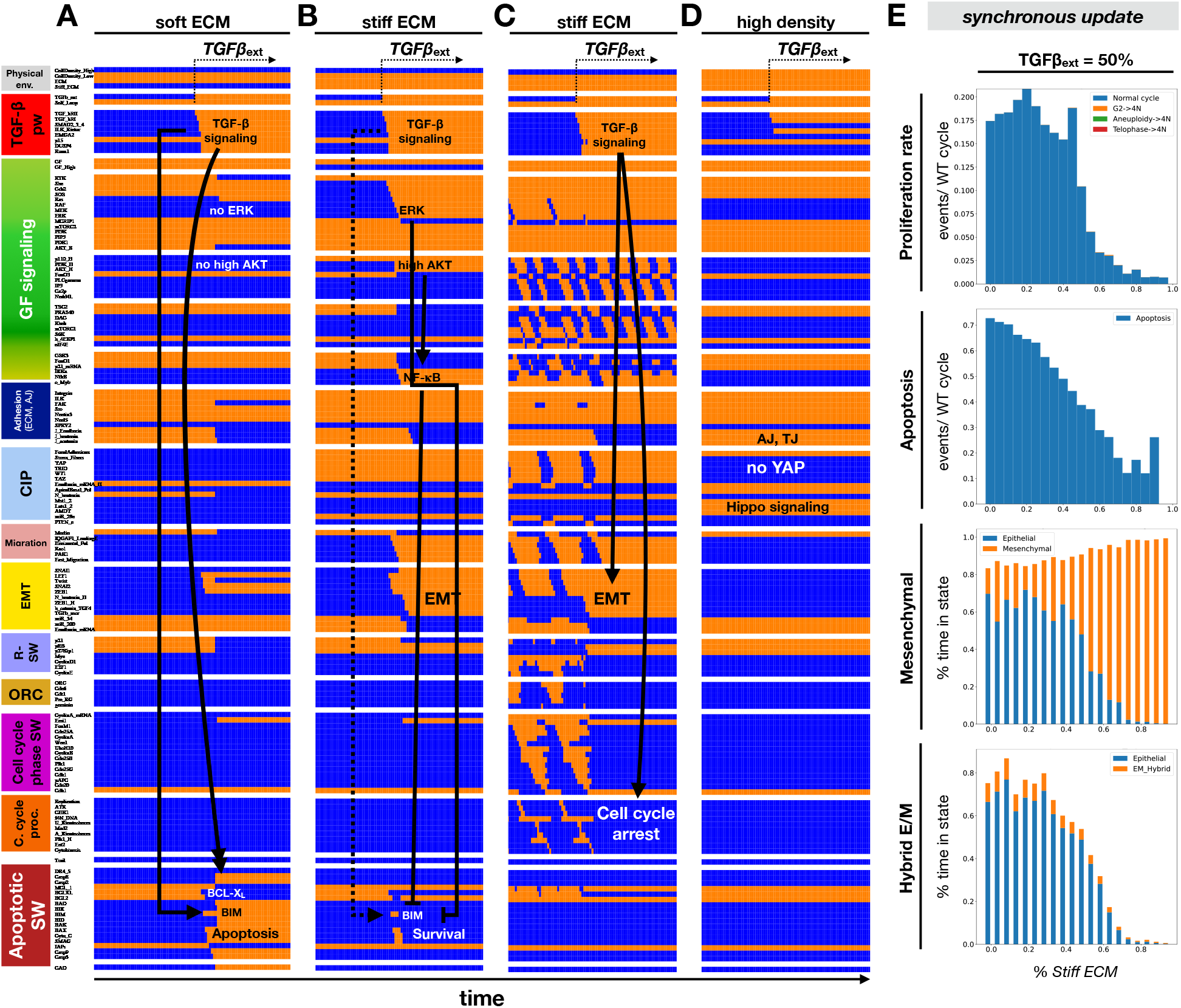
Exposure to external *TGFβ* leads to apoptosis on soft ECM, but EMT and cell cycle arrest on stiff ECM. **A-D)** Dynamics of regulatory molecule expression/activity during exposure to *TGFβ* of (A) growth-stimulated epithelial cells on soft ECM (medium density), (B) epithelial cells in low growth conditions on stiff ECM (medium density), (C) cycling hybrid E/M cells on stiff ECM (medium density), and (D) growth-s**t**imulated but fully confluent (dense) epithelium on stiff ECM. *X-axis:* time-steps; *y-axis:* nodes organized in regulatory modules; *orange/blue:* ON/OFF; *black/white labels & arrows:* molecular changes that drive apoptosis (A), EMT (B-C), or lack of response (D). **E)** Response growth-stimulated epithelial cells at the edge of a monolayer to 50% external *TGFβ* at increasing ECM stiffness. *Top:* rate of normal cell cycle completion (*blue*) vs. G2 → G1 reset (*orange*, rare but observed between 50-70% ECM stiffness), aberrant mitosis (*green*, not observed), or failed cytokinesis (*red*, not observed) followed by genome duplication, relative to the minimum cell cycle length (21 time-steps), shown as stacked bar charts. *Second from top:* rate of apoptosis (per minimum cell cycle length). *Third from top:* fraction of time spent in a mesenchymal (*orange*) vs. epithelial (*blue*) state. *Bottom:* fraction of time spent in a hybrid E/M (*orange*) vs. epithelial (*blue*) state. *Total sampled live cell time:* 100,000 steps; synchronous update. *Initial state for sampling runs:* dividing hybrid E/M at saturating mitogen exposure, at the edge of a monolayer on stiff ECM.

To test the effects of weakening autocrine *TGFβ* signaling, we started with cells exposed to near-saturating growth signals and scanned their responses as a function of increasing ECM stiffness (**Fig. S20A,***y axis*) at densities increasing from isolated to that of a monolayer’s edge (**Fig. S20A,** *x axis*). As **Fig. S20A** illustrates, even a 10% loss of autocrine signal strength in isolated mesenchymal cells (modeled as *Self_Loop* = OFF for a random 10% of time steps), restored proliferation on stiffer ECMs at *all* densities, rather than only in hybrid E/M cells capable of maintaining adherens junctions at a monolayer’s edge (**Fig. S20A***, top*). Intriguingly, our model predicts a small uptick of cell cycle errors such as slippage from G2 back to G1 and failure to complete cytokinesis following telophase, both more common among sparce cells (~ 2-3% of all divisions). An increase of apoptosis with the weakening of autocrine signaling is also predicted on stiff ECMs at very low density, at a slightly higher rate than cell cycle errors. Meanwhile, weakening the autocrine loop severely destabilized the mesenchymal state at very low density, triggering MET on all but the stiffest ECM (**Fig. S20A***, bottom 3 panels*). Next, we repeated the above scan with cells on 95% stiff ECM, scanning their responses as a function of increasing mitogen exposure (**Fig. S20B,** *y axis*) at increasing densities (**Fig. S20B,** *x axis*). In this case a weaker loop increased the prevalence of cell cycle errors even more (**Fig. S20B,** *top panels*), but more importantly reduced the stability of mesenchymal states in favor of hybrid E/M.

To summarize our results, we recreated the map cell phenotypes our model creates in different microenvironment-combinations, accounting for the availability of external *TGFβ* as a 4^th^ dimension (**Fig. 7**; complete list of detected model attractors, including those with a density-dependent *TGFβ* loop and *Trail=ON*, are summarized in **Table S2** and listed in **Table S3**). In contrast to the model with no autocrine *TGFβ* signaling (**Fig. 3E**), the quiescent mesenchymal phenotype is stable in the absence of external *TGFβ* in all environments except on very soft ECMs at very high density – assuming a strong autocrine signaling loop (**Fig. 7***, left panel, red stars*). In contrast, ongoing and saturating external *TGFβ* eliminates the hybrid E/M phenotype characteristic of monolayer edges, while the epithelial phenotype is only stable at very high density (**Fig. 7***, right panel*). Interestingly, our model predicts that cells at this high density do not undergo apoptosis in response to *TGFβ;* rather they use basal AKT activation by *TGFβ* as a survival signal even in the absence of all other mitogens, or in the absence of anchorage – though they cannot survive the absence of both (**Fig. 7***, right panel, green boxes*).

**Figure 7.**
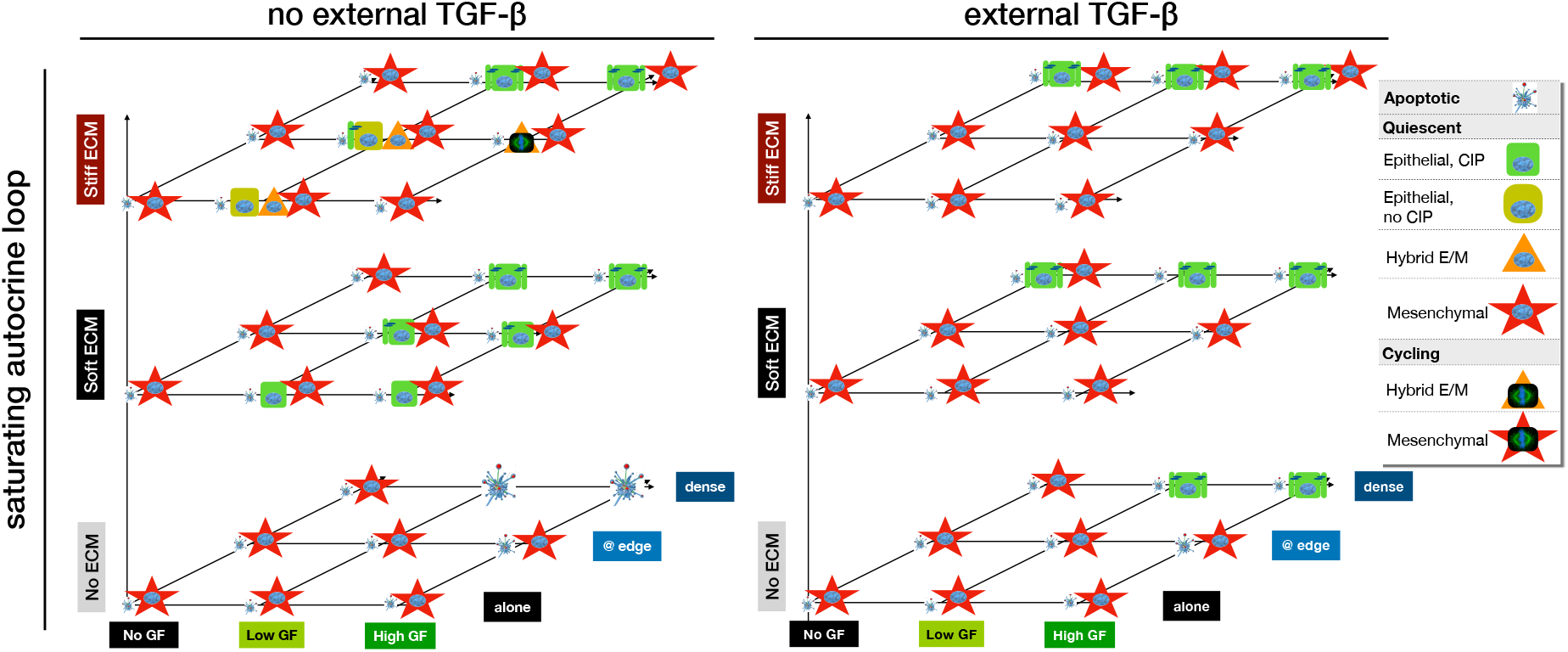
Strong autocrine *TGFβ* signaling stabilizes the mesenchymal state and external *TGFβ* forces full EMT in all but dense monolayers on soft ECM. Summary of cell phenotypes predicted for every combination of no/low/high growth-factor (*x axis*), no/low/high cell density environments (*y axis*), and no/very soft/very stiff ECM (*z axis*), in the absence (*left*) and presence (*right*) of external *TGFβ. Blue fragmented cell:* apoptotic state; *green box*: quiescent epithelial cell; *orange triangle:* hybrid E/M cell, *red star:* mesenchymal cell; *symbol with blue nucleus:* non-dividing cell; *symbol with mitotic spindle:* proliferating cell. Image credits: apoptotic cell ^118^; mitotic spindle: *https://en.wikipedia.org/wiki/Cell_division#/media/File:Kinetochore.jpg.*

## DISCUSSION

Modeling the signaling networks responsible for EMT has been an ongoing focus for the last decade ^22,26,27,30,35–42,71,88,89^. Models range from small ODE-based circuits that examine the core feedback loops responsible for locking in EMT to large Boolean networks that focus on the extensive crosstalk between EMT-promoting biochemical signals. Population-level models, while often considering the 2-3D biophysical environment of individual cells, do not integrate the effects of mechanosensitive molecular pathways such as *Hippo, YAP, Rac1/PAK1* and *NF-κB* ^37,90^. Thus, the biomechanical influences that control partial vs. full EMT, with or without transforming signals, are not adequately linked to molecular mechanisms responsible for each outcome. To address this, we first built a 136-node Boolean regulatory network model of partial as well as full EMT, driven by a cell’s biomechanical environment in the absence of transforming factors such as *TGF-β, Wnt, Notch, Hedgehog* or *IL6* (**Fig. 1C**). Instead, EMT in this model is triggered by mitogenic signals on stiff ECM, together with a lack of contact inhibition leading to migration. Namely, *Rac1/PAK1* activation and *PI3K/AKT-mediated NF-κB* signaling kickstart EMT by increasing the expression, localization, and stability of *SNAI1.* In line with most published models, our network can maintain distinct epithelial, hybrid E/M and mesenchymal states (**Figs. 1B, 3E**). Unlike other models, we also track the coordination between EMT, migration, cell cycle progression and apoptosis. With the aid of mechanosensitive inputs such as low density, our model can reproduce EMT driven by strong mitogens such as EGF on stiff ECMs (**Figs. 2,4, S1B-C**) ^52,53^, along with the observations that loss of *E-cadherin* can promote EMT (**Fig. S1D, S14C**) ^54,55^.

Our extended 150-node model endowed with a TGF*β* signaling module (**Fig. 4A, S7**) reproduces the need for a potent autocrine TGF*β* loop to maintain the mesenchymal state on soft ECM (**Fig. 4B**), in the absence of strong mitogens (**Fig. 4C**), or at high density (**Fig. 5A,** *left*). While this autocrine loop is a common feature to most published EMT models, here we also reproduce TGF*β*-induced apoptosis on soft ECMs vs. EMT on stiff ones (**Fig. 6A-B**) ^46^, TGF*β*-induced inhibition of proliferation in mesenchymal cells (**Fig. 6C**) ^56^, the lack of TGF*β*-induced EMT in very dense monolayers (**Fig. 6D**) ^59^, and mesenchymal anoikis resistance sustained by autocrine TGF*β* (**Fig. S17**) ^57,58^. This board repertoire of behaviors is a direct result of a modular modeling strategy by which we built the EMT transcriptional network as an extension of a model series that started with the detailed control of cell cycle progression ^82^, its coordination with apoptosis ^80^, then examined the effects of contact inhibition and loss of anchorage on both phenotypes ^45^.

In addition to reproducing known cell behaviors, our model offers several mechanistic, experimentally testable predictions:

1. In our model, the density of adherens junctions around a single cell is a biomechanical signal that biases EMT towards hybrid E/M vs. a mesenchymal outcome. Thus, we predict that relatively isolated cells in sparsely plated epithelia exposed to *EGF* on a stiff matrix are more likely to undergo full EMT, whereas cells at the edge of a monolayer or with a similar contact-surface stop at hybrid E/M instead (**Fig. 2**). This can be tested with single-cell resolution live-imaging experiments tracking junctional E-cadherin localization, along with β-catenin levels/localization or miR-200 activity (junctional vs. cytosolic vs. nuclear β-catenin live reporters are descried in ^91^; live E-cadherin probe on an orthogonal imaging channel in ^92^; live miR-200 probes can be designed according to ^93^). Specifically, cells imaged at increasing density on stiff ECM (> 100 kPA) in standard growth-promoting media are predicted to generate three distinct subpopulations as a function of junctional E-cadherin: *i*) cells with low nuclear β-catenin (or high miR-200) and high E-cadherin surrounded by neighbors (epithelial), *ii*) cells with moderate nuclear β-catenin, substantial junctional E-cadherin and some free edges (hybrid E/M), and *iii*) a high nuclear β-catenin group with no E-cadherin and few/no neighbors (mesenchymal). We further expect their balance to shift from mostly mesenchymal at very low density towards hybrid E/M, then to epithelial in very dense monolayers. In contrast, data showing no correlation between local cell density (single cell neighborhood) and nuclear β-catenin and/or miR-200 expression would refute our predictions and necessitate a reevaluation of our understanding of mechanosensitive EMT.
2. We predict that short, intermittent mitogen bursts that lack the strength to induce cell cycle entry can drive sparsely plated epithelial cells into a metastable migratory hybrid E/M state, which is maintained by uninterrupted migration (**Figs. 2D, S11A, S13C**). As a corollary, we predict a biphasic response to growth signal exposure where moderate signaling results in the largest fraction of hybrid E/M cells (**Fig. S14B**), while saturating mitogens interrupt migration and reset cells at each mitotic rounding (**Figs. S11B, S13B**). This could be tested with an experimental setup involving transient growth factor (or serum) exposure followed by washout, repeated with increasing exposure lengths and/or growth factor concentrations^79^. Cells would be live-imaged before, during and after this pulse to track their migration speed, nuclear β-catenin levels, and division. Our model predicts that shorter/weaker pulses preferentially generate persistent migratory cells with moderate nuclear β-catenin (higher than the pre-pulse baseline), while longer/stronger pulses result in divisions, followed by an immediate drop in daughter cell migration. We also expect nuclear β-catenin to be highest in cells imaged after exposure to moderate pulse length/strength (imaged around the time of appearance of daughter cells). Here, it is possible that the migration triggered by growth factor pulses is not as persistent in live cells as our deterministic model predicts. Yet, observing some persistent mitigation and/or sharp drops in speed following mitosis would still validate our model. In contrast, observing a loss of migration immediately upon growth factor withdrawal would require us to reevaluate the bistability of our migration module.
3. We predict that the biomechanical environment alone can trigger MET by blocking stress fiber formation and muting *ERK/AKT* signaling, while *Pak1* hyperactivation can reestablish hybrid E/M (**Fig. 3**). Furthermore, soft ECM and high cell density cooperatively break the autocrine *TGFβ* loop and trigger MET via contact induced *Sprouty 2* activation (**Fig. 5**). This could be tested by triggering full EMT with *TGFβ* in a sparse population, replating them at very high density on a soft ECM (~ 0.5kPa), then imaging the reappearance of junctional E-cadherin followed by phospho-AKT or phospho-ERK staining ^94^. Here we expect to see MET, preferentially in the densest areas of the plate. Repeating the imaging in the presence of *Pak1* overexpression and *Sprouty 2* knockdown would further test the accuracy of our model’s molecular mechanism driving mechano-sensitive MET.
4. In line with previous model predictions and observations about the asymmetry of EMT vs. MET ^27,95^, our model does not undergo a direct transitions from mesenchymal to hybrid E/M state. Rather, model environments that favor hybrid E/M over the mesenchymal state induce full MET followed by partial EMT (**Fig. S16**). This can be tested by live-imaging nuclear β-catenin or miR-200 with junctional E-cadherin in a series of experiments like the one above, at increasing ECM stiffness (0.5 kPA to 100 kPA) and increasing cell densities (sparse to dense enough to forbid stress fiber formation). According to our model, a random selection of mesenchymal cells plated onto moderately stiff ECMs at medium density should undergo an abrupt drop in nuclear β-catenin followed by the re-expression of E-cadherin, appearance of adherens junctions, and a moderate β-catenin re-accumulation at the edges of dense clusters.
5. While TGF*β*-induced EMT requires stiff ECM, we predict that the mesenchymal state can be maintained on soft ECMs by autocrine TGF*β* (**Fig. 4B**). The above setup with very low density and soft ECM should show a robust preservation of the mesenchymal state (no drop in nuclear β-catenin), while the same experiment in the presence of a TGF*β* signaling inhibitor (e.g., a dominant negative soluble receptor that weakens the autocrine loop) is expected to show MET.
6. Finally, we predict that at low-to medium cell density and medium-strength growth signals, autocrine TGF*β* signaling that does not saturate the TGF*β* pathway (does not reliably guarantee its downstream effects) will generate a dynamic mix of mesenchymal and hybrid E/M cells, the latter of which proliferate and thus continually reset to undergo EMT anew (**Figs. S20**). This is accompanied by a small uptick in apoptosis as well as cell cycle errors that generate polyploidy (specifically, slippage from G2 and/or telophase to G1) in the absence of any mutations related to checkpoint control (**Figs. S20**). This can also be tested with the setup above, using stiff ECMs but a range of growth factor levels. Live imaging that tracks key cell state markers (nuclear β-catenin, junctional E-cadherin, miR-200) along with division and apoptosis could identify the hybrid E/M → epithelial transition following mitosis, as well as the link between non-saturating autocrine signaling and apoptosis. Finally, testing the DNA content of the cells at the end of these live imaging runs could test our cell cycle error predictions.

Rather than our ability to reproduce known cell behaviors, we believe that the above predictions represent a more important contribution to characterizing the roles of partial vs. full EMT in non-oncogenic processes such as wound healing. Testing them could boost the credibility of our model in predicting the effects of targeted interventions. Alternately, failed predictions would point to unexpected gaps, helping us further probe the behavior of cells in different biomechanical environments. Ultimately, correct integration of mechanosensing and transforming signals will be required for building predictive cell population models that follow the 2D or 3D processes of wound healing, the initiation of cancer invasion, or the establishment and growth of metastatic lesions.

As detailed under *Limitations of the Study* below, it is possible that the TGF*β*-mediated cell cycle block is too strong in our model, and the experiments described above would show proliferation in fully mesenchymal cells in spite of potent autocrine TGF*β*. Yet there is substantial experimental evidence for the tug of war between proliferative potential and the migratory mesenchymal state ^7^. For example, hepatic oval cells capable of regenerating the liver when hepatocyte proliferation is compromised where shown to have a hybrid E/M phenotype ^96^. Moreover, cancer stem cells in squamous cell carcinoma and ovarian cancer exist as a heterogeneous mix of proliferative hybrid E/M and mesenchymal states, and they dynamically regenerate this mix from either subpopulation ^97,98^. Indeed, *in vivo* metastatic tumor growth in ovarian and breast cancer is primarily due to hybrid E/M cells ^101,102^. We can model this division/migration conflict in environments that can induce and maintain EMT biomechanically, and thus do not rely on uninterrupted autocrine TGF*β* signaling (**Fig. S20**). What our model currently does not account for, however, is the connection between the hybrid E/M state and the regulatory network responsible for a stem cell-like state, observed in all forms of EMT (i.e., developmental, regenerative, and pathological) ^30,99^. Integrating a stemness module, along with an explicit accounting for the likely context-dependent role of ‘phenotypic stability factors’ such as *OVOL* and *GRHL2* known to stabilize the hybrid E/M state and promote stemness, would be an important next step in delineating the similarities and differences between EMT in different tissue and disease contexts ^30,100^.

In addition to limitations, it is important to delineate the boundaries of our model; cell behaviors linked to EMT that we explicitly do not address. Two of these stand out as key to our ability to integrate our regulatory network into multiscale models of tissue behavior. *First*, our model does not address the patterning effects of *Notch* signaling in a population of cells undergoing EMT. *Notch* not only promotes EMT by inducing *Snai1* ^39^, but also generates distinct patterns between neighboring cells depending on the ligands expressed on neighbors ^37^. Cells expressing *Delta* ligands promote lateral inhibition patterns that alternate *Delta*^High^/*Notch*^Low^ cells with the opposite phenotype. This is central to the endothelial to mesenchymal transition, as it generates alternating patterns of tip cells and stalk cells required for sprouting angiogenesis ^71,101^. In contrast, the *Jagged* ligand promotes lateral induction, leading to a strong clustering of similar cells ^102^. Previous computational models of *Notch* signaling showed that lateral induction via *Notch-Jagged* can generate intermingled clusters of epithelial, mesenchymal, and hybrid E/M cells akin to the patterning seen in tumors ^37^. This pattern formation is heavily influenced by the tissue microenvironment such as inflammatory signals (*IL-6, TNFα*) ^103,104^, *Wnt* and *Hedgehog* signaling ^105,106^, as well as hypoxia ^107^, most of which promote EMT and induce *Jagged.*

The second direction that merits our attention is the relationship between EMT and senescence. The process of senescence makes short-term use of damaged cells by shutting down their ability to divide, but using them to boost survival, proliferation and EMT in their surroundings ^108^. While senescent cells trigger their own immune clearance, their slow accumulation is accelerated by chronic damage such as inflammation or oxidative stress and leads to tissue aging ^109^. There is increasing evidence that senescence and the mesenchymal state are mutually exclusive in individual cells ^110,111^. Mesenchymal cells are largely protected from senescence, while senescent cells do not respond to signals that trigger EMT ^112^. During localized healing of an epithelium, this creates a useful distribution of labor: damaged cells go senescent but induce division and EMT in neighbors. These mesenchymal cells are then protected from senescence until they reestablish an intact epithelium. As neither senescent nor mesenchymal cells contribute to barrier formation, this can accelerate dysfunction in aging tissues ^113^. Worse, in epithelial tumors this crosstalk protects invasive but damaged cells from senescence, aiding metastasis ^114^. Yet, the two phenomena are usually studied in isolation, and the relationship between hybrid E/M cell states and senescence remains unexamined. Building a predictive molecular model of the mechanisms that forbid their coexistence and probe the transition from healing to the pathological roles of EMT and senescence is a key next step that benefits from our current integration of EMT with cell cycle control, apoptosis, and contact inhibition. Ultimately, we envision these single-cell network models as the engines that drive 3D tissue simulations, able to predict effective interventions that can nudge diseased tissues back to health.

## LIMITATIONS OF THE STUDY

Our models have several limitations that can be addressed by future refinement; some related to the way our previous contact inhibition model’s environment was built, some specific to EMT and TGF*β* signaling. As detailed in ^45^, we use a crude approximation of the spatial asymmetry of *Rac1/Pak1* activity at the leading edge and a minimal migration module. Yet, feedback between cell contractility and ECM stiffness – itself modulated by the migrating cell – was shown to render the behavior of invading cells nonlinear, such that intermediate stiffness is optimal for high-speed migration ^115^. Moreover, hybrid E/M cells use their lower contractility to move faster and invade more effectively than mesenchymal cells ^116^. The migration module of our model needs significant refinement to address these phenomena before it is suited to model collective behavior such as wound healing in 2D. Other areas of improvement inherited from ^45^ hinge on the limits of how well Boolean models capture gradual changes in ECM stiffness and cell density (detailed in Supplementary Methods and ^45^).

Another limitation relates to our Boolean approximation of the biophysical microenvironment. Namely, our model draws explicit distinctions between ECMs that do or do not support sufficient adhesion for survival, then another one between ones that do or do not aid force generation required for both migration and cell cycle entry. Similarly, our cell density nodes draw a sharp distinction between isolated cells that cannot form any adherens junctions and models slightly higher densities with intermittent junction formation. Another boundary is between cells that have room to polarize horizontally vs. those that are so dense that they cannot form stress fibers. While our model can modulate stiffness and/or density within these two regimes by toggling their inputs ON/OFF with a tunable probability, it does not allow overlap between the two regimes. Such artificial boundaries between district cell response regimes could lead to mismatch with experimental data.

Finally, a potential limitation related to our integration of the EMT switch with the cell cycle is that our model’s mesenchymal cells do not show anchorage-independent growth ^7,117^. While we reproduce anoikis resistance, our cell cycle control relies entirely on spreading on a stiff matrix. A deeper examination of how cancer cells acquire anchorage-independent growth, and whether there are pre-requisite mutations that could drive such growth in our current model, requires further work. A related potential problem is that our mesenchymal cells with strong autocrine TGF*β* loop do not divide at all, owing to a TGF*β*-mediated cell cycle block. Instead, in environments that do not force full EMT (sub-optimal matrix stiffness, non-saturating autocrine TGF*β* signaling, moderate density, non-saturating growth signals) our model predicts a fluid coexistence of mesenchymal and hybrid E/M cells, the latter of which divide with high frequency but then briefly reset to an epithelial state before undergoing EMT once more. It is possible that our model’s TGF*β*-mediated cell cycle block is too strong, in that fully mesenchymal cells with sufficiently potent autocrine signals to keep them mesenchymal may nevertheless divide.

## Supporting information

File S1 - Biological justification for models

Files S2-S10 (models in different formats)

Supplemental Tables S1-S3 (all model attractors)

## ACKGNOWLEDGEMENTS

We thank Peter Regan for his work on designing the Dynamically Modular Model Specification (*.dmms*) file format, developing the *dynmod* software used to parse our model files, and helping us create code repositories for publication. All funding for this study was provided by the College of Wooster. E.R. would like to thank the Henry Luce III Fund for Distinguished Scholarship. E.G and I.Z thank the Henry J. Copeland Independent Study Fund for research support. A.B. thanks APEX (Advising, Planning, & Experiential Learning) at the College of Wooster for support to present their work.

## AUTHOR CONTRIBUTIONS

E.S. and E.G. designed the first version of our Boolean EMT switch and started its integration with the model in ^45^, leading to the first mechanosensitive version. M.H., A.B. and I.Z. tested and refined the model to converge in the version included here. E.R.R. supervised the research, analyzed/refined the final models, ran final simulations, and wrote the paper with edits from E.S.

## STAR Methods

### RESOURCE AVAILABILITY

#### Lead contact

Further information and requests for help with code or simulation interpretation will be fulfilled by the lead contact, Erzsébet Ravasz Regan.

#### Materials availability

This study did not generate any wet-lab materials or resources (entirely computational).

#### Data and code availability

All original code has been deposited at Zenodo and is publicly available as of the date of publication. DOIs are listed in the Key Resources table. These include the Haskell code package *dynmod* used to parse .*dmms* model files (see below), generate network visualizations, and render the LaTeX-formatted supplementary file containing the model’s biological justification. The *dynmod* parser generates files read by a C++ XCode package called *ReganLabBooleanSims*, used here to run all *in silico* experiments. Instructions to reproduce all figures are detailed below and included with each code packet.

Any additional information required to re-run the simulations reported in this paper is available from the lead contact upon request.

### METHOD DETAILS

#### Boolean model building

Boolean network models focus on the complex combinatorial logic by which molecular species and pathways intersect, sacrificing the precise concentration and time-dependence of interactions in favor for scalability ^43,119^. The ON (expressed and active) vs. OFF (not expresses or inactive) response of each molecule to every combination of its regulators dictate the model’s temporal dynamics. Ideally, Boolean regulatory rules should be directly inferred from experimental data. Yet, detailed computer-accessible time-series data on the dose-dependent response of single cells to their biochemical environment, along with molecular activity responsible for these responses are not currently available. Instead, we compiled qualitative experimental data from 479 papers into a 150-node Boolean model in which each of the 630 links is justified by experiments (**File S1**; *blue:* changes from ^45^; *green:* further additions in version with *TGFβ*).

##### Defining and refining the model’s Boolean regulatory logic

Boolean gates were inferred from literature and/or chosen based on reported effects of molecular input combinations. These gates are further tuned, within the constraints of experimental literature, to a) generate model behaviors that match all known cell behaviors within the scope of the model and b) eliminate phenotypically similar attractor states within identical environments where known expression / activity pattern of the molecules that differ between these attractors single out one of these states as the most biologically accurate. This fine-tuning is most common for complex gates with several inputs, where preserving the known activating / repressive influence of individual inputs, along with known synergy or independence between pairs, is insufficient to fully specify the Boolean gate. In such cases, the model’s dynamical behavior across all environments and environment changes is used to further constrain the gate. Namely, most gate options allowed by experimental data specific to its output node can be eliminated because they produce non-biological behaviors in some context (e.g., non-biological stable state, loss of an expected state, incorrect transition dynamics). Fixing these behaviors occasionally necessitates the inclusion of additional regulators (node) or interactions, always drawn from the experimental literature.

During this initial construction phase we use synchronous update ^120^, where every node of the network updates synchronously in every time-step. The advantage is that it renders the dynamics of the system completely deterministic, allowing us to account for the precise role of each molecular species at every causal step along a biological process and compare it to experimental data. By building a synchronous Boolean model first, we could identify the molecular causes of behaviors that deviated from known cell dynamics and iteratively fix the model by altering / expanding the logic gates leading to non-biological behavior. Our analysis of the model’s robustness to errors indicates that the average dynamical behavior is difficult to alter by a random change to a few logic gates (**Figs. S4, S9, S10**).

##### Storing Boolean models in Dynamically Modular Model Specification (.dmms) files

To keep track of all model metadata such as its modular structure, experimental data justifying each node, link and gate, module structure, node visualization data, etc., we collated all model-specific information into human-editable and machine-readable files (**Files S2**, **S3**; do not rename). These *.dmms* files are parsed by the *dynmod* software package written in Haskell. The command

~~~
dynmod -t -e EMT_Mechanosensing.dmms
~~~

parses **File S2**, exports the model in BooleanNet format (**Files S4**-**S6**), and generates a folder of truth tables and graphical information read by *ReganLabBooleanSims (C++ code*) used for running virtual experiments.

##### Generating and storing network visualizations

Model files in *.dmms* format allow us to store node layout and color to ease the iterative visual rendering of large networks. The command

~~~
dynmod -g EMT_Mechanosensing.dmms
~~~

generates a *.gml* graphical layout using the coordinates and node colors stored in the file, if any. This file can be read and organized by the yED graph visualization software ^121^, and re-saved as *.gml* (**Files S7**, **S8**). Assuming that the network structure is completely unchanged by these edits, *dynmod* can read and update coordinates and colors in the .dmms file to preserve the layout:

~~~
dynmod -u EMT_Mechanosensing.gml EMT_Mechanosensing.dmms
~~~

##### Generating Supplementary Tables with detailed model justification

The command

~~~
dynmod -s EMT_Mechanosensing.dmms
~~~

generates a formatted LaTeX document (.tex) and bibliography file (.bib) with a series of tables that organize the model’s nodes by modules and use metadata from the .dmms file to detail the biological role of each node, the molecular mechanism of action for each link, and the assumptions made in constructing large Boolean logic gates. These can be typeset using LaTeX to generate the large, referenced table in **File S1**.

#### Sampling the state space of a Boolean model to map its synchronous attractors

We used synchronous Boolean modeling to find every stable phenotype and/or oscillation (attractor) of the model ^45,80,82^. To do this we use a stochastic state space sampling procedure adapted from ^122^, implemented in *dynmod* and *ReganLabBooleanSims*. First, we implemented a noisy version of synchronous Boolean dynamics, in which each regulatory node is affected by a small amount of noise, *p*_noise_, in every time-step, resulting in the incorrect output ^123^. We then use the noisy dynamics to aid our sampling procedure by starting the network from a random initial condition and simulating a time-course of *N*_series_ noisy time-steps. As the model generates its noisy dynamical trajectory, the algorithm pauses at each state it visits to find the attractor basin this state would fall into if the dynamics were to continue in a deterministic fashion. In addition, the version implemented in *ReganLabBooleanSims* also scans the immediate neighborhood of this state by enumerating every state the system could reach from the current one via a single node-state flip and identifying their attractor membership (deterministic dynamics) ^82^. This allows the algorithm to access parts of the state space the noisy dynamics might never go near and find smaller basins. As a result, the algorithm in *ReganLabBooleanSims* is slow on random Boolean networks with large numbers of small basins. By contrast, our model’s attractor basins, representing robust phenotypes, are typically large and rapidly found. The full algorithm descried in ^82^ tracks the visitation probability of each state, basin, and transition (not used here). To run the noisy trajectory from multiple random initial conditions, our code automatically detects all environmental input nodes, partitions the full state space of the model into sub-spaces corresponding to each unique environmental node state-combination, and samples each subspace from *N*_md_ random initial conditions.

First, we used *dynmod* for an exhaustive attractor search with the parameters:

~~~
dynmod --grid 15,15,0.02,5 EMT_Mechanosensing.dmms
~~~

With these settings, *dynmod* re-runs the algorithm multiple times to generate a 15 by 15 grid with *N*_md_ ∈ {5, 10, 15, …, 75}, *N*_series_ ∈ {5, 10, 15, …, 75}, and*p*_noise_ = 0.02 (the last number in --grid 15,15,0.02,5 is the gap between increasing *N*_md_ and *N*_series_ values, the first two set the number of sampling-runs along the grid; ~ 3 hours on a 2022 Mac Studio for smaller model; ~ 81 hours for model with *TGFβ*). As its output, *dynmod* generates a heatmap of the number of attractors found in each individual sampling run (to test for sampling convergence) and collates all detected attractors into a single *.csv* file. Formatted versions in Excel, generated for both models with the above setting, are included as **Tables S1-3** (including a summary table for the larger model that marks attractors detected while using *ReganLabBooleanSims*).

Once the attractors are mapped, we use *ReganLabBooleanSims* for *in silico* experiments. Thus, *ReganLabBooleanSims* also samples the state space once, to generate unique attractor IDs corresponding to specific phenotypes in each desired environment -- needed to initialize time courses. To do this, *ReganLabBooleanSims* reads a *Virtual_Experiment_List.txt* file from the model’s folder (where required information generated by *dynmod* is also stored). This file starts with the following two lines:

~~~
EMT_Mechanosensing GF GF_High
Sampling 200 (Trail=0)
~~~

This initiates a sampling run with *N*_md_ = 200, *N*_series_ = 5, and *p*_noise_ = 0.02, with the *Trail* input node locked permanently OFF (we do not investigate *Trail* responses here, and this cuts the state space in half). Once the sampling is finished, *ReganLabBooleanSims* runs a time course from each non-apoptotic attractor by changing each environmental variable at a time, and identifies the attractor reached in the new environment, noting whether the sampling missed it. This is a very efficient way to detect live cell states in environments where most of the state space is occupied by apoptotic or other large attractor basins. **Table S2** marks several attractors identified by this process but not detected by *dynmod.*

*ReganLabBooleanSims* outputs a series of pdfs that show all detected attractors organized by the external environments they are stable in (e.g., **File S9**). These are marked by a numerical ID, as well as a quick visual indicator of their phenotype. The numerical IDs are *not* preserved between independent sampling runs but remain unchanged if *ReganLabBooleanSims* can read the results it saved from a prior sampling from the model folder (this also saves repeated samplings of unchanged models). The phenotype indicators are *orange circle* = quiescent; *red oval* = cell cycle; *orange circle with dark border* = apoptotic; *orange circle with vertical line* = quiescent, duplicated genome; *E/H_Migr/M* = EMT module in epithelial / hybrid / mesenchymal state. Lines indicate transitions in response to environmental change. This rudimentary attractor visualization can be used to specify initial phenotypes and environments for further experiments described below. **Figures 3E** and **7** offer a simplified illustration of the attractor repertoires of our models by omitting all apoptotic attractors in conditions with *ECM=0* and *Trail* =1 (**Fig. 3E**), or *Self_Loop* = 0 and *Trail* = 1 (**Fig. 7**). In addition, we omit all non-cycling live attractors with double DNA content (4N), but otherwise identical to their counterpart with 2N DNA (described in ^80,82^). Finally, omission of the *Trail* = 1 condition from ***Fig. 7*** also hides 24 quiescent mesenchymal attractors that appear resistant to *Trail* exposure (not discussed).

#### Automated isolation of a subnetwork from a larger (multi-switch) Boolean network

*ReganLabBooleanSims* can automatically isolate and examine the dynamical behavior of any network module, such as the EMT switch (**Fig 1A-B**). To this end, we have previously developed an algorithm that defines the Boolean gates of nodes when they lose some of their incoming connections ^82^. The main goal was to optimally preserve the regulation of a node by its remaining inputs. Briefly, whenever a subset of inputs is removed from a Boolean gate, the algorithm assumes that they are frozen into either an ON or an OFF state. To preserve the dynamical influence of the remaining nodes, *ReganLabBooleanSims* finds one of the 2^k^ possible combinations of frozen inputs such that: a) all remaining input nodes are functional, if possible and b) the entropy of the remaining Boolean gate fragment, *H_G_* = - *p* · log(*p*) - (1-*p*) · log(1-*p*), is maximal (*p* = fraction of OFF-outputs). **File S4** includes the resulting BooleanNet model file for the isolated EMT switch. To generate this file, as well as to detect and visualize the attractors of the isolated EMT module (along with the *Apoptotic_SW*, for example), *ReganLabBooleanSims* requires a *Virtual_Experiment_List.txt* file with the following line (-- denotes a comment):

~~~
-- Figure 1B
**Modules** (EMT, Apoptotic_SW)
~~~

#### Biased asynchronous update

Asynchronous update schemes change the state of one node at a time and use this new state as they update the node’s targets. Random order asynchronous models, the scheme adopted here, update every node in every time-step, but they do so sequentially in a random order re-shuffled before every step. Asynchronous update schemes are favored in biological modeling, as they simulate the unfolding of the same regulatory process along many slightly different paths, each with different likelihood ^124^, mimicking the stochasticity present *in vitro.* Moreover, asynchronous update eliminates potential artifacts of synchronous modeling, such as behaviors that rely on perfect and deterministic coordination of parallel signals. That said, they also generate biologically non-realistic sequences of molecular events by failing to follow up on the effects of short-lived signals that live cells reliably respond to. As a result, we use a hybrid update scheme developed in ^80^ where the update order of most, but not all nodes is random (biased asynchronous update). Specifically, we update 12 of the 150 nodes either first or last, depending on their correct state, as follows (for detailed justification for each node, see ^80^):

- Update at the ***start*** of each asynchronous time-step (first in list updated first): *Pre_RC* if ON, *Replication* if OFF, *U_Kinetochores* of OFF, *A_Kinetochores* if ON, *Plk1_H* if ON, *CyclinB* if ON, *Cdc20* if OFF.
- Update at the ***end*** of each time-step (first in list updated *last*): *Replication* if ON, *f4N_DNA* if ON or OFF, *Cytokinesis* if OFF, *Ect2* if OFF, *A_Kinetochores* if OFF, *CyclinE* if ON, *FoxM1* if ON, *Cdc20* if ON, *Plk1_H* if OFF

The update-robustness of our cell cycle / apoptosis model was examined in greater detail in ^80^.

#### Time courses with reversible environmental change

To test the model cell’s responses to reversible changes in their environment, we choose a specific phenotypic state in a desired initial environment as an initial condition, then specify the length of time we wish to flip one environmental variable. *ReganLabBooleanSims* shows the model’s state or behavior for 50 timesteps, flips the environment node for the designated length of time, then flips it back and further follows the network’s dynamics for a total of 400 time-steps. These *in silico* experiments for synchronous and biased asynchronous update are specified in the *Virtual_Experiment_List.txt* file as (bias rules are currently hard coded in *ReganLabBooleanSims*):

~~~
-- Figures 2A and S2A
**Pulse1** 41 Stiff_ECM 100
**Async_Pulse1** 41 Stiff_ECM 100
~~~

#### Modeling an ensemble of mutant networks (robustness to network errors)

To test the robustness of our models to random mutations and/or errors in construction, we generated three distinct types of mutant network ensembles. In the first ensemble, each network has *n*_node_ randomly selected nodes permanently locked ON or OFF (this too is random). In the second ensemble we remove *n*_link_ randomly selected links. Boolean gates for the targets of each such link are derived using the procedure used when isolating modules; namely the portion of the original gate where the removed input is locked in its least canalizing value is preserved. For the third ensemble, we choose *n*gate random nodes, and we flip a single random output value of their response logic table. This alters their response for a single, specific combination of input values.

The following virtual experiment line added to *Virtual_Experiment_List.txt* generates an expression/activity time course for all three ensembles (1000 mutant networks each) starting from an initial state matching that of attractor ID **31**, responding to 100 time-steps of high growth factor exposure (followed by its reversal), with 15 random errors / network in each ensemble.

~~~
-- for Figure S4
**ModelErrors_Pulse1** 31 GF_High 100 15
~~~

#### Modeling non-saturating inputs and partial knockdown / overexpression within a Boolean framework

To test the effect of non-saturating, intermediate environmental inputs we simulated the model while overriding one or more input node in each time-step with a stochastically generated ON/OFF value of a set probability, thus tuning the cell’s average ‘‘signal exposure” between 0 (none) to 1 (saturating). Similarly, we model partial or complete knockdown/ hyperactivation of a node by forcing it OFF/ON with a set probability in each time-step, otherwise leaving it to obey the Boolean rule that normally governs its behaviors ^45,80^. This offers a way to model partial inhibition / hyperactivation seen in cells or tissues treated with a drug (e.g., a chemical inhibitor), siRNA silencing, or a pool of a constitutively active proteins alongside the endogenous one. All these result in partial inhibition / saturation of the molecule, leading to blunted downstream effects. In contrast, full knockdown only mimics gene-edited null cell lines where both gene copies (or all homologues represented by a single Boolean node) were removed. Furthermore, as the intermediate levels we model represent cellular concentrations at which the targets our knocked down molecule influences are near the inflection point of S-shaped response curves (i.e., near their activation threshold), the effects of intermediate activity in real cells become highly noisy ^77,78^. Our method of randomly allowing a node to affect its targets or be locked OFF/ON mimics an intermittent signal that can stochastically activate its downstream cascade, but the signal flowing through the cascade is unreliable. This is a coarse method to model the coupled effects of losing some but not all signal propagation and the resulting noisy response.

The following *Virtual_Experiment_List.txt* lines generate a stochastic time course of the model’s dynamics in non-saturating environments, with or without partial node knockdown/hyperactivation (*ReganLabBooleanSims* expects a complete list of input node settings, as shown below):

~~~
-- Figure S16
**NonSaturating_Draw** 218 (GF=1, GF_High=0.95, TGFb_ext=0, Stiff_ECM=0.75,
Self_Loop=1, CellDensity_Low=1, CellDensity_High=0.2)
-- same time course with 95% junctional E-cadherin knockdown/overexpression **NonSaturating_Draw** 218 (GF=1, GF_High=0.95, TGFb_ext=0, _ECM=0.75, Self_Loop=1,
CellDensity_Low=1, CellDensity_High=0.2) (J_Ecadherin KD 0.95)
**NonSaturating_Draw** 218 (GF=1, GF_High=0.95, TGFb_ext=0, Stiff_ECM=0.75,
Self_Loop=1, CellDensity_Low=1, CellDensity_High=0.2) (J_Ecadherin OE 0.95)
~~~

#### Boolean network modules representing distinct cellular regulatory functions

##### Overview of previously published regulatory modules

1. <u>*Growth factor signaling*</u> (green node clusters on **Fig. 1C**, details in ^80^) is dynamic module tracking basal survival signaling through PI3K/AKT, the mitogen-induced MAPK cascade (*lime green*), cell cycle linked oscillations in PI3K/AKT signaling (*bright green*), and mTORC1 activation (*mustard*).
2. <u>*Replication origin licensing switch*</u> (light brown nodes on **Fig. 1C**, details in ^80^) is a bistable module that tracks licensing and firing of replication origins.
3. <u>*Restriction switch*</u> (lilac nodes on **Fig. 1C**, details in ^82^) is a bistable module that controls cell cycle entry at the restriction point.
4. <u>*Mitotic phase switch*</u> (purple nodes on **Fig. 1C**, details in ^80,82^) is a tri-stable module that controls the DNA damage and spindle assembly checkpoints, as well as the reset of mitotic cell cycle control molecules to a G1 state.
5. <u>*Apoptotic switch*</u> (dark red nodes on **Fig. 1C**, details in ^80^) is a bistable module that commits cells to apoptosis or maintains survival.
6. <u>*Cell cycle processes*</u> (dark orange nodes on **Fig. 1C**, details in ^80^) are a collection of processes during proliferation that track DNA replication, spindle assembly and cytokinesis without the need to model each in full molecular detail.
7. <u>*Adhesion*</u> (dark blue nodes on **Fig. 1C**, details in ^45^) are signaling pathways that mediate anchorage dependence (ECM adhesion via integrin receptors) and adherens junction formation (cell-cell adhesion via *Nectins* and *E-cadherin*).
8. <u>*Contact Inhibition*</u> (light blue nodes on **Fig. 1C**, details in ^45^) is a mechanosensitive bistable module that either promotes proliferation and migration via YAP/TAZ on stiff ECMs or maintains Hippo pathway activity and contact inhibition of both.
9. <u>*Migration*</u> (light orange/pink nodes on **Fig. 1C**, details in ^45^) is a bistable module that can sustain migration on stiff ECM via internal positive feedback. Namely, focal adhesion mediated *Rac1* activation at the leading and the establishment of lamellipodia support further focal adhesion formation powered by *Rac1*, while its effector *IQGAP1* promotes horizontal polarization and further concentrates *Rac1* at the leading edge.

##### Overview of regulatory modules introduced here

1. <u>*NF-κB pathway*</u> (in *“Connectors”* block, *dark green* nodes on **Fig. 1C, S3**) is a linear pathway that models *NF-κB* activation by mitogen- or *TGFβ-*induced AKT signaling ^125,126^, or by *PAK1* kinase ^127^. Thus, we linked *IKKα* activation to the *AKT_H* node and accounted for the degradation of *IκB* implicitly by assuming that *NF-κB* is on when either *IKKα* or *PAK1* release it by degrading *IκB.* In addition, *NF-κB* induces *c-Myb* expression (in parallel with *E2F1*) ^128^, which in turn helps maintain the expression of the epithelial microRNA *miR-200* ^129^. This establishes an incoherent feed-forward loop from *NF-κB* to *miR-200*, allowing *miR-200* to maintain its expression in hybrid E/M cells.
2. <u>*EMT switch*</u> (yellow nodes on **Fig. 1C, S3**) is a tri-stable module that locks the cell into an epithelial, mesenchymal or hybrid state. At its core, the model includes a cascading series of positive feedback loops that leverage mutual inhibition between the epithelial microRNAs *miR-34* and *miR-200*, and the EMT transcription factors *Snai1* and *Zeb1* ^112,130–136^. In our model, *miR-34* is active unless repressed by either *Snai1* or *Zeb1*, while *miR-200* can be maintained by *p21* activity which binds to and interferes with *Zeb1*’s ability to repress it ^133^. In its absence, *Snai1, Twist*, and *Zeb1* are required to repress it ^112,134–136^. Our model assumes that high levels of *Zeb1 (Zeb1_H* = ON) can override *c-Myb* and/or shut down the actively transcribed *miR-200* locus, while moderate *Zeb1* activity can maintain epigenetic silencing in the absence of *c-Myb* or ongoing transcription ^137^. The EMT switch is flipped from its epithelial state by the cooperative effect of growth signals and mechanosensitive inputs. Namely, mitogenic signaling through *PI3K /AKT* turns off the *Snai1* inhibitor *GSK3* ^75^ and activates its transcriptional inducer *NF-κB* ^76^. In addition, *Pak1* activation in horizontally polarized cells preparing to migrate induces nuclear localization of *Snai1* ^138^. *Snai1*, in turn, represses *miR-34*, activates *Twist* ^139^ and indirectly, *Snai2* (via *Twist*) ^140^. *Snai2*, in turn, increases transcription of *Zeb1* ^141^, though to moderate levels seen in hybrid E/M cells in which *miR-200* is not yet repressed. Flipping of the *miR-34* ⊢ ⊣ *Snai1* part of the tri-stable EMT switch thus leads to partial EMT to a hybrid E/M state, where cells can migrate but still maintain adherens junctions. In the absence of exogenous signals such as *TGFβ*, the second step from hybrid E/M to a complete mesenchymal state and full loss of *E-cadherin* expression requires high nuclear *β-catenin*, which contributes to *Zeb1* and *Snail2* (*Slug*) activation and is required for high *Zeb1* activity ^142–144^. Here we assume that this requires at least partial disassembly of adherens junctions to release *β-catenin* to the cytosol ^145^ (in addition to repressed *miR-34* ^23^), either direct activation of *β-catenin* by *Smads* ^146,147^ or complete lack of adherens junctions ^145^, and loss of inhibition by either *miR-200* ^148^ or *GSK3* ^149^. High nuclear *β-catenin* and *LEF1* (induced by *Zeb1* as well as *Smad* or *NF-κB* ^150–152^) aid *Snai2* in inducing high levels of *Zeb1* ^65,141,144^, and subsequently repressing *miR-200* expression ^134,135^. This results in full *E-cadherin* repression and flips the EMT switch from a hybrid E/M to a mesenchymal state. Finally, high nuclear *β-catenin*, in complex with *TCF4*, induces the expression of secreted *TGFβ*, setting up an autocrine *TGFβ* signaling loop ^153^.
3. <u>*TGFβ signaling*</u> (bright red nodes on **Fig. S3**) tracks the activation of *TβRI / TβRII* by either externally supplied TGF*β* (*TGFb_ext* = ON), or the autocrine signal secreted by a mesenchymal cell (*TGFb_secr* = ON). To model varying strengths of autocrine signaling depending on cell type as well as the biophysical environment that modulates diffusion and/or receptor access to secreted TGF*β*, we assume that either moderate cell density (*CellDensity_Low* = ON) or a strong autocrine self-loop (*Self_Loop* = ON) are required for secreted TGF*β*’s effect. Active *TβRI*/ *TβRII*then phosphorylate and activate *Smad2/3* ^83^ (counteracted in dense monolayers by *Sprouty2* ^154^). *Smad2/3* partner with *Smad4* and Yap/TAZ to enter the nucleus ^83,155^ and hinder proliferation (induce *p21* ^156^, *p15* ^157^ and *4EBP1* ^158^, block *Myc* ^159^) and potentiate apoptosis (reduce *BCL-XL* ^160^ and *ERK* via *DUSP4* ^161,162^, induce *BIK* ^160^, and *BIM* via Runx1 ^163^). Critically, *Smad2/3/4* also induce EMT by inducing *LEF1* ^151^ and *Snai2* ^164^, by aiding nuclear accumulation of *β-catenin* ^146,147^, and by inducing *HMGA2* ^165^. The *HMGA2* transcription factor is then responsible for TGF*β*-induced *Snai1*, *Twist*, and *Snai2* expression ^165^. In addition to its canonical *Smad*-mediated effects, *TβRI / TβRII* also activate *Src* kinase, the *PI3K/AKT* pathway ^166,167^ (the latter aided by *Smad*-mediated *ILK/Rictor* complex formation ^73,168,169^), as well as MAPK signaling via *Shc/Grb2* recruitment ^170^. This paradoxical regulation of pathways that mediate both survival and proliferation render TGF*β* highly context dependent.

### QUANTIFICATION AND STATISTICAL ANALYSIS

#### Automated phenotype detection

The code package *ReganLabBooleanSims* uses an automated process to assign specific biological phenotypes to any state or time-course of the model and monitor phenotypic transitions ^45^; namely healthy vs. erroneous cell cycle progression, apoptosis, as well as contact inhibited, migratory, epithelial, mesenchymal and hybrid E/M states. The signatures of these biological states are user-defined as follows

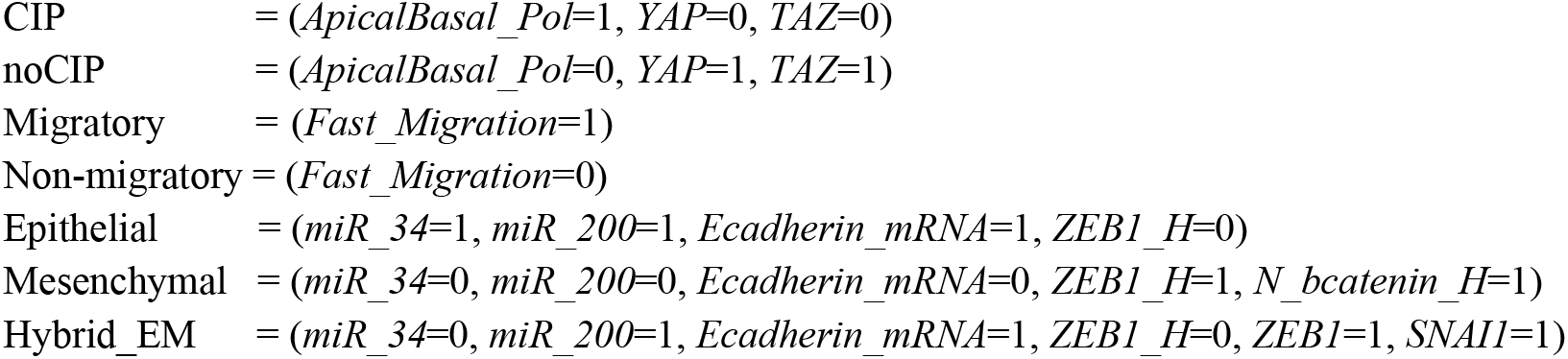

and read by *ReganLabBooleanSims* from a file named *VirtualExp_Phenotypes_for_STATS.txt* (**File S10**).

#### Phenotype statistics in non-saturating stochastic environments

To generate phenotype statistics in response to non-saturating environmental inputs, we ran time courses of *T*_stats_ = 50,000 time-steps in which each input node was randomly toggled ON/OFF in each time-step with an input-specific ON-probability. The simulations tracked the number of cell cycle events, marking cycles completed without error (**Fig. S1B**, *blue bars*), the number of genome duplication even from G2 (*orange bars*), the number of premature metaphase-anaphase transitions that did not involve completion of the mitotic spindle followed by genome duplication (*green bars*), and the number of genome duplication events in the absence of a cytokinesis step between telophase and the next S-phase (*red bars*). In addition, the simulation counted and stopped at each apoptotic event. Time courses that resulted in apoptosis before time *T*_stats_ were restarted until a minimum of *T*_stats_ steps of live-cell dynamics were sampled. In addition, the simulation tracked the average time spent in each reversible phenotype listed above by counting time-steps in which the model’s state satisfies the constraints that define each phenotype of interest. To generate phenotype statistics with incomplete knockdown or overexpression of target molecule (s), we combined the non-saturating stochastic inputs described above with a similar stochastic locking of the target molecule OFF or ON with a tunable probability *p*_KD_ (knockdown) or *p*_OE_ (over-expression), respectively. In time-steps where the molecule was not locked ON or OFF, it followed the internal Boolean regulatory influences of the rest of the network as if it was unperturbed.

All phenotype statistics are generated by *ReganLabBooleanSims* via *in silico* experiment(s) in *VirtualExp_Phenotypes_for_STATS.txt*, as illustrated below. Once the statistics are completed and written to file, *ReganLabBooleanSims* auto-generates the Python code required to show the results as stacked bar graphs such as **Figure S1B-D**. The simplest of these runs generates results as a function of a single, increasing input variable:

~~~
-- Figure S1B; code generates a figure panel with bar graphs of all phenotype statistics at increasing GF_High from 0% to 100% (x axis).
**NonSaturating_Stats_Scan_1Env_fnKDOE** 34 GF_High (Stiff_ECM=0.9, GF=1,
GF_High=0, CellDensity_Low=0, CellDensity_High=0)
**NonSaturating_Stats_Scan_1Env_fnKDOE** 34 Stiff_ECM (Stiff_ECM=0, GF=1,
GF_High=0.9, CellDensity_Low=0, CellDensity_High=0)
-- Figure S3B
**AsyncBias_NonSaturating_Stats_Scan_1Env_fnKDOE** 34 GF_High (Stiff_ECM=0.9,
GF=1, GF_High=0, CellDensity_Low=0, CellDensity_High=0)
**AsyncBias_NonSaturating_Stats_Scan_1Env_fnKDOE** 34 Stiff_ECM (Stiff_ECM=0,
GF=1, GF_High=0.9, CellDensity_Low=0, CellDensity_High=0)
~~~

Revealing the effects of a partial knock-out often requires choosing the right environment where the effect can manifest. To this end, adding an additional (*J_Ecadherin* KD) parenthesis at the end of **NonSaturating_Stats_Scan_1Env_fnKDOE** commands generates a different output. Namely, *ReganLabBooleanSims* generates a grid of bar graphs, all as a function of increasing knockdown. The horizontal direction along the grid shows results for increasing input settings (e.g., *GF_High* exposures from 5% to 95% below). The assayed phenotypes are organized vertically.

~~~
-- Figure S1D and S3D; phenotype statistics as a function of J_Ecadherin KD/OE (x axis of plots), for increasing GF_High exposures from 5% to 95% **NonSaturating_Stats_Scan_1Env_fnKDOE** 34 GF_High (Stiff_ECM=0.95, GF=1, GF_High=1, CellDensity_Low=1, CellDensity_High=0) (J_Ecadherin KD)
**AsyncBias_NonSaturating_Stats_Scan_1Env_fnKDOE** 34 GF_High (Stiff_ECM=0.95, GF=1, GF_High=1, CellDensity_Low=1, CellDensity_High=0) (J_Ecadherin KD)
~~~

Finally, *ReganLabBooleanSims* can generate results two increasing, non-saturating environmental inputs (e.g., *Stiff_ECM* and *CellDensity_Low*) as a 2D grid of bar graphs (1 page for each phenotype). The *x* axis within each bar graph corresponds to increasing KD/OE of the node in the last parenthesis (e.g., *Self_Loop*). All other inputs are dictated by the ON-probabilities indicated in the first parenthesis; the starting state is the attractor ID (e.g., 146).

~~~
-- Figure S20 data; 2D grid of bar graphs as a function of KD/OE
**NonSaturating_Stats_Scan_1KDOE_2Env** 146 Stiff_ECM CellDensity_Low
(Stiff_ECM=0, GF=1, GF_High=0.95, CellDensity_Low=0, CellDensity_High=0, TGFb_ext=0, Self_Loop=1) (Self_Loop KD)
~~~

### KEY RESOURCES TABLE

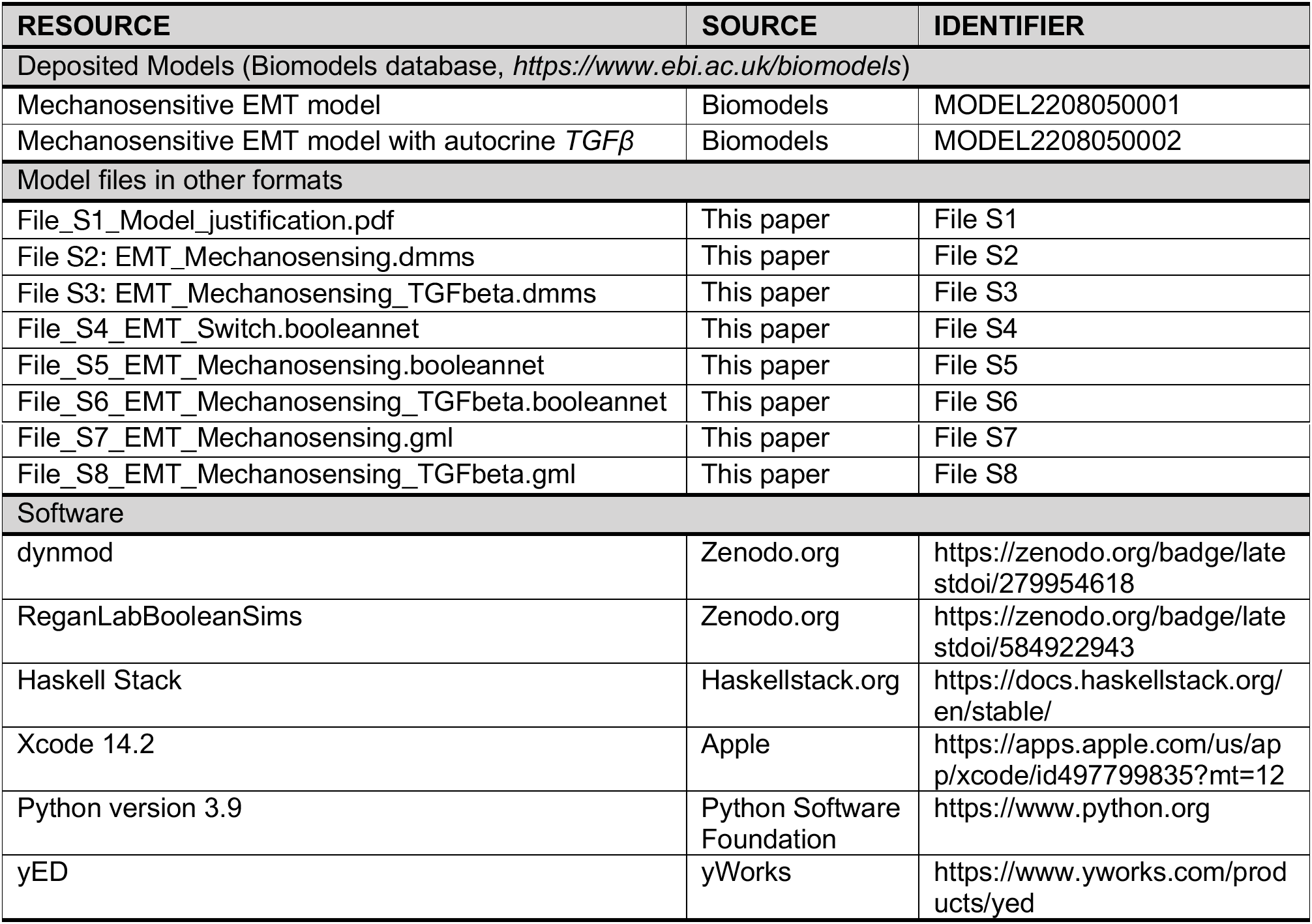

| RESOURCE | SOURCE | IDENTIFIER |
| --- | --- | --- |
| Deposited Models (Biomodels database, https://www.ebi.ac.uk/biomodels) |  |  |
| Mechanosensitive EMT model | Biomodels | MODEL2208050001 |
| Mechanosensitive EMT model with autocrine *TGFβ* | Biomodels | MODEL2208050002 |
| Model files in other formats |  |  |
| File_S1_Model_justification.pdf | This paper | File S1 |
| File S2: EMT_Mechanosensing.dmms | This paper | File S2 |
| File S3: EMT_Mechanosensing_TGFbeta.dmms | This paper | File S3 |
| File_S4_EMT_Switch.booleannet | This paper | File S4 |
| File_S5_EMT_Mechanosensing.booleannet | This paper | File S5 |
| File_S6_EMT_Mechanosensing_TGFbeta.booleannet | This paper | File S6 |
| File_S7_EMT_Mechanosensing.gml | This paper | File S7 |
| File_S8_EMT_Mechanosensing_TGFbeta.gml | This paper | File S8 |
| Software |  |  |
| dynmod | Zenodo.org | https://zenodo.org/badge/latestdoi/279954618 |
| ReganLabBooleanSims | Zenodo.org | https://zenodo.org/badge/latestdoi/584922943 |
| Haskell Stack | Haskellstack.org | https://docs.haskellstack.org/en/stable/ |
| Xcode 14.2 | Apple | https://apps.apple.com/us/app/xcode/id497799835?mt=12 |
| Python version 3.9 | Python Software Foundation | https://www.python.org |
| yED | yWorks | https://www.yworks.com/products/yed |

