## Supplemental Figures S1-S20 for "Boolean modeling of mechanosensitive Epithelial to Mesenchymal Transition and its reversal"

**Supplementary Figures**

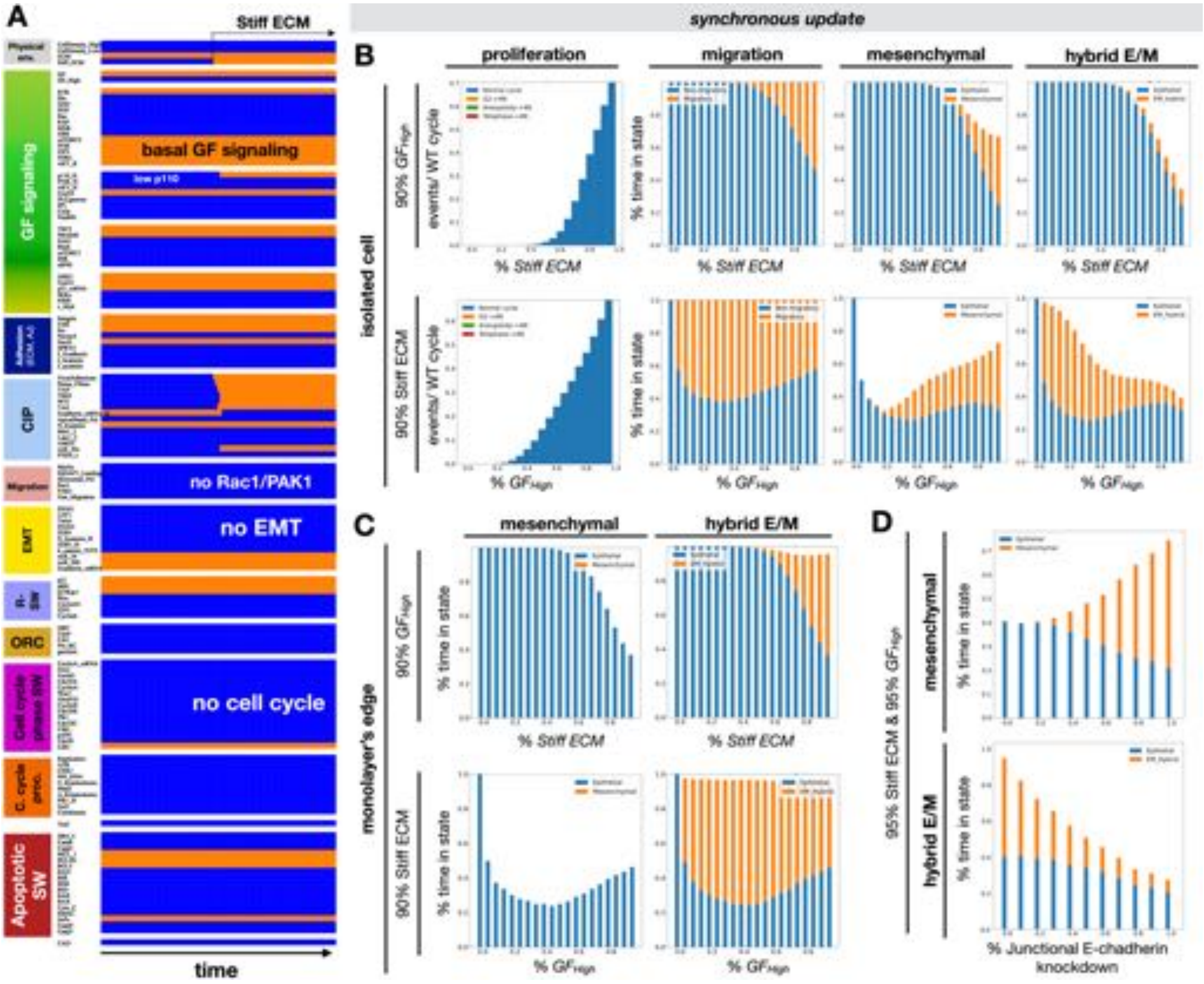


**Figure S1. Stiff ECM, strong growth signals, and loss of adherens junctions are all required for mechanically induced EMT; related to Figure 2. A)** Synchronous dynamics of regulatory molecule expression/activity during exposure of isolated, epithelial cells in low growth conditions to stiff ECM. *X-axis*: time-steps; *y-axis:* nodes organized in regulatory modules; *orange/blue:* ON/OFF; *black/white labels*: relevant molecular patterns. **B)** *Top row:* response of isolated cells to increasing *Stiff ECM* exposure in the presence of 90% saturating growth stimuli. *Top leftmost graphs:* Rate of normal cell cycle completion (*blue*), G2 → G1 reset followed by genome duplication (*orange*, not observed), aberrant mitosis followed by genome duplication (*green*, not observed), and failed cytokinesis followed by genome duplication (*red*, not observed) relative to the minimum cell cycle length (21 time-steps), shown as stacked bar charts. *Top row, 3 rightmost graphs:* fraction of time spent in a *i)* migratory (*orange*) vs. non-migratory (*blue*) state, *ii)* mesenchymal (*orange*) vs. epithelial (*blue*) state, and iii) hybrid E/M (*orange*) vs. epithelial (*blue*) state. *Bottom row:* parallel measurements as a function of increasing growth factor exposure on 90% stiff ECM. **C)** *Top/bottom:* response of cells at a monolayer's edge to increasing *Stiff ECM* exposure in the presence of 90% saturating growth stimuli (*top*) or increasing growth factor exposure on 90% stiff ECM (*bottom*). *Left/right:* fraction of time spent in a mesenchymal (*orange*) vs. epithelial (*blue*) state (*left*), or hybrid E/M (*orange*) vs. epithelial (*blue*) state (*right*). **D)** *Top/bottom:* fraction of time spent in a mesenchymal (*orange*) vs. epithelial (*blue*) state (*top*), or hybrid E/M (*orange*) vs. epithelial (*blue*) state (*bottom*) as a function of increasing *junctional E-cadherin* inhibition at a monolayer's edge, 95% *Stiff ECM* and 95% saturating growth stimuli. *Total sampled live cell time:* 100,000 steps; synchronous update; *Initial state for sampling runs:* isolated epithelial cell in low mitogens on a soft ECM.

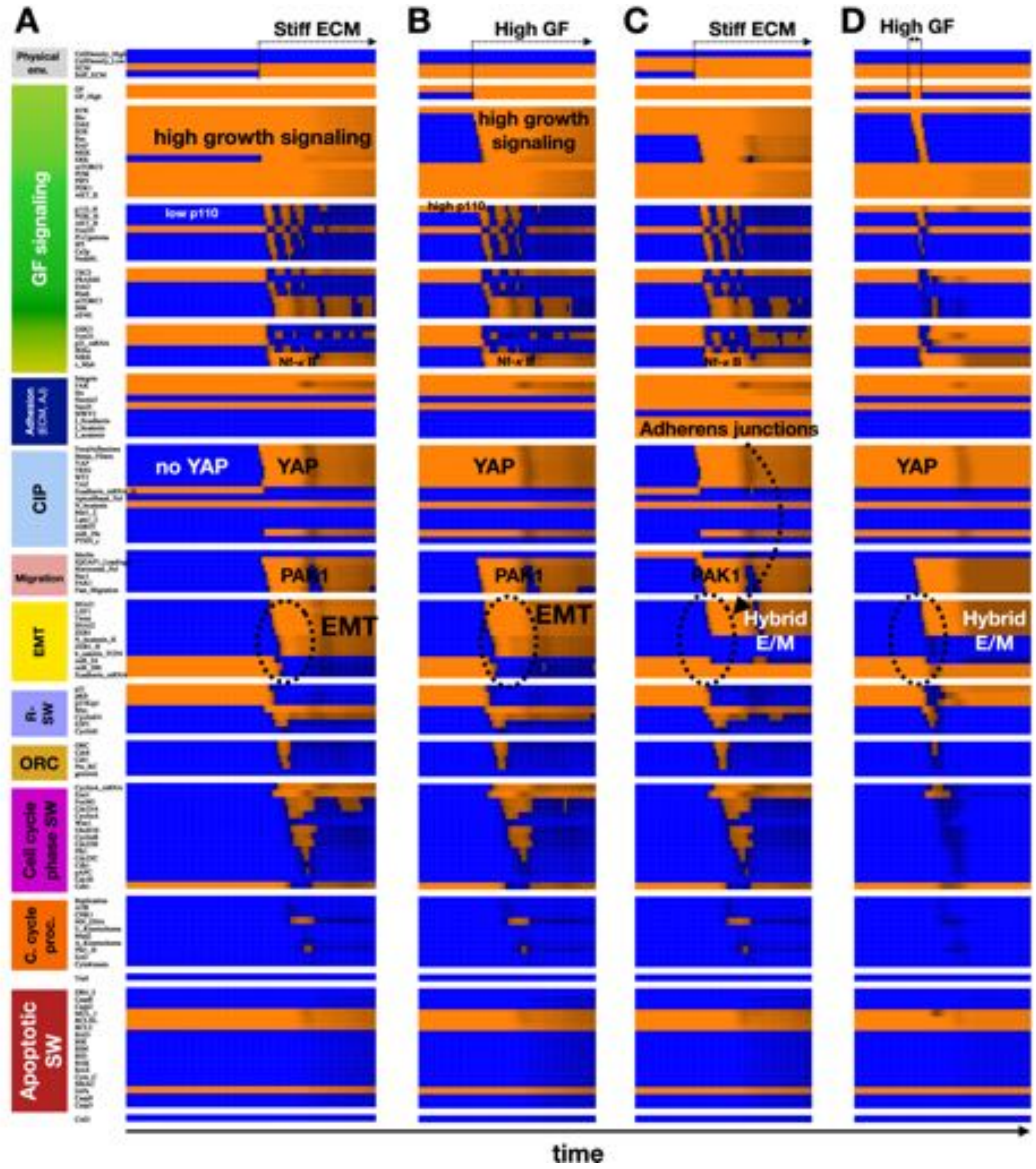


**Figure S2. Simulations reproducing Figure 2 with biased asynchronous update (biomechanically controlled EMT). A-D)** Biased asynchronous dynamics of regulatory molecule expression/activity during exposure of (A) isolated, growth-stimulated epithelial cells to stiff ECM, (B) isolated epithelial cells in low growth conditions on a stiff ECM to strong mitogens, (C) isolated, growth-stimulated epithelial cells at a monolayer edge to stiff ECM, and (D) isolated epithelial cells in low growth conditions on a stiff ECM to a brief (6-step) pulse of strong mitogens. *X-axis:* time-steps; *y-axis:* nodes organized in regulatory modules; *orange/black/blue color-scale:* average expression of each molecule across 1000 independent runs with biased asynchronous update (*orange* = all ON; *black* = 50% ON/OFF; *blue* = all OFF); *black/white labels & arrows:* molecular changes that drive EMT.

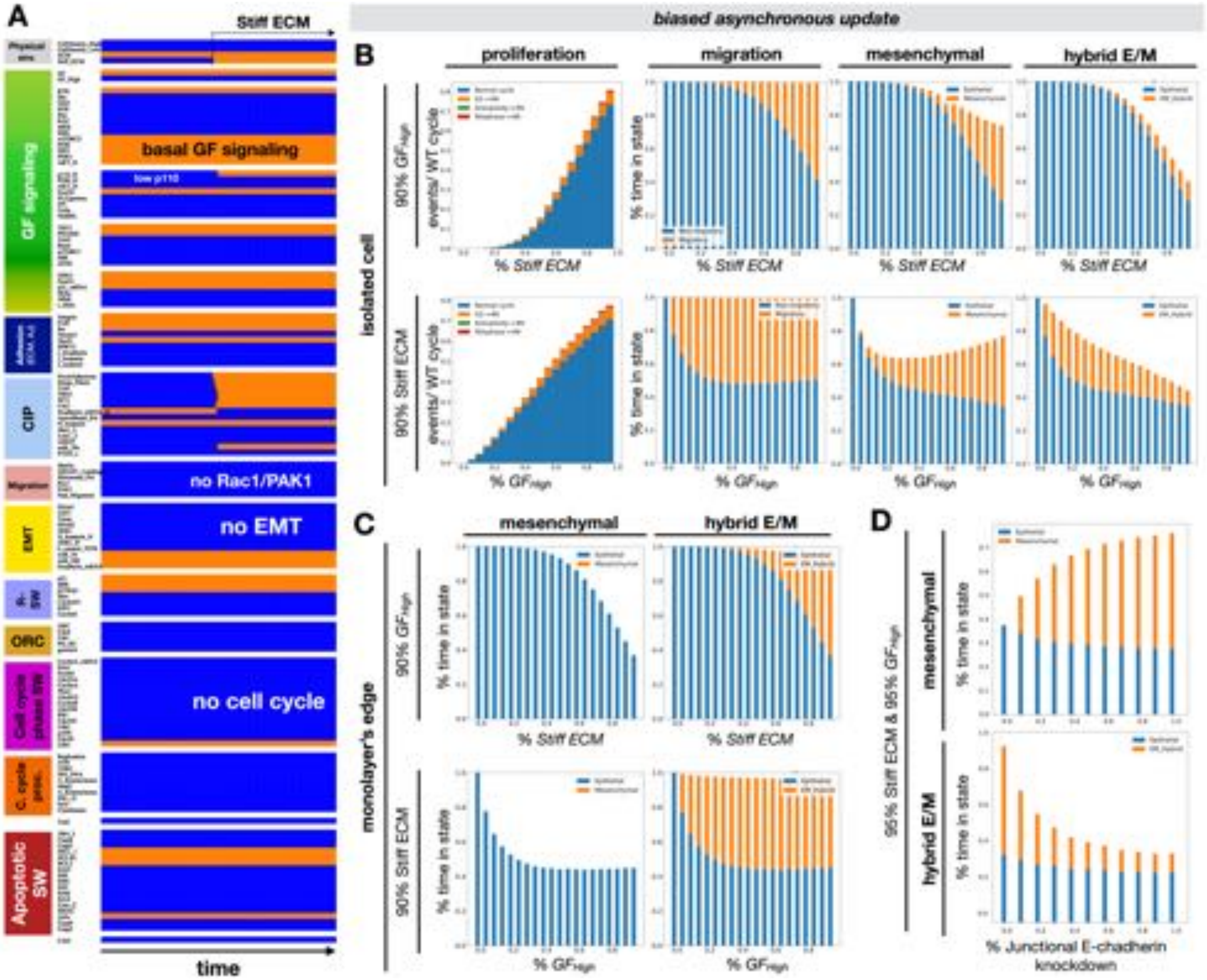


**Figure S3. Simulations reproducing Figure S1 with biased asynchronous update (joint requirements of biomechanically induced EMT); related to Figure 2. A)** Biased asynchronous dynamics of regulatory molecule expression/activity during exposure of isolated, epithelial cells in low growth conditions to stiff ECM. *X-axis*: time-steps; *y-axis:* nodes organized in regulatory modules; *orange/black/blue color-scale:* average expression of each molecule across 1000 independent runs with biased asynchronous update (*orange* = all ON; *black* = 50% ON/OFF; *blue* = all OFF); *black/white labels*: relevant molecular patterns. **B)** *Top row:* response of isolated cells to increasing *Stiff ECM* exposure in the presence of 90% saturating growth stimuli. *Top leftmost graphs:* Rate of normal cell cycle completion (*blue*), G2 → G1 reset followed by genome duplication (*orange*), aberrant mitosis followed by genome duplication (*green*, not observed), and failed cytokinesis followed by genome duplication (*red*) relative to the minimum synchronous cell cycle length (21 time-steps), shown as stacked bar charts. *Top row, 3 rightmost graphs:* fraction of time spent in a *i)* migratory (*orange*) vs. non-migratory (*blue*) state, *ii)* mesenchymal (*orange*) vs. epithelial (*blue*) state, and iii) hybrid E/M (*orange*) vs. epithelial (*blue*) state. *Bottom row:* parallel measurements as a function of increasing growth factor exposure on 90% stiff ECM. **C)** *Top/bottom:* response of cells at a monolayer's edge to increasing *Stiff ECM* exposure in the presence of 90% saturating growth stimuli (*top*) or increasing growth factor exposure on 90% stiff ECM (*bottom*). *Left/right:* fraction of time spent in a mesenchymal (*orange*) vs. epithelial (*blue*) state (*left*), or hybrid E/M (*orange*) vs. epithelial (*blue*) state (*right*). **D)** *Top/bottom:* fraction of time spent in a mesenchymal (*orange*) vs. epithelial (*blue*) state (*top*), or hybrid E/M (*orange*) vs. epithelial (*blue*) state (*bottom*) as a function of increasing *junctional E-cadherin* inhibition at a monolayer's edge, 95% *Stiff ECM* and 95% saturating growth stimuli. *Total sampled live cell time:* 100,000 steps; synchronous update; *Initial state for sampling runs:* isolated epithelial cell in low mitogens on a soft ECM.

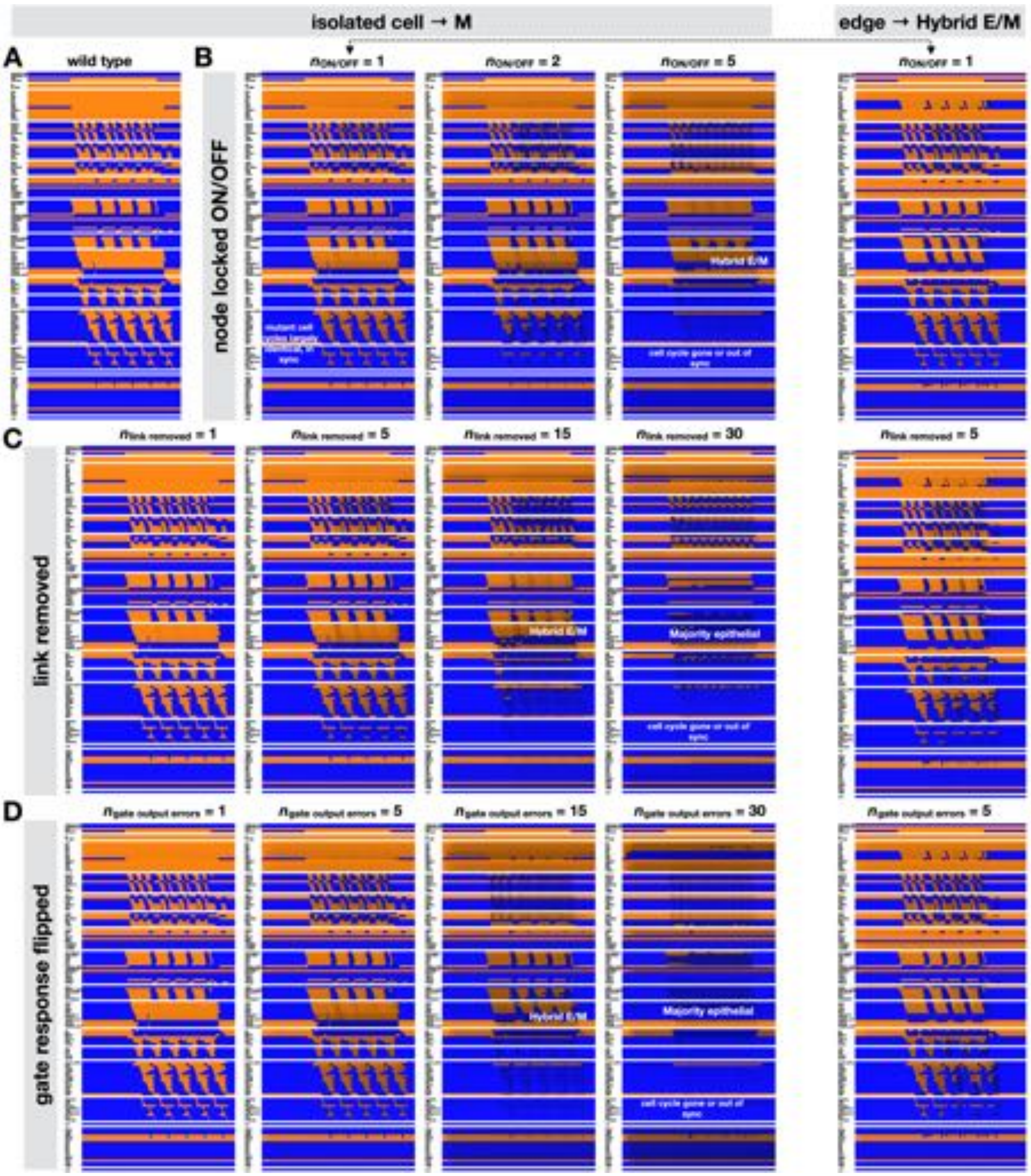


**Figure S4. Mechanosensitive EMT and MET are robust to random mutations / errors in model construction; related to Figure 2. A)** Synchronous dynamics of regulatory molecule expression/activity during exposure of an isolated, growth-stimulated epithelial cell to stiff ECM (wild-type model). **B-D)** *Leftmost panels:* simulation in (A) averaged over 1000 distinct mutant networks with (B) $n \in \{1,2,5\}$ random nodes per network locked ON or OFF, (C) $n \in \{1,5,15,30\}$ random links per network removed, and (D) $n \in \{1,5,15,30\}$ random gate outputs per network flipped. *Right column:* matching experiments starting with a cell at a monolayer's edge with (B) one locked node, (C) 5 links removed, or (D) 5 gate outputs flipped. *X-axis*: time-steps; *y-axis:* nodes organized in regulatory modules; *orange/black/blue color-scale:* average expression of each molecule across 1000 time-courses from independently generated mutant models (*orange* = all ON; *black* = 50% ON/OFF; *blue* = all OFF; synchronous update); *black/white labels*: relevant molecular patterns.

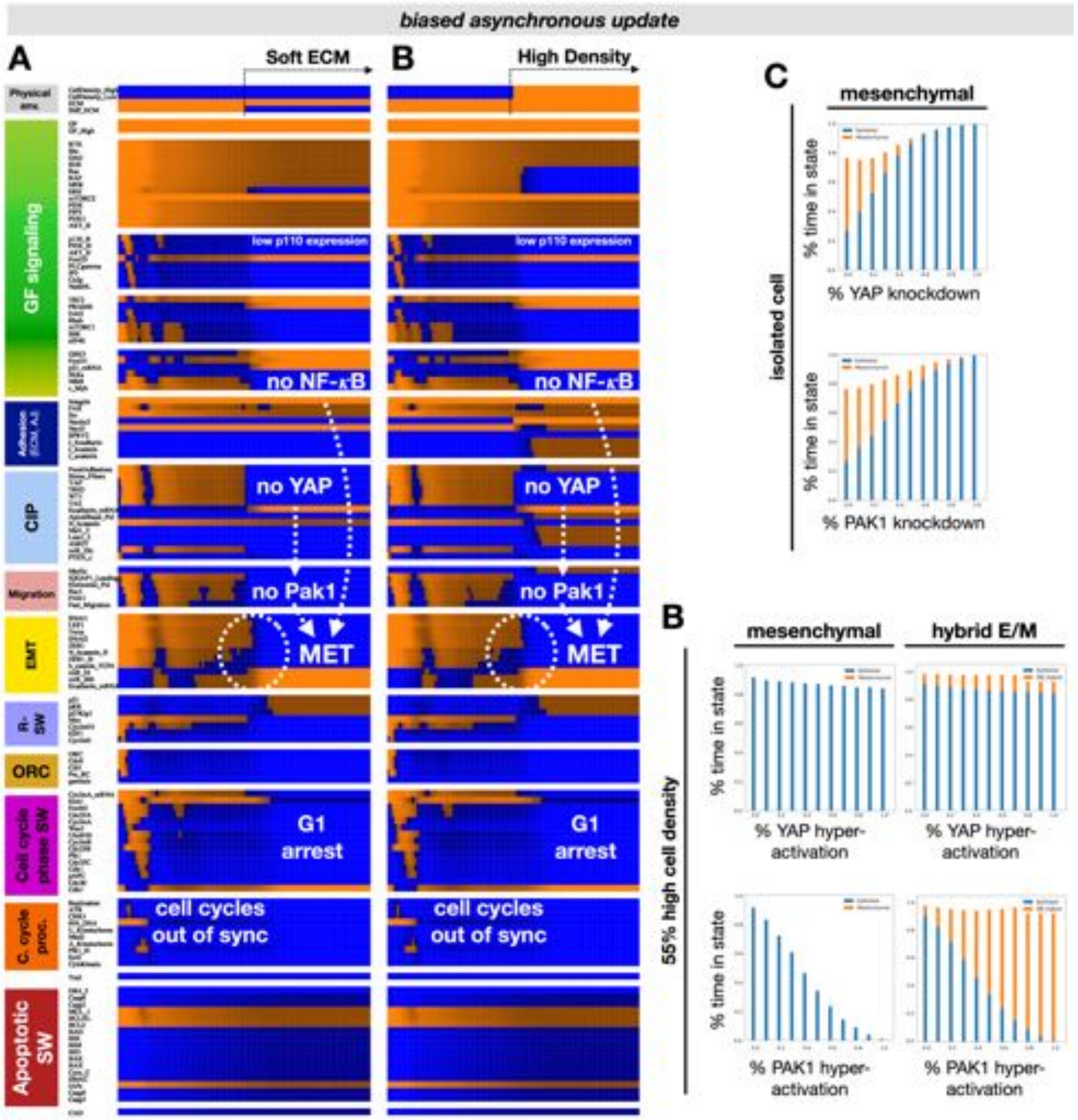


**Figure S5. Simulations reproducing Figure 3 with biased asynchronous update (MET in cells unable to spread). A-B)** Biased asynchronous dynamics of regulatory molecule expression/activity during exposure of an isolated, growth-stimulated and proliferating mesenchymal cell to (A) soft ECM, and (B) very high cell density. *X-axis:* time-steps; *y-axis:* nodes organized in regulatory modules; *orange/black/blue color-scale:* average expression of each molecule across 1000 independent runs with biased asynchronous update (*orange* = all ON; *black* = 50% ON/OFF; *blue* = all OFF); *black/white labels & arrows:* molecular changes that drive MET. **C)** *Top/bottom:* fraction of time spent in a mesenchymal (*orange*) vs. epithelial (*blue*) state as a function of increasing *YAP1* (*top*) or *PAK1* (*bottom*) inhibition in isolated cells on 95% *Stiff ECM* and exposed to 95% saturating growth stimuli. **D)** *Top/bottom:* fraction of time spent in *i*) mesenchymal (*orange*) vs. epithelial (*blue*) state (*left*) or *ii*) hybrid E/M (*orange*) vs. epithelial (*blue*) state (*right*) as a function of increasing *YAP1* (*top*) or *PAK1* (*bottom*) inhibition in cells at 55% high density (no free space to spread 55% of the time) on 95% *Stiff ECM* and exposed to 95% saturating growth stimuli. *Total sampled live cell time:* 100,000 steps; synchronous update; *Initial state for sampling runs:* isolated epithelial cell in low mitogens on a soft ECM.

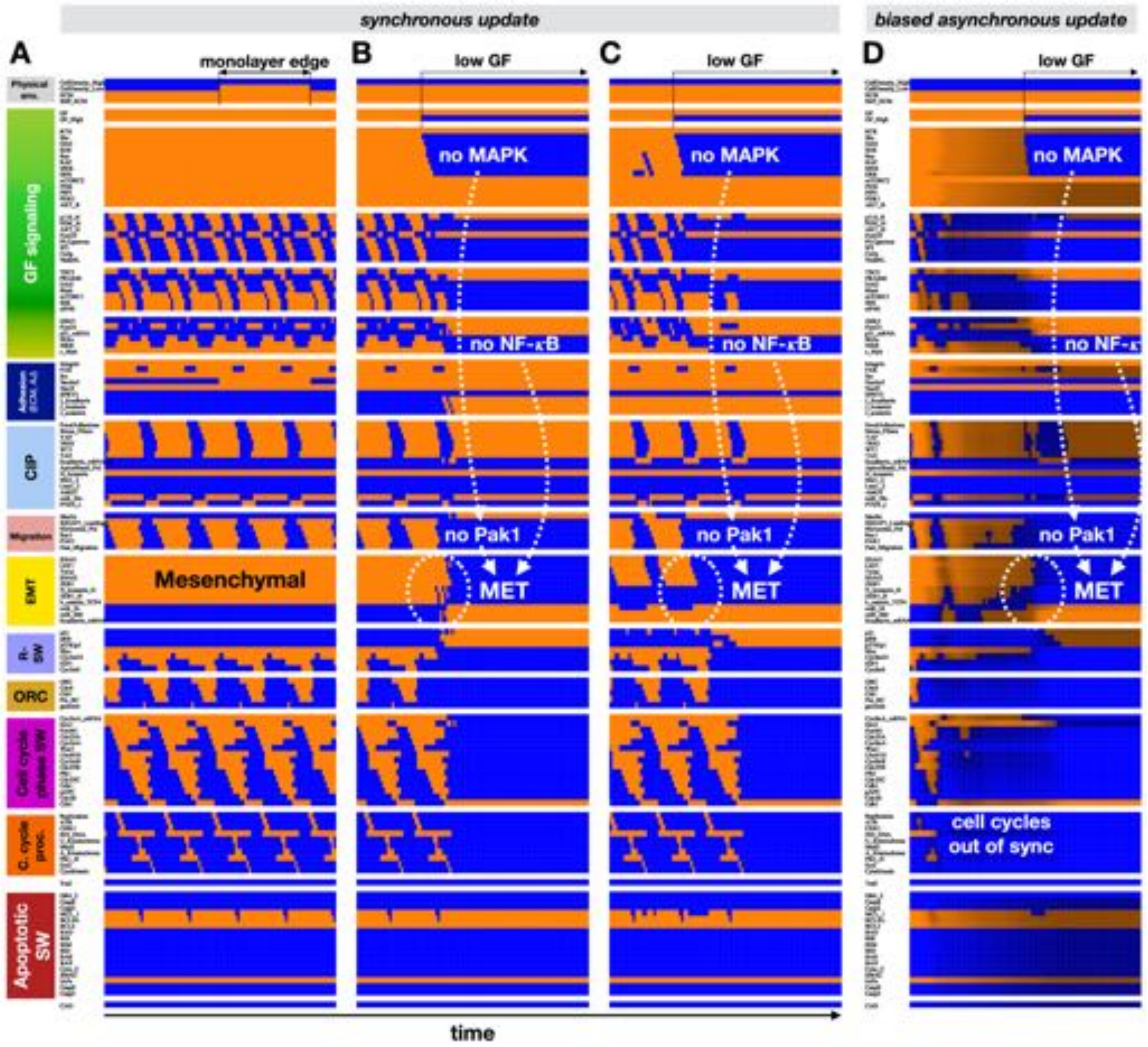


**Figure S6. Mitogen withdrawal triggers MET in proliferating mesenchymal and hybrid E/M cells without autocrine *TGFβ* signaling; related to Figure 3. A-C)** Synchronous dynamics of regulatory molecule expression/activity during exposure of (A) isolated, growth-stimulated mesenchymal cells to densities akin to a monolayer's edge, (B) dividing mesenchymal cells at a monolayer's edge to a decrease in growth signals, and (C) dividing hybrid E/M cells at a monolayer's edge to a decrease in growth signals. **D)** Biased asynchronous dynamics of regulatory molecule expression/activity during exposure of dividing mesenchymal cells at a monolayer's edge to a decrease in growth signals. *X-axis:* time-steps; *y-axis:* nodes organized in regulatory modules; *orange/black/blue color-scale:* average expression of each molecule across 1000 independent runs with biased asynchronous update (*orange* = all ON; *black* = 50% ON/OFF; *blue* = all OFF); *black/white labels & arrows:* molecular changes that drive MET.

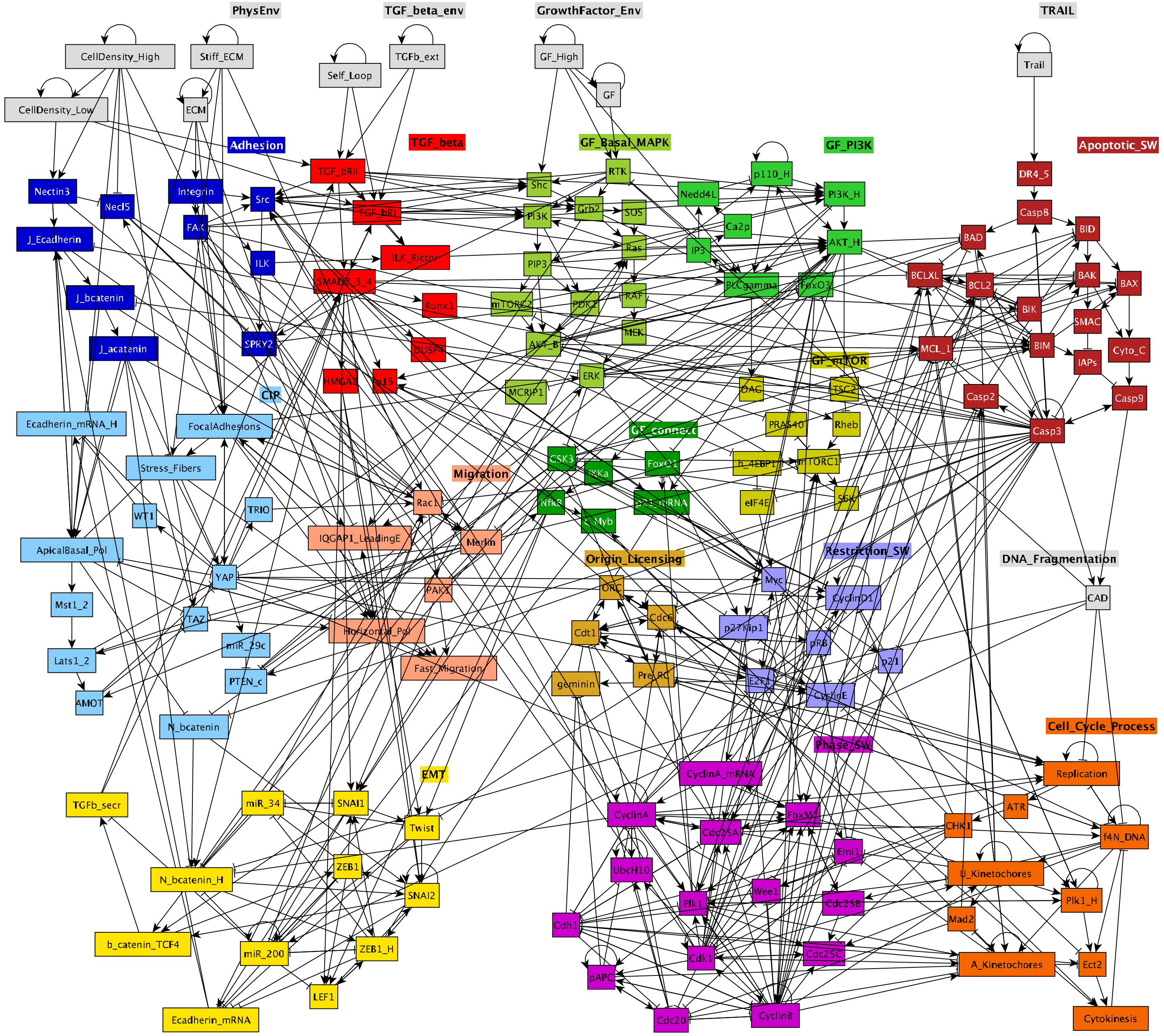


**Figure S7. Modular model of *TGFβ* signaling, EMT, growth factor signaling, cell cycle and apoptosis, adhesion, adherens junction formation, contact inhibition, and migration; related to Figure 4.** Modular network representation of our extended Boolean model. *Gray:* inputs representing environmental factors; *dark blue:* Adhesion signals; *red: TGFβ* signaling; *green*: Growth Signaling (*lime green*: basal AKT & MAPK, *bright green:* PI3K/AKT oscillations, *mustard*: mTORC1, *dark green: NF-κB, GSK3, FoxO1*); *dark red:* Apoptotic Switch*; light blue:* Contact Inhibition; *pink/light orange:* Migration; *light brown:* Origin of Replication Licensing; *lilac:* Restriction Switch; *purple:* Phase Switch; *dark orange:* cell cycle processes; *yellow:* EMT switch; → : activation; ⊣ : inhibition.

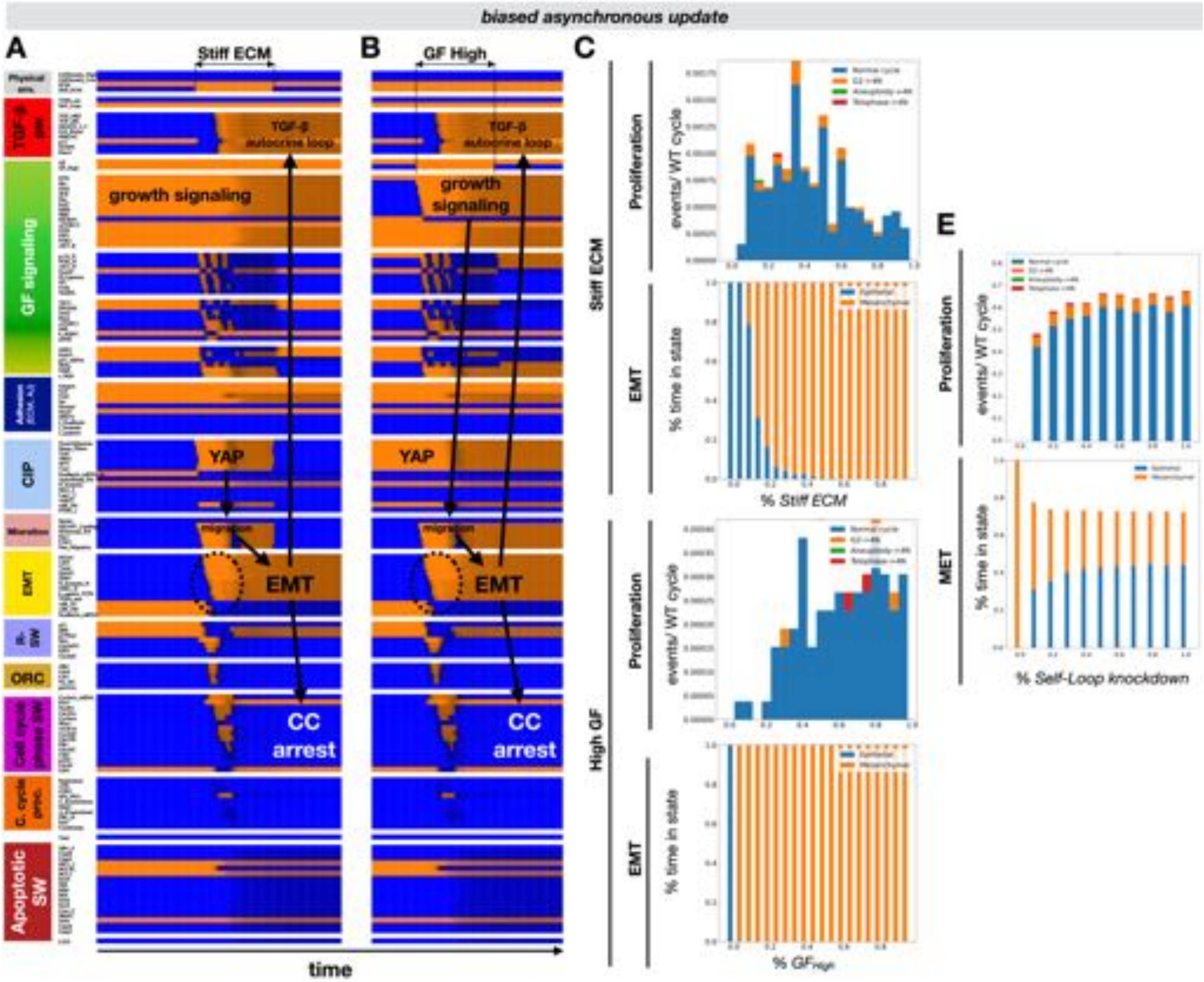


**Figure S8. Simulations reproducing Figure 4 with biased asynchronous update (EMT with autocrine *TGFβ*-induced cell cycle arrest). A, B)** Biased asynchronous dynamics of regulatory molecule expression/activity during exposure of (B) an isolated, growth-stimulated epithelial cell to stiff ECM, and (C) an isolated epithelial cell on stiff ECM to strong mitogens. *X-axis:* time-steps; *y-axis:* nodes organized in regulatory modules; *orange/black/blue color-scale:* average expression of each molecule across 1000 independent runs with biased asynchronous update (*orange* = all ON; *black* = 50% ON/OFF; *blue* = all OFF); *black/white labels & arrows:* molecular changes that drive EMT. **C)** *Top two vs. bottom 2 panels:* response of isolated cells to *i*) increasing *Stiff ECM* exposure in the presence of 95% saturating growth stimuli (*top two*) vs. ii) increasing growth factor exposure on 95% stiff ECM (*bottom two*). *Proliferation label:* rate of normal cell cycle completion (*blue*) vs. G2 → G1 reset (*orange*), aberrant mitosis (*green*, not observed), or failed cytokinesis (*red*) followed by genome duplication, relative to the minimum cell cycle length (21 time-steps), shown as stacked bar charts. *EMT label:* fraction of time spent in a mesenchymal (*orange*) vs. epithelial (*blue*) state. **D)** Response of isolated cells on 85% Stiff ECM exposed to 85% saturating growth stimuli to increasing inhibition of autocrine signaling (*Self-Loop* knockdown). *Top:* rate of normal cell cycle completion (*blue*) vs. G2 → G1 reset (*orange*), aberrant mitosis (*green*, not observed), or failed cytokinesis (*red*) followed by genome duplication. *Bottom:* fraction of time spent in a mesenchymal (*orange*) vs. epithelial (*blue*) state. *Total sampled live cell time:* 100,000 steps; synchronous update; *Initial state for sampling runs:* isolated epithelial cell in low mitogens on a soft ECM.

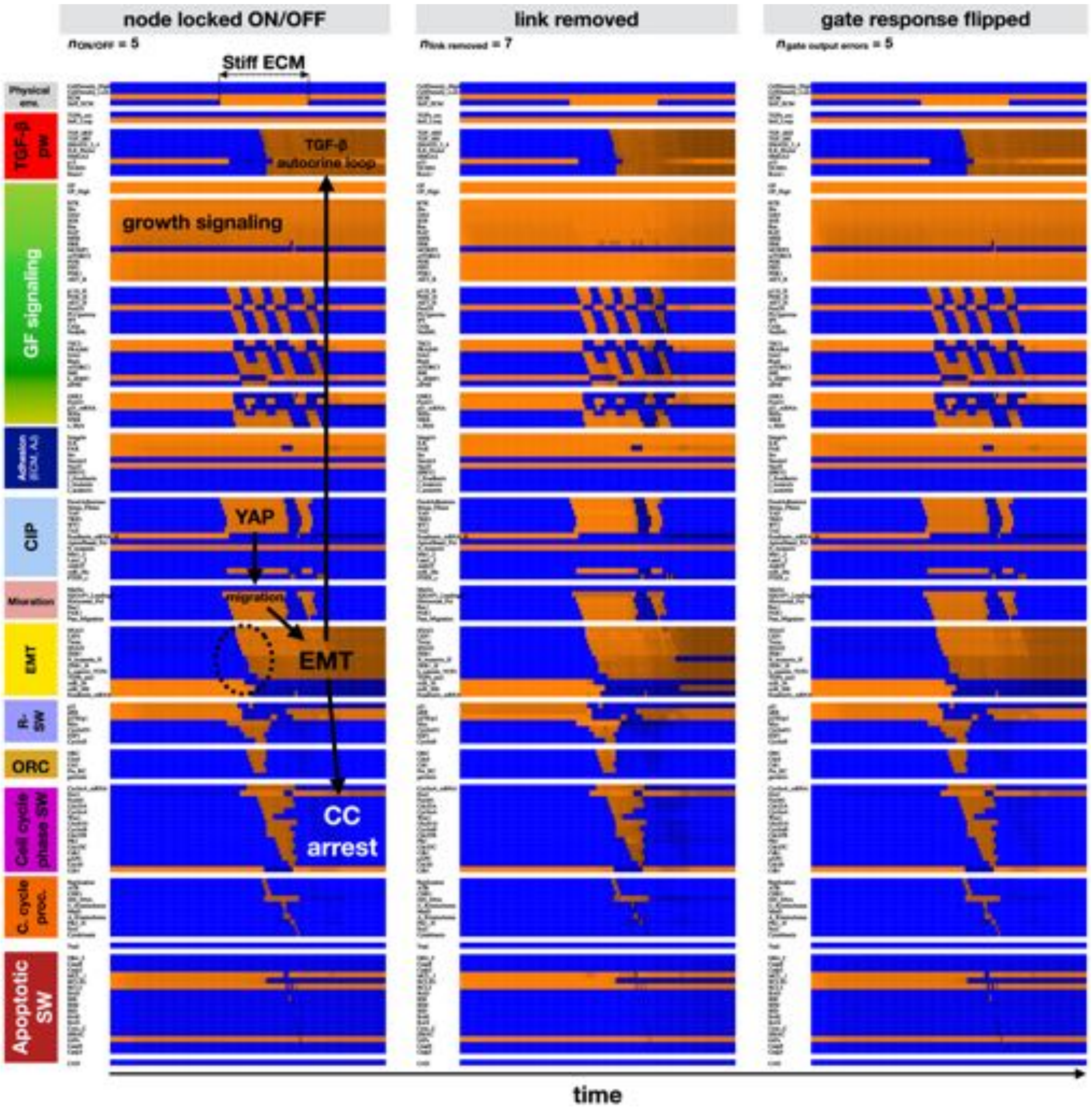


**Figure S9. Stabilization of EMT in response to stiff ECM by autocrine *TGFβ* is robust to random mutations / errors in model construction; related to Figure 4.** Average synchronous dynamics of regulatory molecule expression/activity during reversible exposure of an isolated, growth-stimulated epithelial cell to stiff ECM in an ensemble of mutant networks with 5 random nodes locked ON/OFF (*left*), 7 random links removed (*middle*), and 5 random gate outputs flipped per network (*right*). *X-axis*: time-steps; *y-axis:* nodes organized in regulatory modules; *orange/black/blue color-scale:* average expression of each molecule across 1000 time-courses from independently generated mutant models (*orange* = all ON; *black* = 50% ON/OFF; *blue* = all OFF; synchronous update); *black/white labels*: relevant molecular patterns.

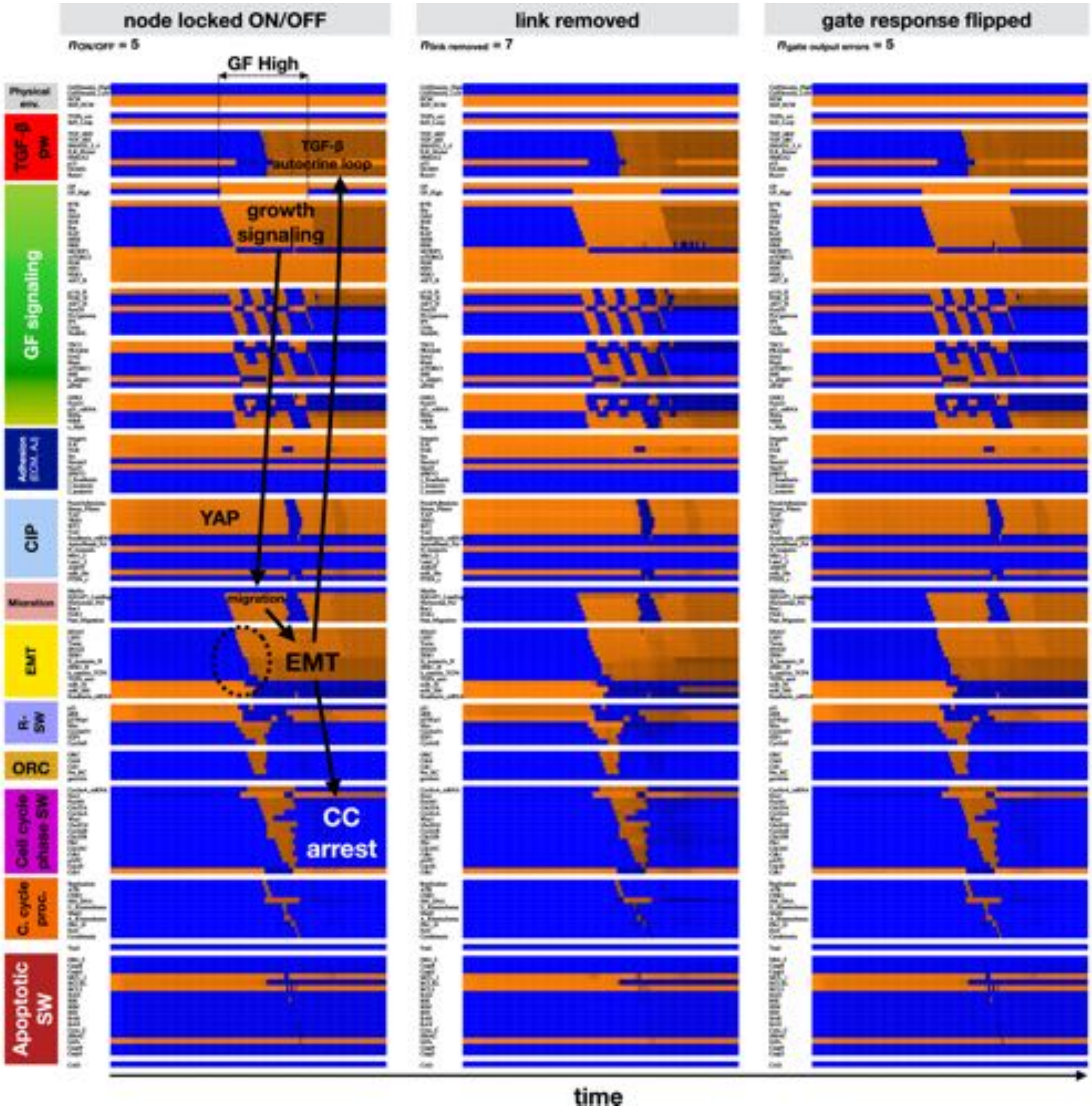


**Figure S10. Stabilization of EMT in response to growth factors by autocrine *TGFβ* is robust to random mutations / errors in model construction; related to Figure 4.** Average synchronous dynamics of regulatory molecule expression/activity during exposure of an isolated epithelial cell on a stiff ECM reversibly exposed to high levels of growth stimulus in an ensemble of mutant networks with 5 random nodes locked ON/OFF (*left*), 7 random links removed (*middle*), and 5 random gate outputs flipped per network (*right*). *X-axis*: time-steps; *y-axis:* nodes organized in regulatory modules; *orange/black/blue color-scale:* average expression of each molecule across 1000 time-courses from independently generated mutant models (*orange* = all ON; *black* = 50% ON/OFF; *blue* = all OFF; synchronous update); *black/white labels*: relevant molecular patterns.

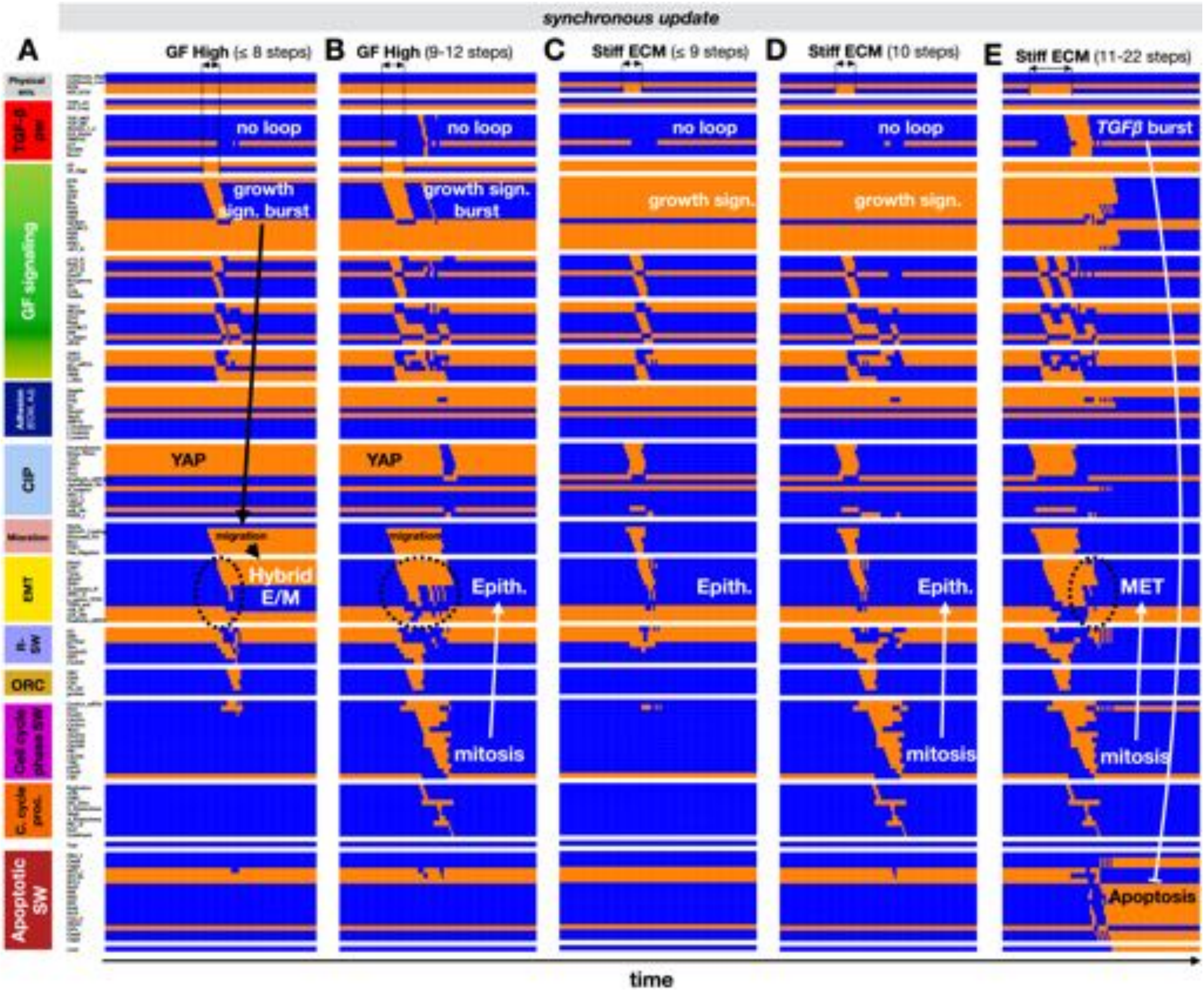


**Figure S11. In isolated cells with strong autocrine *TGFβ* signaling, short-lived growth signal bursts can trigger sustained hybrid E/M or cell cycle and no EMT, while exposure to stiff ECM that stops before cell cycle arrest can trigger apoptosis; related to Figure 4. A-B)** Synchronous dynamics of regulatory molecule expression/activity during exposure of isolated epithelial cells on stiff ECM to strong but brief growth signal bursts; (A): 8 timesteps leading to hybrid E/M; (B): 11 timesteps leading to a single division and reset to an epithelial state. **C-E)** Dynamics during exposure of isolated, growth-stimulated epithelial cells on soft ECM to stiff ECM for brief intervals; (C): 9 timesteps trigger no response; (D): 10 timesteps leading to a single division without EMT; (E): 22 timesteps leading to apoptosis. *X-axis:* time-steps; *y-axis:* nodes organized in regulatory modules; *orange/blue:* ON/OFF; *black/white labels & arrows:* molecular changes that drive MET.

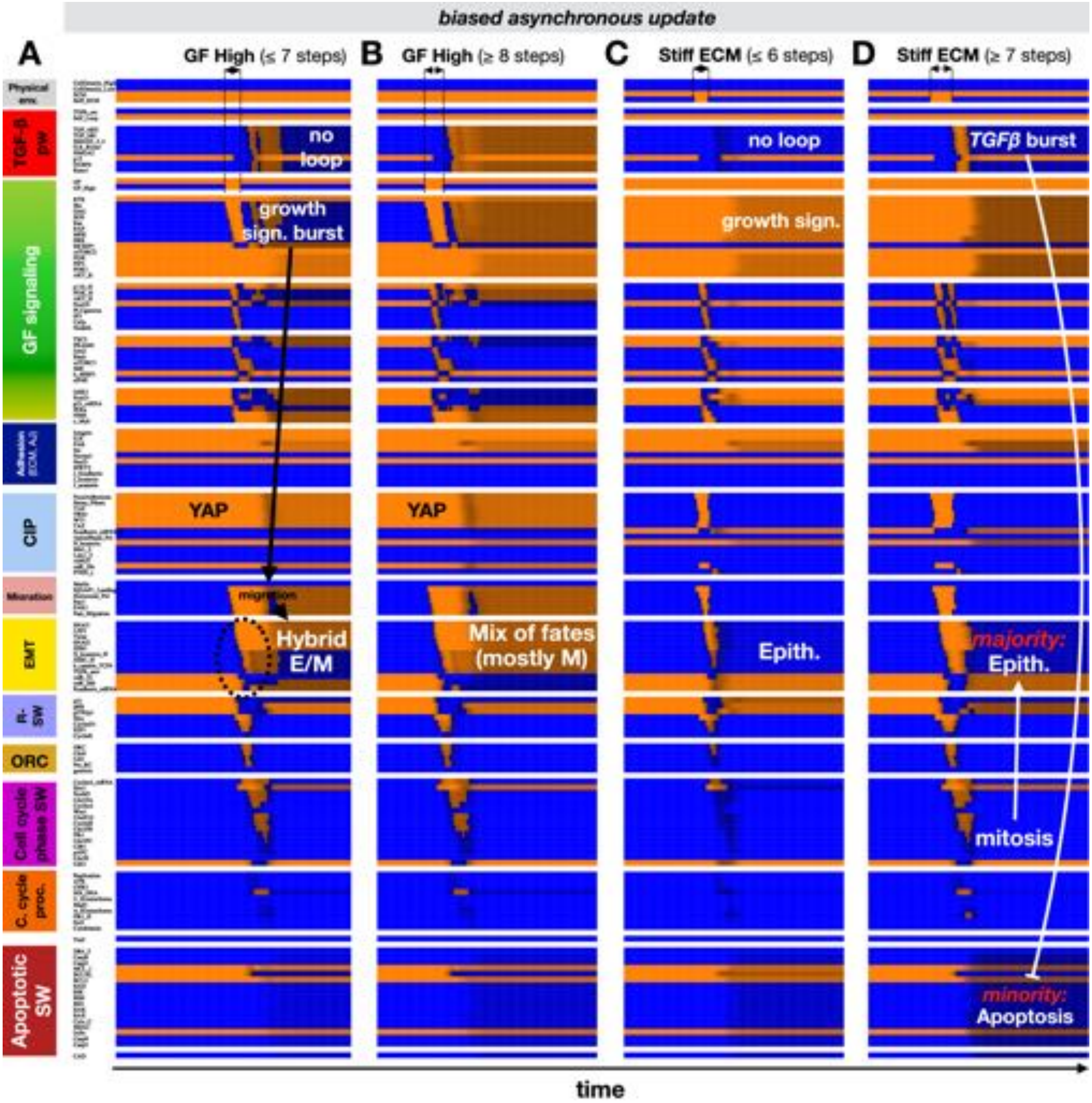


**Figure S12. Simulations reproducing Figure S11 with biased asynchronous update (short-lived growth signals or stiff ECM exposure); related to Figure 4. A-B)** Biased asynchronous dynamics of regulatory molecule expression/activity during exposure of isolated epithelial cells on stiff ECM to strong but brief growth signal bursts; (A): 7 timesteps leading to hybrid E/M; (B): 8 or more timesteps leading to a single division and a mix of fates, including mesenchymal and epithelial ones (dark orange/blue shading, towards black). **C-D)** Biased asynchronous dynamics during exposure of isolated, growth-stimulated epithelial cells on soft ECM to stiff ECM for brief intervals; (C): 6 timesteps trigger no response; (D): 7 or more timesteps lead to mix of fates, including a single division without EMT and apoptosis. *X-axis:* time-steps; *y-axis:* nodes organized in regulatory modules; *orange/black/blue color-scale:* average expression of each molecule across 1000 independent runs with biased asynchronous update (*orange* = all ON; *black* = 50% ON/OFF; *blue* = all OFF); *black/white labels & arrows:* molecular changes that drive MET.

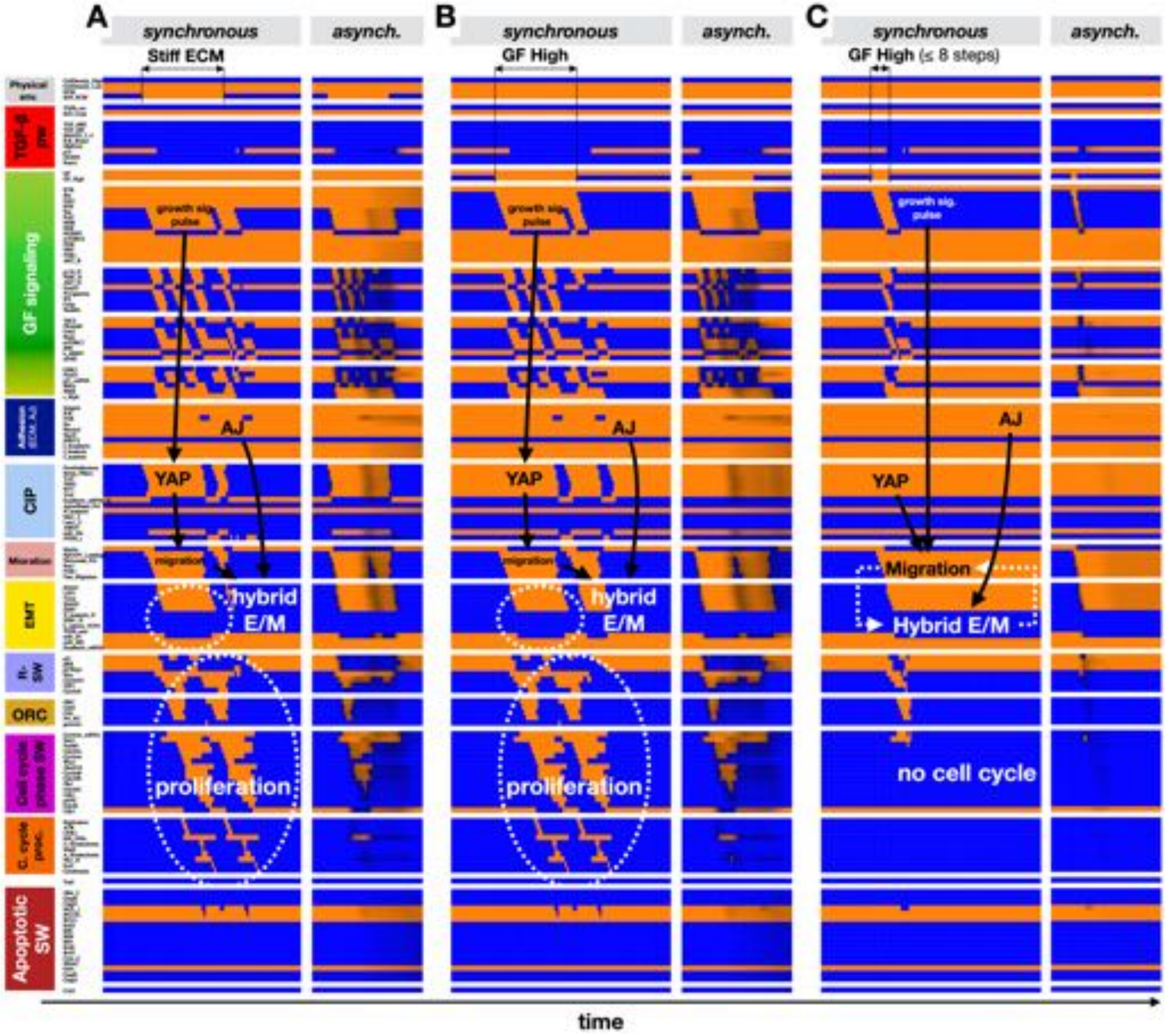


**Figure S13. In cells at a monolayer's edge, exposure to stiff ECM and prolonged mitogens both trigger partial EMT to a proliferative, reversible hybrid E/M state, while short-lived growth signals can trigger sustained hybrid E/M; related to Figure 4. A)** Synchronous (*left*) and biased asynchronous (*right*) dynamics of regulatory molecule expression/activity during exposure of growth-stimulated epithelial cells at a monolayer's edge on soft ECM to stiff ECM (40 timesteps), leading to reversible, proliferative hybrid E/M state. **B-C)** Synchronous (*left*) and biased asynchronous (*right*) dynamics during exposure of epithelial cells at a monolayer's edge on stiff ECM to strong growth signals; (B): 40 timesteps leading to reversible, proliferative hybrid E/M; (C): 8 synchronous (*left*) or 3 asynchronous (*right*) timesteps leading to a sustained hybrid E/M state. *X-axis:* time-steps; *y-axis:* nodes organized in regulatory modules; *orange/blue:* ON/OFF (*left*); *orange/black/blue color-scale:* average expression of each molecule across 1000 independent runs with biased asynchronous update (*orange* = all ON; *black* = 50% ON/OFF; *blue* = all OFF); *black/white labels & arrows:* molecular changes that drive EMT/MET.

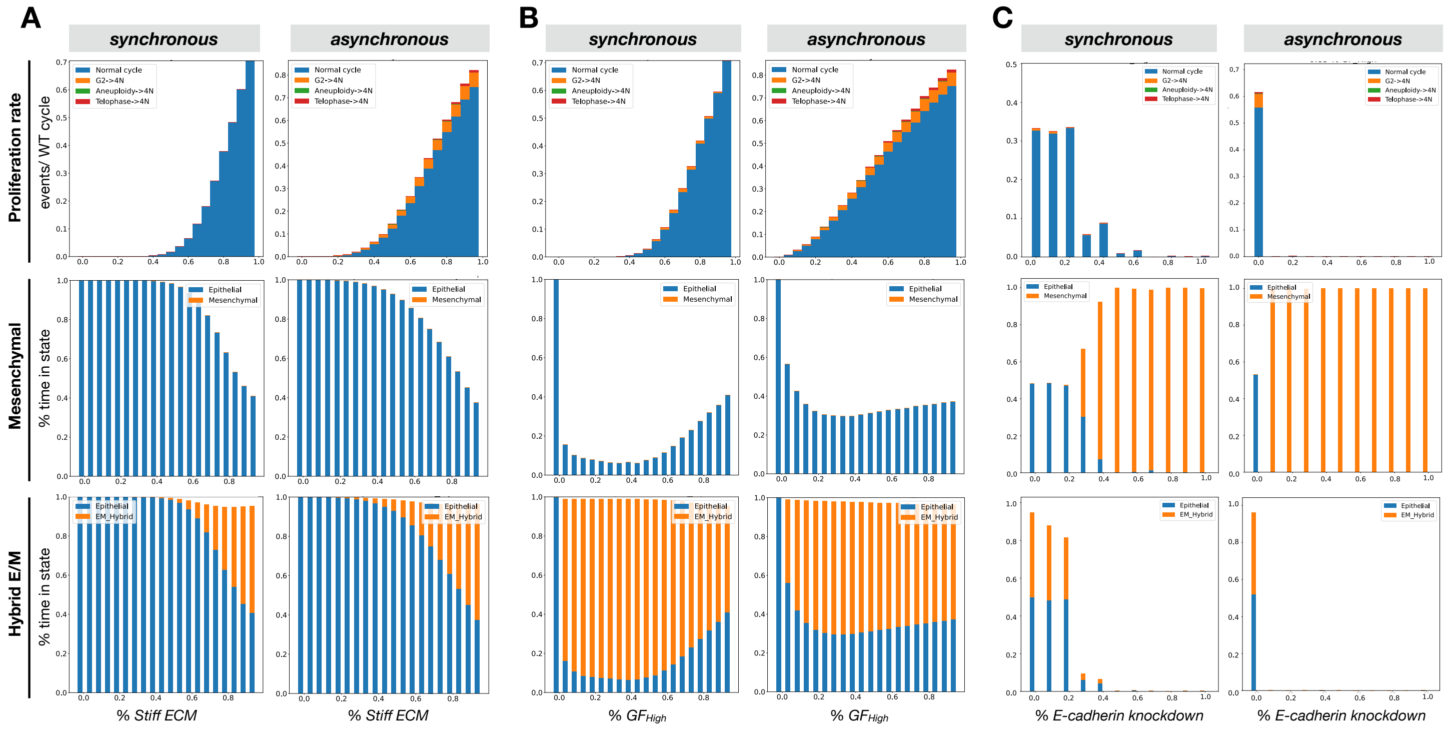


**Figure S14. In cells at a monolayer's edge, stiff ECM and mitogens promote proliferation and the hybrid E/M state, while knockdown of E-cadherin pushes cells to a mesenchymal state.** Synchronous (*left*) and biased asynchronous (*right*) response of cells at the edge of a monolayer cells to **A)** increasing *Stiff ECM* exposure in the presence of 95% saturating growth stimuli, **B)** increasing growth factor exposure on 95% stiff ECM, and **C)** increasing inhibition of *junctional E-cadherin* at 85% *Stiff ECM* and 85% saturating growth stimuli. *Left panels:* synchronous update; *right panels:* biased asynchronous update. *Top row (Proliferation):* rate of normal cell cycle completion (*blue*) vs. G2 → G1 reset (*orange*), aberrant mitosis (*green*), or failed cytokinesis (*red*) followed by genome duplication, relative to the minimum synchronous cell cycle length (21 time-steps), shown as stacked bar charts. *middle row (Mesenchymal):* fraction of time spent in a mesenchymal (*orange*) vs. epithelial (*blue*) state; *bottom row (Hybrid E/M):* fraction of time spent in a mesenchymal (*orange*) vs. epithelial (*blue*) state. *Total sampled live cell time:* 100,000 steps; synchronous update; *initial state for sampling runs:* isolated epithelial cell in low mitogens on a soft ECM.

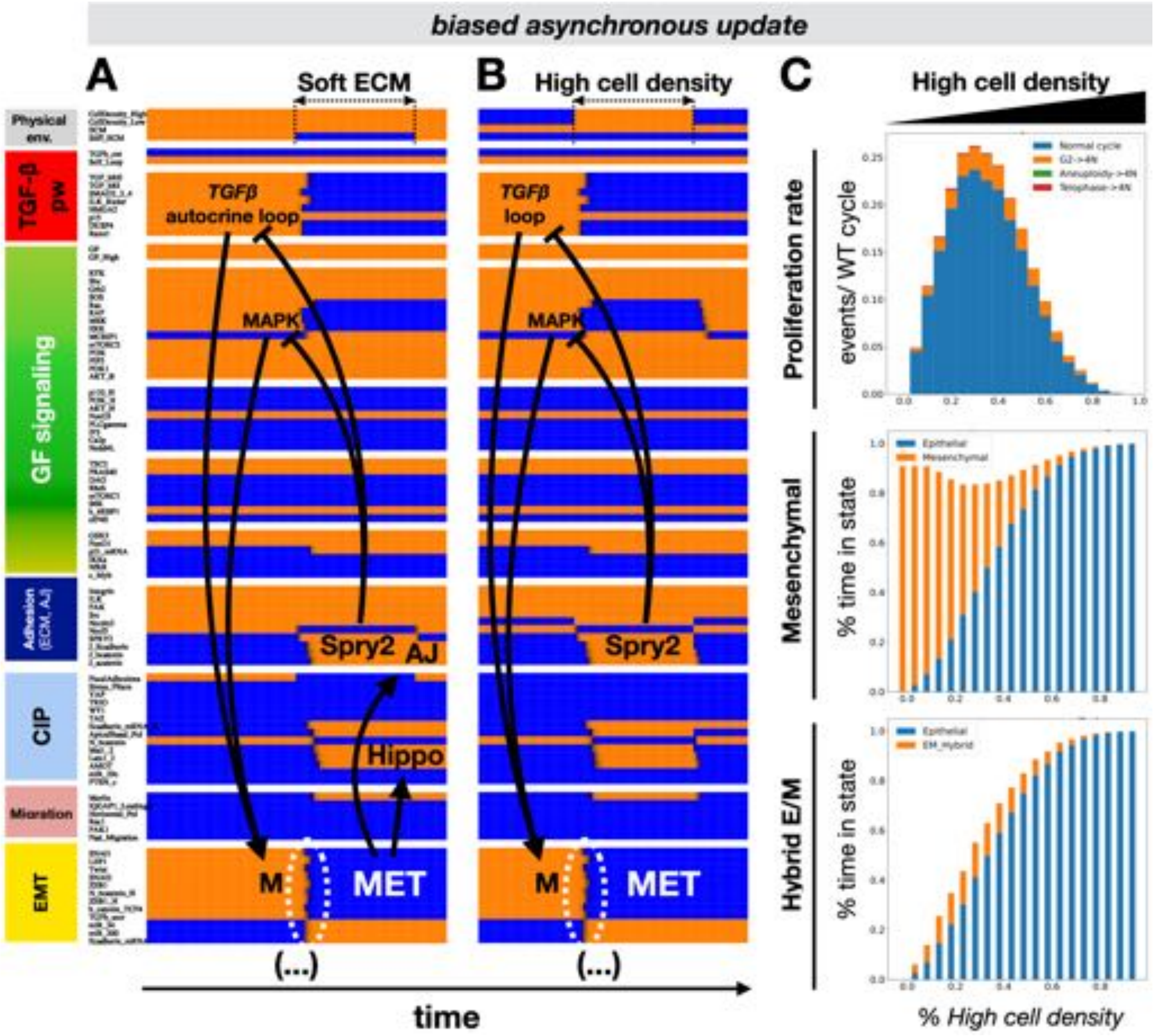


**Figure S15. Simulations reproducing Figure 5 with biased asynchronous update (MET at high density on soft ECM). A-B)** Biased asynchronous dynamics of regulatory molecule expression/activity during exposure of (A) a growth-stimulated mesenchymal cell on a stiff matrix but at very high density to a soft ECM, and (B) an isolated mesenchymal cell on soft ECM to high density. *X-axis:* time-steps; *y-axis:* nodes organized in regulatory modules; *orange/black/blue color-scale:* average expression of each molecule across 1000 independent runs with biased asynchronous update (*orange* = all ON; *black* = 50% ON/OFF; *blue* = all OFF); *black/white labels & arrows:* molecular changes that drive MET. **C)** Response of mesenchymal cells at the edge of a monolayer to increasing cell density in the presence of 95% saturating growth stimuli on 95% stiff ECM. *Top:* rate of normal cell cycle completion (*blue*) vs. G2 → G1 reset (*orange*), aberrant mitosis (*green*), or failed cytokinesis (*red*) followed by genome duplication, relative to the minimum cell cycle length (21 time-steps), shown as stacked bar charts. *Middle:* fraction of time spent in a mesenchymal (*orange*) vs. epithelial (*blue*) state. *Bottom:* fraction of time spent in a hybrid E/M (*orange*) vs. epithelial (*blue*) state. *Total sampled live cell time:* 100,000 steps; synchronous update; *Initial state for sampling runs:* isolated epithelial cell in low mitogens on a soft ECM.

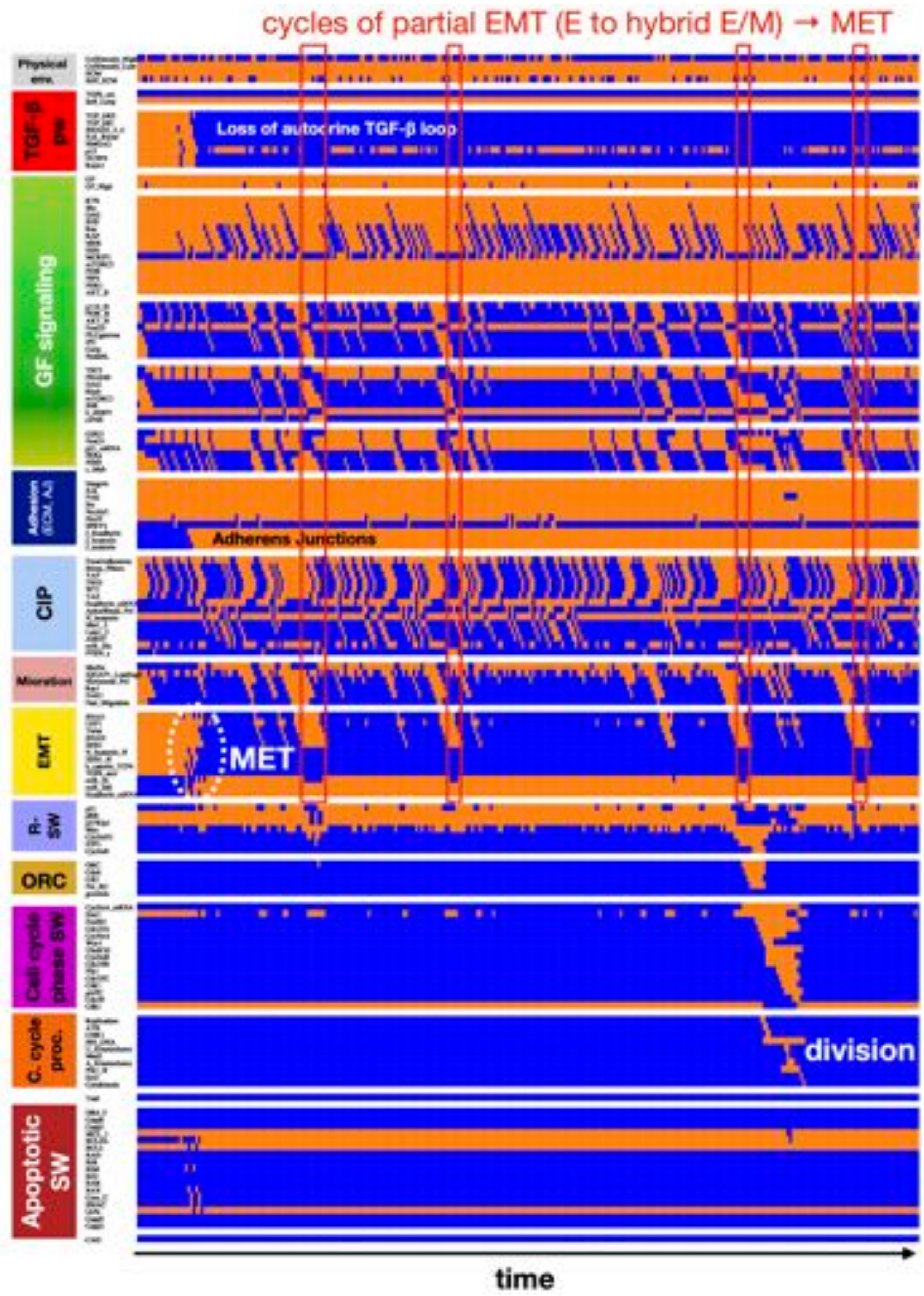


**Figure S16. Transitioning from mesenchymal to an epithelial - hybrid E/M mix requires full MET followed by reversible partial EMT; related to Figure 5.** Dynamics of regulatory molecule expression/activity of an initially mesenchymal cell on 75% stiff ECM at 20% high cell density (cell has no room to spread 20% of the time), showing irreversible EMT (white oval) followed by occasional partial EMT (E to hybrid E/M) and its reversal (red rectangles). *X-axis:* time-steps; *y-axis:* nodes organized in regulatory modules; *orange/blue:* ON/OFF.

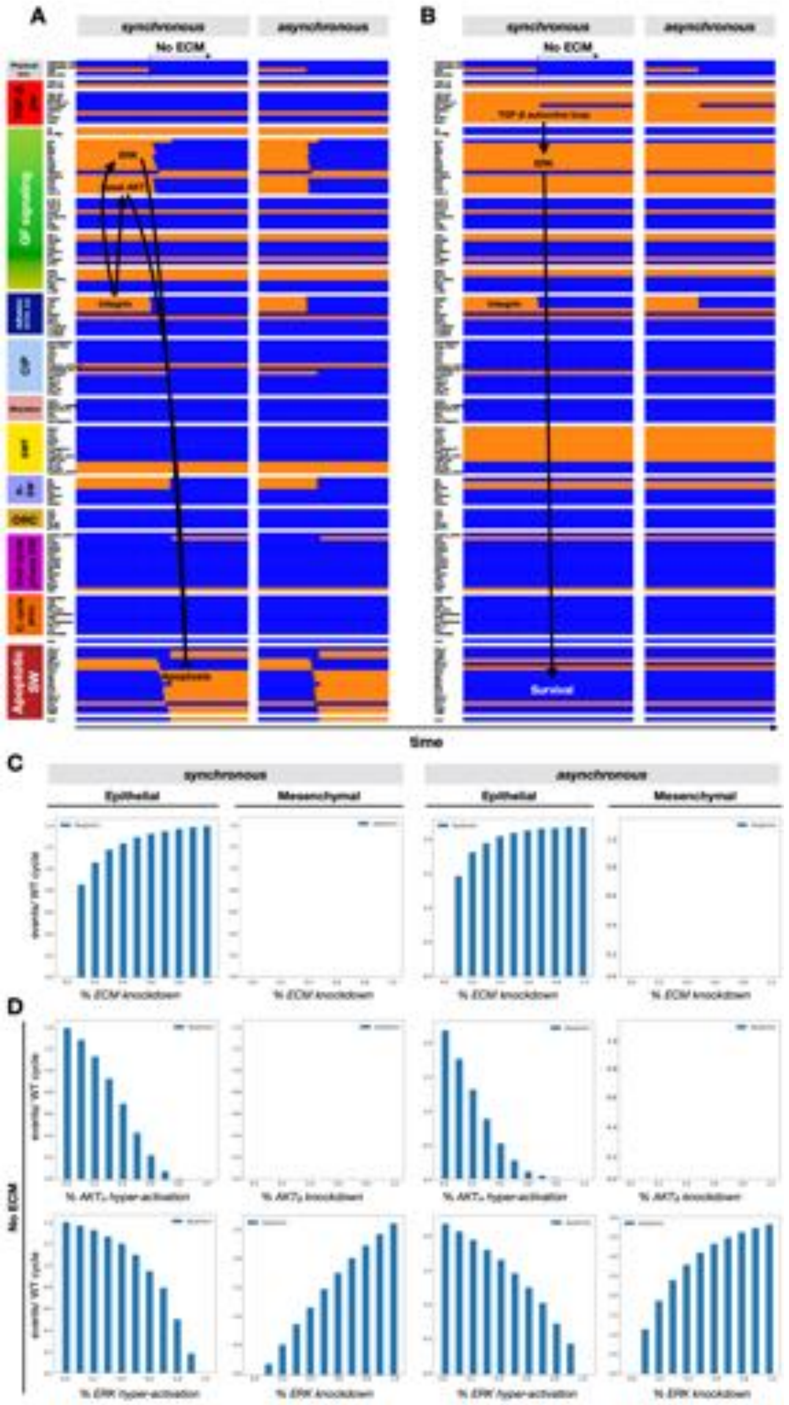


**Figure S17. Loss of anchorage to ECM leads to apoptosis in epithelial cells but *TGFβ*-mediated survival in mesenchymal cells. A-B)** Synchronous (*left*) and biased asynchronous (*right*) dynamics of regulatory molecule expression/activity in response to ECM detachment from soft ECM of (A) a growth-stimulated, isolated epithelial cell, and (B) a growth-factor starved, isolated mesenchymal cell. *X-axis:* time-steps; *y-axis:* nodes organized in regulatory modules; *orange/blue:* ON/OFF; *black/white labels & arrows:* molecular changes that drive MET. *Initial state for sampling runs:* isolated epithelial vs. mesenchymal cell in low mitogens on a soft ECM. **C)** Rate of apoptosis in isolated epithelial vs. mesenchymal cells on soft ECM in the presence of 95% saturating growth stimuli to an increasing loss of ECM adhesions. *Left/right:* synchronous/biased asynchronous update. **D)** *Left two panels:* rate of apoptosis in isolated epithelial cells upon complete loss of anchorage to an ECM together with increasing, forced activation of $AKT_H$ (*top*), or *ERK* (*bottom*). *Right:* rate of apoptosis in isolated mesenchymal cells upon complete loss of anchorage to an ECM together with increasing, forced knockdown of $AKT_B$ (*top*), or *ERK* (*bottom*) in the presence of 95% saturating growth stimuli (synchronous vs. biased asynchronous update). *Initial state for sampling runs:* isolated epithelial (*left*) or mesenchymal (*right*) cell in low mitogens on a soft ECM. *Total sampled live cell time:* 100,000 steps; synchronous update; *rate*: relative to normal cell cycle length.

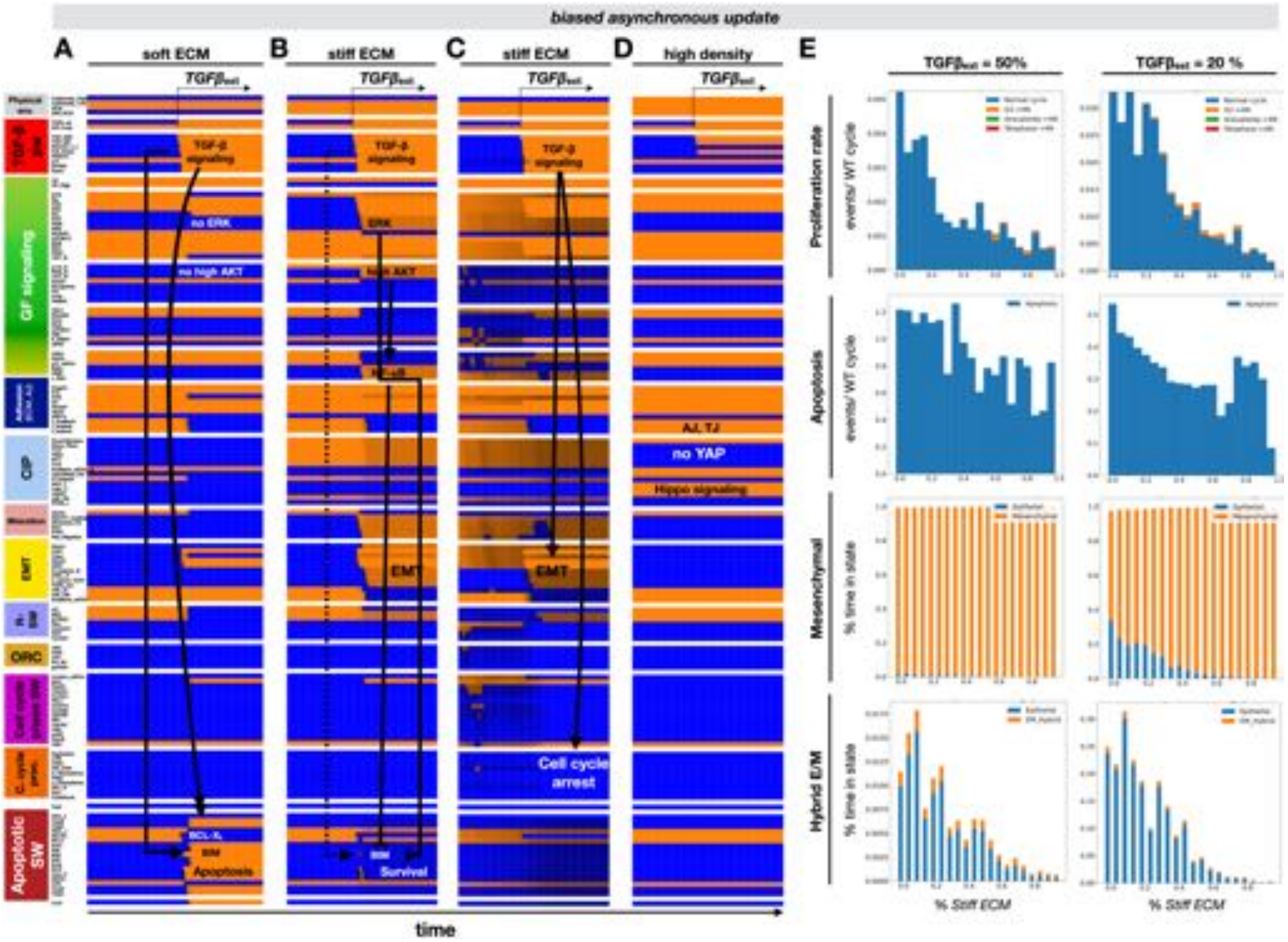


**Figure S18. Simulations reproducing Figure 6 with biased asynchronous update (ECM stiffness-dependent effects of *TGFβ*). A-D)** Biased asynchronous dynamics of regulatory molecule expression/activity during exposure to *TGFβ* of (A) growth-stimulated epithelial cells on soft ECM (medium density), (B) epithelial cells in low growth conditions on stiff ECM (medium density), (C) cycling hybrid E/M cells on stiff ECM (medium density), and (D) growth-stimulated but fully confluent (dense) epithelium on stiff ECM. *X-axis:* time-steps; *y-axis:* nodes organized in regulatory modules; *orange/black/blue color-scale:* average expression of each molecule across 1000 independent runs with biased asynchronous update (*orange* = all ON; *black* = 50% ON/OFF; *blue* = all OFF); *black/white labels & arrows:* molecular changes that drive apoptosis (A), EMT (B-C), or lack of response (D). **E)** Biased asynchronous response growth-stimulated epithelial cells at the edge of a monolayer to 50% (*left*) or 20% (*right*) external *TGFβ* at increasing ECM stiffness. *Top:* rate of normal cell cycle completion (*blue*) vs. G2 → G1 reset (*orange*), aberrant mitosis (*green*), or failed cytokinesis (*red*) followed by genome duplication, relative to the minimum cell cycle length (21 time-steps), shown as stacked bar charts. *Second from top:* rate of apoptosis (per minimum cell cycle length). *Third from top:* fraction of time spent in a mesenchymal (*orange*) vs. epithelial (*blue*) state. *Bottom:* fraction of time spent in a hybrid E/M (*orange*) vs. epithelial (*blue*) state. *Total sampled live cell time:* 100,000 steps; synchronous update. *Initial state for sampling runs:* dividing hybrid E/M at saturating mitogen exposure, at the edge of a monolayer on stiff ECM.

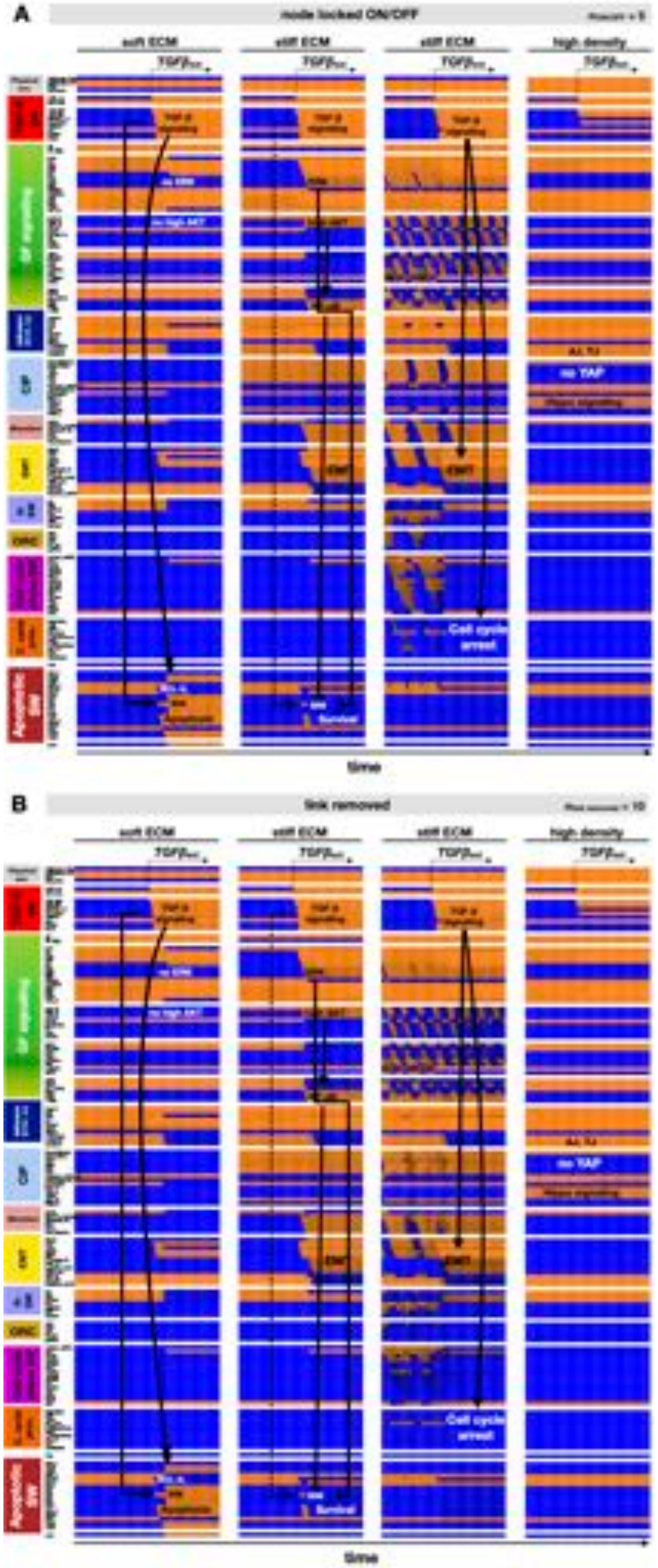


**Figure S19. The model's mechanosensitive *TGFβ* response is robust to random mutations / errors in model construction; related to Figure 6. A-B)** Average synchronous dynamics of regulatory molecule expression/activity during reversible exposure of an isolated, growth-stimulated epithelial cell to stiff ECM in an ensemble of mutant networks with (A) random nodes locked ON/OFF or (B) 10 random links removed, showing apoptosis on soft ECM (*1st panel*), EMT and cell cycle arrest on stiff ECM (*2nd and 3rd panels*), and no response at high cell density (*4th panel*). *X-axis*: time-steps; *y-axis:* nodes organized in regulatory modules; *orange/black/blue color-scale:* average expression of each molecule across 1000 time-courses from independently generated mutant models (*orange* = all ON; *black* = 50% ON/OFF; *blue* = all OFF; synchronous update); *black/white labels*: relevant molecular patterns.

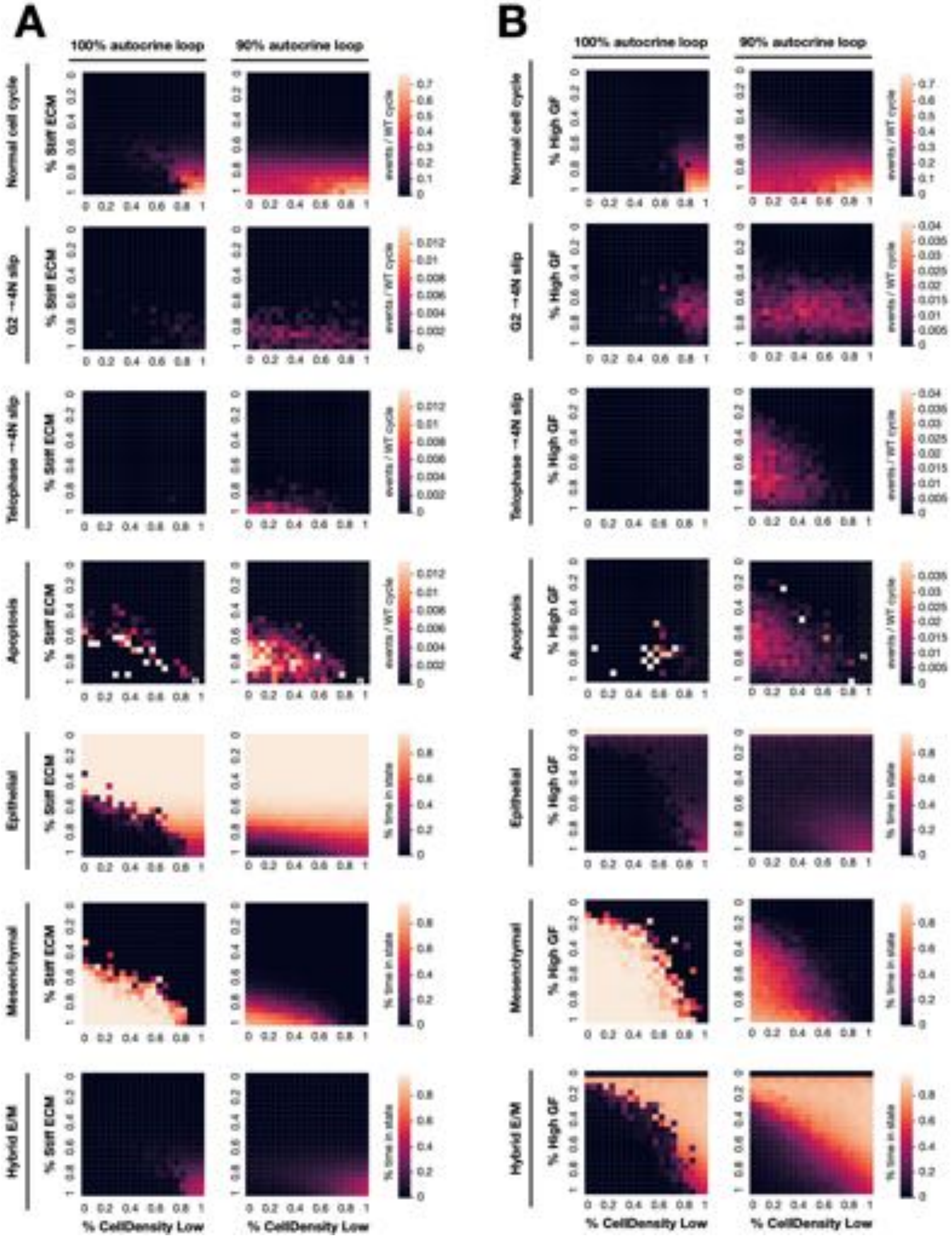


**Figure S20. Below-saturating autocrine *TGFβ* signaling destabilizes the mesenchymal state and promotes proliferation; related to Figure 6. A)** Behavior of 95% growth-stimulated epithelial cells at varying levels of density (*x* axis, from isolated to monolayer edge) and ECM stiffness (*y* axis) with 100% (*left*) or 90% (*right*) saturating autocrine *TGFβ* signaling capability (synchronous update). **B)** Behavior of epithelial cells on 95% stiff ECM at varying levels of density (*x* axis, from isolated to monolayer edge) and growth factor stimulus (*y* axis) with 100% (*left*) or 90% (*right*) saturating autocrine *TGFβ* signaling capability. *Top 4 panels:* normal cell cycle completion, G2 → G1 reset, failed cytokinesis followed by genome duplication, apoptosis (relative to minimum cell cycle length of 21 time-steps); *bottom 3 panels:* fraction of time spent in the epithelial, mesenchymal, hybrid E/M state. *Total sampled live cell time:* 100,000 steps; synchronous update; *initial state for sampling runs:* isolated epithelial cell in low mitogens on a soft ECM.
