## Supplementary material for "Boolean modeling of mechanosensitive Epithelial to Mesenchymal Transition and its reversal": Files S2-S10 (models in different formats): File_S9_EMT_Mechanosensing_Steady_state_MAP.pdf

56 attractors for all Trail = 0 environments
(an additional 26 apoptotic attractors exist in Trail = 1 environments)

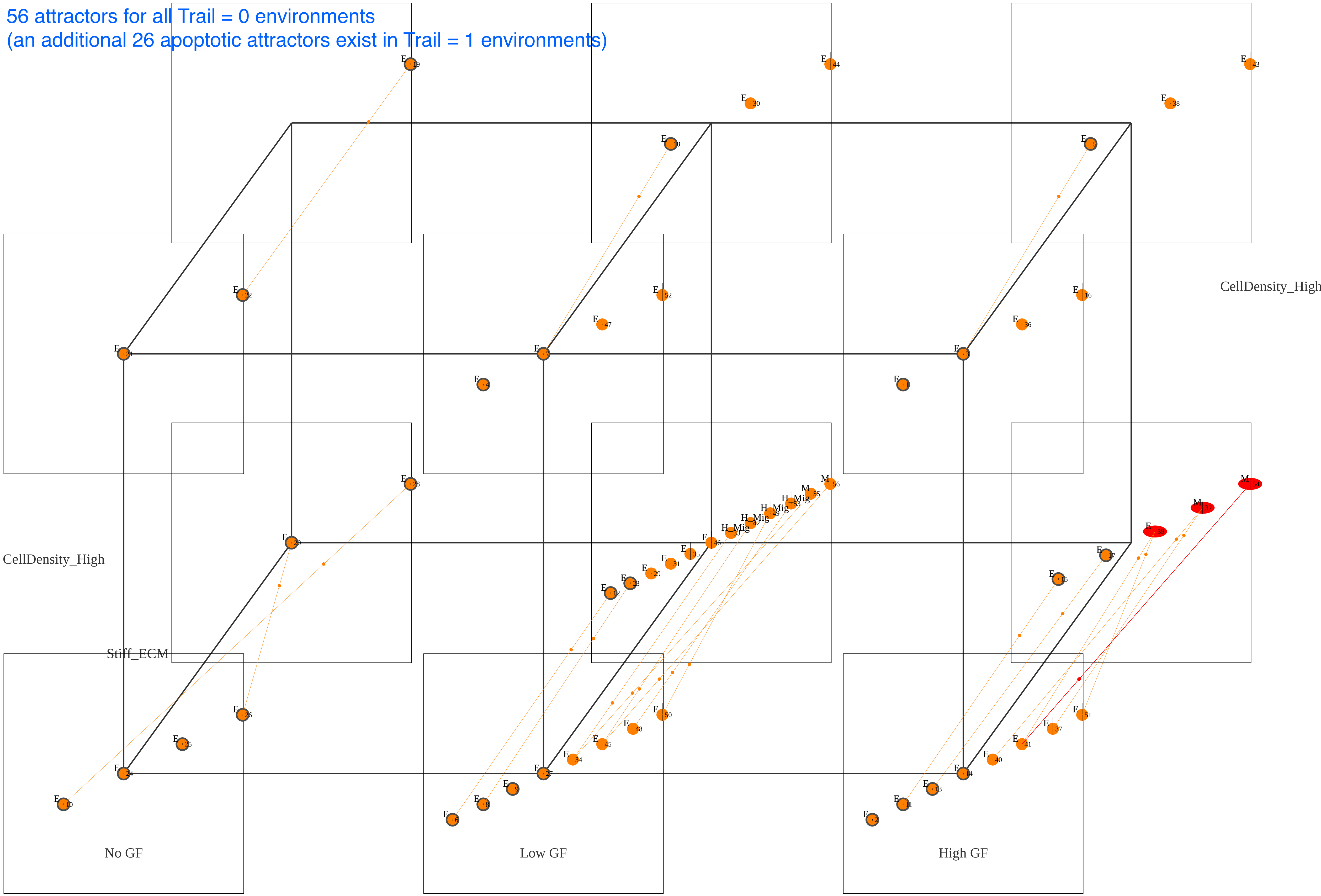
